# Reconstruction of single cell lineage trajectories and identification of diversity in fates during the epithelial-to-mesenchymal transition

**DOI:** 10.1101/2023.09.19.558325

**Authors:** Yu-Chen Cheng, Yun Zhang, Shubham Tripathi, BV Harshavardhan, Mohit Kumar Jolly, Geoffrey Schiebinger, Herbert Levine, Thomas O. McDonald, Franziska Michor

**Affiliations:** Department of Data Science, Dana-Farber Cancer Institute, Boston, MA, USA; Department of Biostatistics, Harvard T.H. Chan School of Public Health, Boston, MA, USA; Center for Cancer Evolution, Dana-Farber Cancer Institute, Boston, MA, USA; Department of Stem Cell and Regenerative Biology, Harvard University, Cambridge, MA, USA; State Key Laboratory of Molecular Oncology, National Cancer Center/National Clinical Research Center for Cancer/Cancer Hospital, Chinese Academy of Medical Sciences and Peking Union Medical College, Beijing, 100021, China; Department of Immunology, Yale University, New Haven, CT, USA; Interdisciplinary Mathematics Initiative, Indian Institute of Science, Bangalore, 560012, India; Centre for BioSystems Science and Engineering, Indian Institute of Science, Bangalore, 560012, India; Department of Mathematics, University of British Columbia, Vancouver, BC, Canada; Center for Theoretical Biological Physics, Northeastern University, Boston, MA, USA; Department of Physics, Northeastern University, Boston, MA, USA; The Broad Institute of MIT and Harvard, Cambridge, MA, USA; The Ludwig Center at Harvard, Boston, MA, USA

**Keywords:** Epithelial-mesenchymal transition, Optimal transport, Trajectory inference

## Abstract

Exploring the complexity of the epithelial-to-mesenchymal transition (EMT) unveils a diversity of potential cell fates; however, the exact timing and intricate mechanisms by which early cell states diverge into distinct EMT trajectories remain unclear. Studying these EMT trajectories through single cell RNA sequencing is challenging due to the necessity of sacrificing cells for each measurement. In this study, we employed optimal-transport (OT) analysis to reconstruct the past trajectories of different cell fates during TGF-beta-induced EMT in the MCF10A cell line. Our analysis revealed three distinct trajectories leading to low EMT, partial EMT, and high EMT states. Cells along partial EMT trajectory showed substantial variations in the EMT signature and exhibited pronounced stemness. Throughout this EMT trajectory, we observed a consistent downregulation of the *EED* and *EZH2* genes. This finding was validated by recent inhibitor screens of EMT regulators and CRISPR screen studies. Moreover, we applied our analysis of early-phase differential gene expression to gene sets associated with stemness and proliferation, pinpointing *ITGB4*, *LAMA3*, and *LAMB3* as genes differentially expressed in the initial stages of the partial versus high EMT trajectories. We also found that *CENPF*, *CKS1B*, and *MKI67* showed significant upregulation in the high EMT trajectory. While the first group of genes aligns with findings from previous studies, our work uniquely pinpoints the precise timing of these upregulations. Finally, the latter group of genes represents newly identified regulators, shedding light on potential targets for modulating EMT trajectories.

**Significance Statement:** In our study, we investigated cellular trajectories during EMT using a time-series scRNAseq dataset. OT analysis was used to infer cell-to-cell connections from scRNAseq data, allowing us to predict cell linkages and overcome limitations of sequencing such as the need to sacrifice cells for each measurement. This approach allowed us to identify diverse EMT responses under uniform treatment, a significant advancement over previous studies limited by the static nature of scRNAseq data. Our analysis identified a broad set of genes involved in the EMT process, uncovering novel insights such as the upregulation of cell cycle genes in cells predisposed to a high EMT state and the enhancement of cell adhesion marker genes in cells veering towards a partial EMT state. This work enriches our understanding of the dynamic processes of EMT, showcasing the varied cellular fates within the same experimental setup.

## Introduction

The epithelial-mesenchymal transition (EMT) is a pivotal process underpinning a range of biological phenomena from embryonic development and wound healing to tumor metastasis^1–5^. During EMT, epithelial cells lose their apical-basal polarity and adhesion to other cells and acquire mesenchymal traits such as invasiveness and migratory capabilities^3–5^. At the molecular level, this process is accompanied by the downregulation of epithelial markers such as E-cadherin (*CDH1*) and a concurrent upregulation of mesenchymal markers like N-cadherin (*CDH2*), vimentin (*VIM*), and fibronectin (*FN*)^6,7^ . Importantly, EMT is not merely a binary transition from an epithelial (E) to a mesenchymal (M) state. Recent findings redefine EMT as a continuum, with cells capable of occupying intermediate states, often referred to as ’partial’ EMT^8,9^. Progression along this spectrum is tightly regulated by a set of key transcription factors, including members of the Snail, Zeb, and Twist families^10,11^. The expression and activities of these transcriptional factors are governed by a complex network of several epigenetic regulators and signaling pathways, encompassing TGF-beta, Wnt, EGF, FGF, PI3K/Akt/mTOR, IL6-JAK-STA3, and NOTCH^5,12–16^.

Cells in a syngeneic, phenotypically homogeneous population have been observed to adopt distinct fates upon treatment with an EMT inducer^17,18^. However, the intricate mechanisms that drive early cell states to branch into unique EMT trajectories are yet to be fully understood. The idea of divergent trajectories, through a developmental Waddington landscape^19^, is well-accepted in stem cell biology^20^. Given the close association between EMT and stemness^21,22^, we aimed to investigate whether the heterogeneous response to EMT inducers extends beyond mere temporal variations and involves multiple distinct trajectories. To this end, we analyzed previously published time series scRNAseq data from MCF10A cells treated with TGF-beta^18^ (Fig 1A).

**Figure 1.**
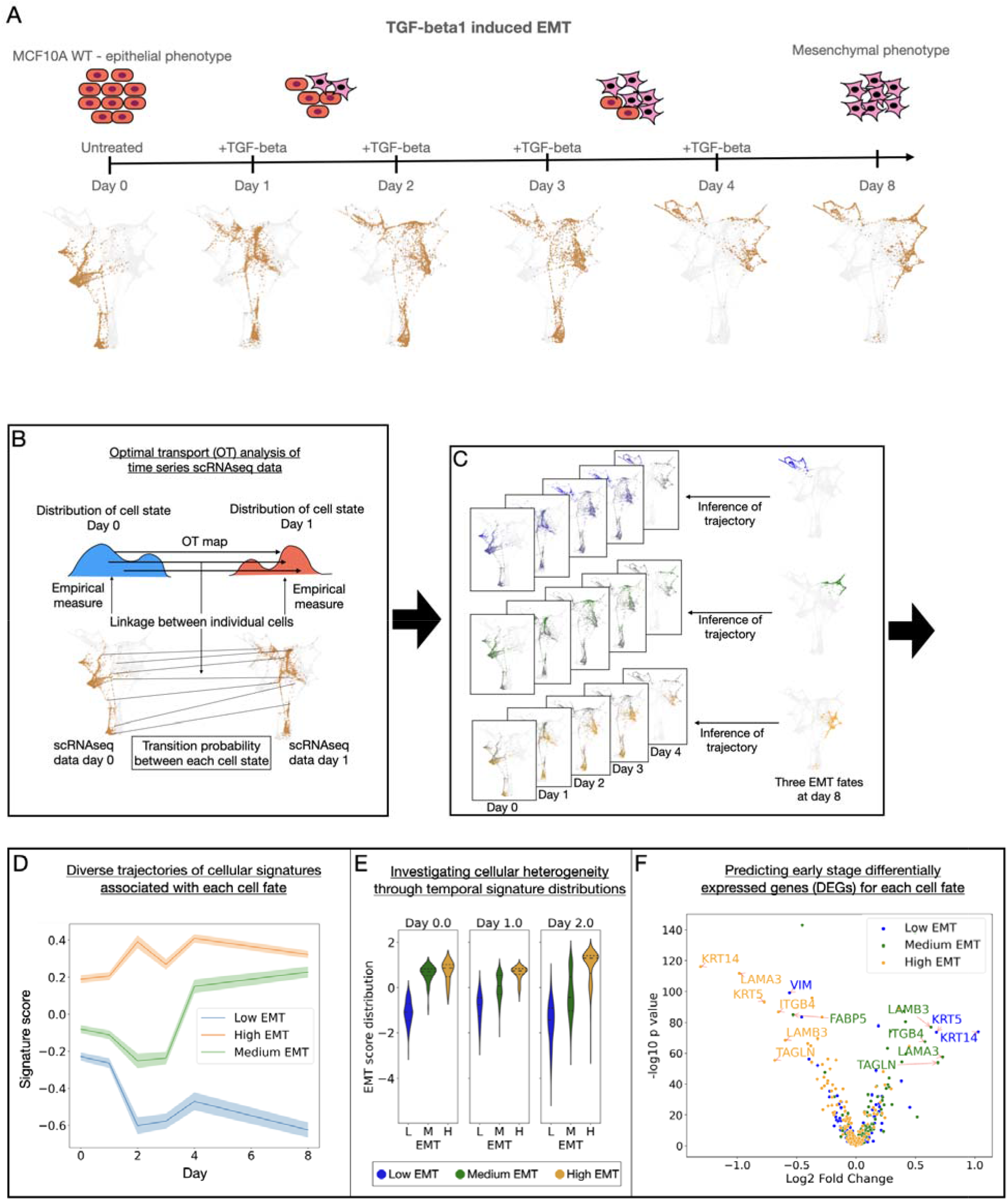
Optimal transport analysis of scRNAseq time course data of the epithelial mesenchymal transition. (A) Schematic representation of the experiments performed to induce EMT in MCF10A cells, accompanied by a time series of single-cell RNA sequencing data visualized using force-directed layout embedding (FLE). The total cell count amounts to 12,588, encompassing all cells shown as grey and brown dots. The daily distribution is as follows: 2,734 cells on day 0, 2,303 on day 1, 2,381 on day 2, 2,132 on day 3, 1,147 on day 4, and 1,891 on day 8, with each day’s count represented by brown-colored dots. This figure was adapted from Deshmukh *et al*^17^ (B-F) General framework of optimal-transport analysis of EMT single-cell RNA sequencing data: (B) OT analysis was employed to identify the transition probability of cell-to-cell connections between adjacent time points within the scRNAseq datasets. (C) From the entire consecutive time-series scRNAseq data, transition probabilities, inferred by OT, were integrated to determine the likelihood of each early cell state acting as an ancestor for the three fate subpopulations. (D-F) Three downstream trajectory inference analyses are depicted from left to right: (D) reconstruction of diverse cellular signature trajectories, (E) exploration of cellular heterogeneity across these diverse trajectories, and (F) differential analysis of early gene expression in ancestral cells associated with each distinct cell fate. In this instance, we utilized a set of 480 genes identified as a stemness signature from Lim *et al*^55^.

While scRNAseq data offers a wealth of insights into the heterogeneity of cellular states^23,24^, the inherent need to sacrifice cells at each time point precludes the ability to trace individual cell lineages over time. This restriction poses a challenge to reconstructing trajectories from time-series scRNAseq data^25^. To address this challenge, we employed a method based on OT analysis^26,27^, known as Waddington OT (WOT)^28^. This method stands in contrast to other widely-used trajectory tools such as pseudotime analysis, which infers a temporal sequence within a cell population but cannot deduce direct cell-to-cell transitions ^29,30^. Another method, RNA velocity, utilizes additional information from unspliced and spliced RNA to predict the direction of movement across RNAseq space of individual cells ^31,32^. This method deepens our insight into the velocity field and short-term cellular changes. However, its application to long-term cell-to-cell transition reconstruction is indirect, leading to imprecise and noisy inferences^33^. In contrast, WOT is specifically tailored for analyzing direct cell-to-cell transitions within scRNAseq data collected at discrete, predetermined time points, distinguishing it from these other methodologies.

Utilizing the WOT technique, we reconstructed lineage trajectories at single-cell resolution using the time series scRNAseq data from MCF10A cells undergoing EMT stimulated by TGF-beta^18^, enabling identification of diverse trajectories leading to distinct EMT fates. In this study, we extend previous EMT research by not only examining state heterogeneity at various time points within a single EMT process but also by uncovering the diversity of EMT responses as unique, distinct processes under the same treatment. We delved into the roles of stemness, proliferation, and cellular hypoxic response signatures. While these signatures have known associations with EMT^5,21,34^, their variations across different EMT trajectories have not been extensively explored.

Furthermore, our trajectory analysis at the single gene level enabled us to predict differentially expressed genes (DEGs) in the earliest phases of each fate. Early gene expression linked to a specific fate was then partially validated through methods such as inhibitor screens of EMT regulators and CRISPR-associated gene knockout screens^35^, highlighting the robustness of our predictions. We then broadened our method’s application to include a wider array of genes implicated in EMT regulation but not yet fully examined. This approach led to several novel insights, notably that cell cycle-related genes are upregulated in the ancestors of cells entering the high EMT state. Additionally, we found that genes linked to cell surface markers that play a critical role in cell-matrix and cell-cell adhesion are markedly upregulated in the ancestors of cells transitioning to the partial EMT state. An overview of the general framework is provided in Fig 1B-F.

## Results

### Uncovering Three Distinct EMT Trajectories via Optimal Transport Analysis

Given that scRNAseq data cannot be obtained from cells obtained during multiple time points of lineage tracing experiments, we set out to computationally infer likely ancestor cell states for different EMT fates. In the study by Deshmukh *et al.*^18^, an immortalized human mammary epithelial cell line, MCF10A, was treated with TGF-beta for 1, 2, 3, 4, or 8 days (Fig 1A) and scRNAseq data was obtained from populations sacrificed at each time point. Through cluster analysis of the scRNAseq data at day 8, we identified three subpopulations representing three significantly different cell fates (Methods). These fates were categorized as low, medium, and high EMT by utilizing the 76GS and KS scoring metrics to compute the average EMT scores^36–39^, for each subpopulation (Fig 2A, day 8). For instance, using the 76GS method, we derived average EMT scores of -0.63, 0.23, and 0.32 for the low, medium, and high EMT categories, respectively, with significant p-values (*t*-test, p < 0.005) for each pairwise comparison (Fig 3B and Table S4, day 8). Use of the KS method yielded consistent outcomes (Fig S5).

**Figure 2.**
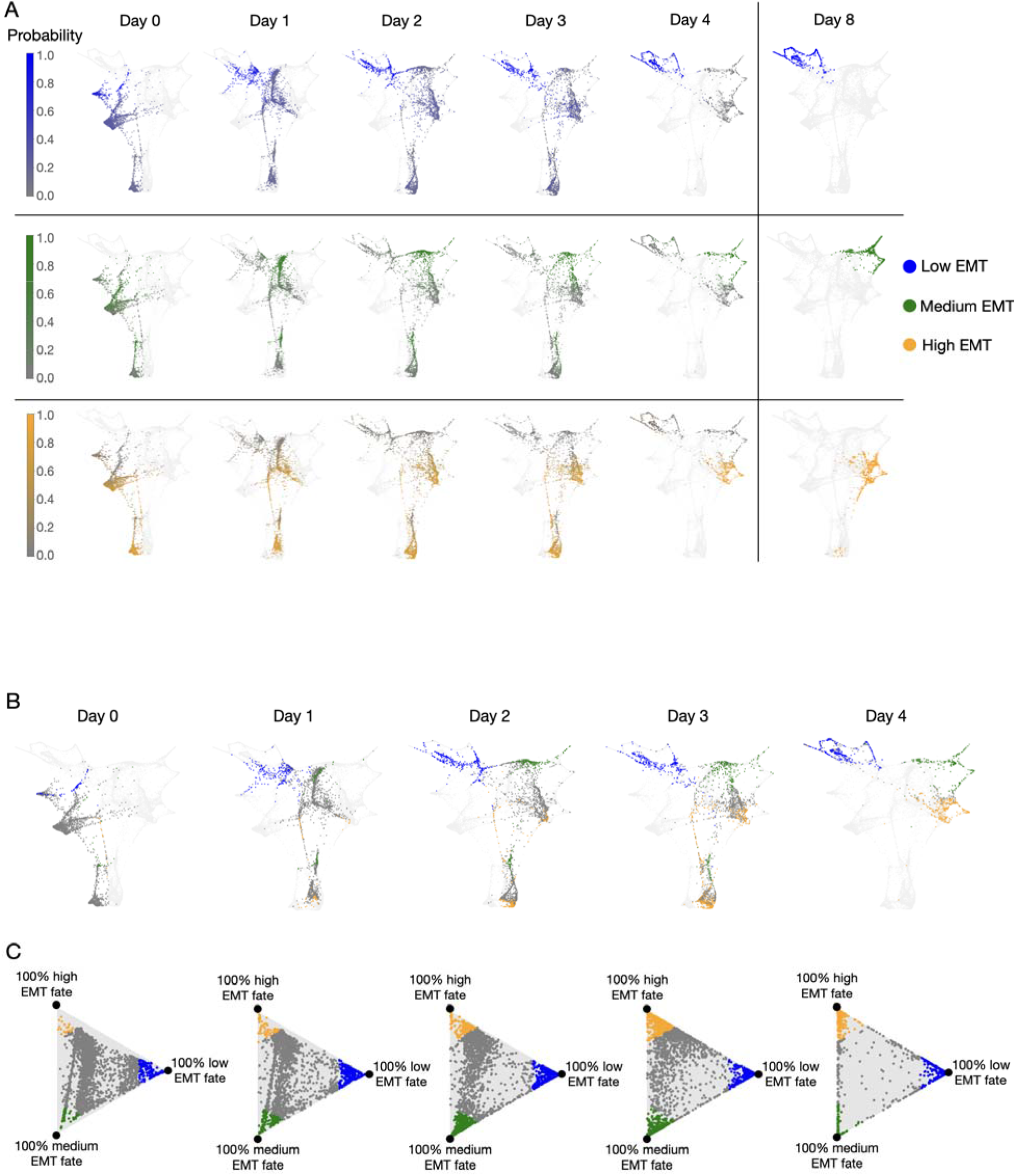
Optimal transport recovers diverse trajectories of EMT. (A) The colormap presents the inferred ATF distributions, showcasing the probability of early cell states (from day 0 to day 4) serving as ancestors for the three fate subpopulations by day 8. (B) Early cell state classification via ATF distributions: blue cells have over 75% probability of becoming cells with the low EMT fate; green cells have over 75% probability of becoming the medium EMT fate; orange cells have over 75% probability of becoming the high EMT fate; and gray cells are not highly committed to any of the fates. (C) Barycentric coordinate projection visualizes ATF distributions: this coordinate system represents the three-dimensional simplex within a two-dimensional space. For each time point, every individual cell is associated with a three-dimensional probability vector, as determined by that specific time point’s ATF distributions of the three fates (each column in A). This vector is then mapped onto an equilateral triangle (Methods). A position at one of the triangle’s vertices indicates a 100% commitment of the cell state to the corresponding fate.

**Figure 3.**
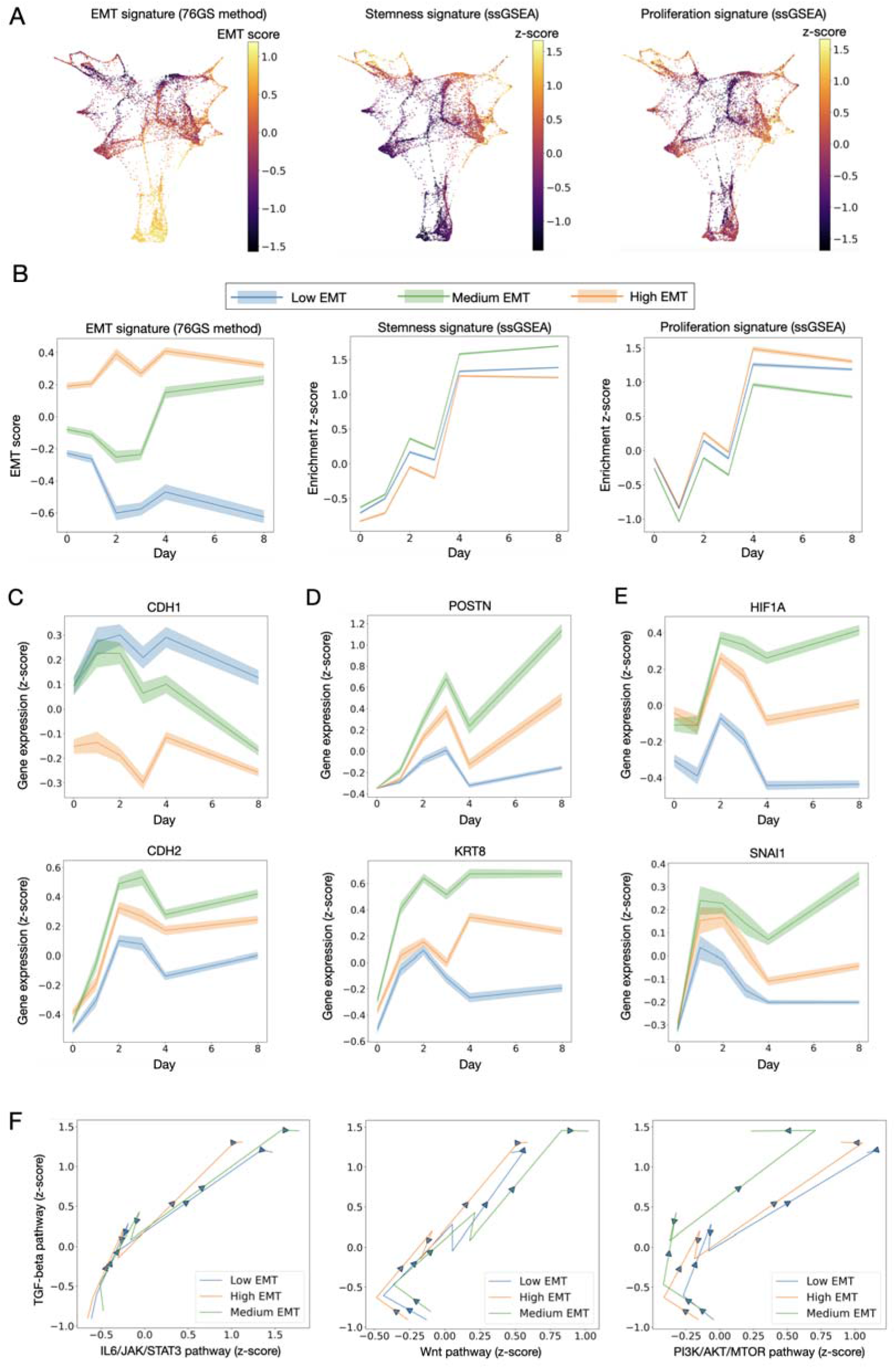
OT analysis reveals unique cellular signatures across distinct EMT trajectories. (A) Color maps illustrate the EMT signature score (using the 76GS method), stemness signature score (via ssGSEA), and proliferation signature score (via ssGSEA) for all cellular states gathered from day 0 to day 8. (B) The panels depict the time progression of average cellular signature scores (left to right: EMT, stemness, and proliferation) across the three distinct EMT trajectories. The p-values, generated via paired-sample *t*-tests for each pairwise comparison at individual time points, are all < 0.005. Shaded regions denote the 95% confidence intervals. (C-E) Temporal evolution of mean gene expression, from day 0 to day 8, across the three EMT trajectories. Shaded regions denote the 95% confidence intervals (C) for *CDH1* and *CDH2* genes, (D) for *HIF1A* and *SNAI1* genes, and (E) for *POSTN* and *FN8* genes. (F) Two-dimensional plots illustrate the time-course progression of average cellular signature scores for paired signaling pathways. Lines connect daily average scores for each signature pair, with arrows highlighting the directional flow of time.

To infer the trajectories of individual cell states across the sequential scRNAseq dataset, we utilized WOT^28^. The dataset consists of six distinct batches, each sourced from a uniformly mixed single culture of around 10,000 cells. This setup provides a consistent starting point for each batch before the application of TGF-beta, allowing us to assume uniform initial conditions across the batches. Leveraging this baseline, the WOT method predicts a unique transition probability (i.e., the likelihood that one cellular state is the ancestor and the other the descendant) between two adjacent scRNAseq time points. This prediction assumes that cellular states navigate the gene expression space using the shortest overall distance (Fig S1 and Methods). By multiplying the inferred transition probabilities from initial to subsequent time points within our scRNAseq data series, we computed the probability of each early cell state, termed ‘ancestors’, transitioning into a final cell state at day 8 or ‘fate’ (Fig 2A). We refer to these transition probabilities as ’ancestor-to-fate (ATF) distributions’. To validate our inferred distributions, we followed an approach of omitting data of a specific time point, designated as test data, and comparing our estimated cell state distribution to the actual data of this time point. The results showed minimal deviations between predictions and actual data, confirming our predictions’ reliability when contrasted with other intrinsic cellular variations and unbiased interpolations (Methods).

To identify cellular origins leading to various fates, we categorized cells with over 75% probability of transitioning to specific fates as ’top ancestors’ (Methods). Notably, prior to treatment, the percentage of top ancestors for the low EMT fate constituted double the combined percentage of the other two fates (5.52% vs. 2.85% at day 0). By the second day of treatment, the proportions of top ancestors across all three fates converged, with values of 13.57%, 12.35%, and 13.31% for low, medium, and high EMT (Fig 2BC). This temporal shift in proportions indicates a delayed inclination towards the medium and high EMT fates, induced by TGF-beta. Additionally, cells falling below the probability threshold for any EMT fate were classified as ‘undetermined ancestors’. With ongoing TGF-beta treatment, the portion of the undetermined ancestors decreased sharply from 91.62% on day 0 to 15.92% on day 4. This trend may suggest that initially, a high percentage of undetermined ancestors indicates a high level of cell plasticity before the treatment; however, following the treatment, this plasticity might reduce as more cells advance towards predetermined fates. Consistently, these interpretations are supported across alternative probability thresholds for defining top ancestors, ranging from 75% to 90% (Methods and Supplementary Appendix S1).

Upon inspection of the full trajectories, we observed that the ancestors of the three fates were dispersed without clear boundaries, unlike the three distinct, well-outlined regions seen at day 8 for the three fates (Fig 2A and Fig S2). This observation, combined with the profound reduction in the percentage of undetermined ancestors post-treatment (75% decrease, Fig 2C) suggests that over time, cells exhibit decreased plasticity and increasingly tend toward more determined states. This trend indicates a divergence in EMT phenotypes. To quantify this divergence, we computed the Wasserstein 2-distance^40^ between the cell state distributions of each pair of trajectories at every time point (Methods). Our analysis revealed a marked divergence between every pair among the three trajectories: by day 8, the distance had increased 2.67 times from its day 0 measurement for both the low vs. medium and low vs. high EMT trajectories, and 2.17 times for the high vs. medium EMT trajectory. Notably, this divergence was most pronounced before day 4, accounting for 90% of the total increase. The divergence then leveled off, with only a 10% increase observed afterward, indicating that the divergence between trajectories increases most significantly during early stages of TGF-beta treatment (Fig S3B).

### Deciphering Unique Gene Signatures Across Trajectories of Distinct EMT States

To trace the EMT characteristics of the three fate subpopulations to their origins, we first determined the EMT score for each cellular state, from day 0 to day 8, using the 76GS and KS EMT scoring methods (Fig 3A, Fig S5, and Methods). For each time point, we integrated the EMT scores across all cell states, each weighted by their likelihood of being the ancestor for a particular fate subpopulation as determined by the ATF distributions (Fig S4 and Methods). This approach unveiled three distinct trajectories, each showing unique average EMT score trajectories with non-overlapping 95% confidence intervals, throughout the course of TGF-beta treatment (Fig 3B and Fig S5). The clear separation into low, medium, and high EMT trajectories was consistently observed using both the 76GS and KS EMT scoring methods (Fig S5 and Methods). Notably, the separation of trajectories was observed even before the initiation of TGF-beta treatment on day 0, implying that early EMT hallmarks could predestine cellular EMT fates (Fig. 3B). However, these hallmark trajectories only reflect the mean EMT scores for cells on a specific path, and a subset of early state cells—undetermined ancestors—remain (Fig. 2C). In the following section, we expand our analysis to include the full distribution beyond average EMT scores. This approach specifies the EMT hallmarks in cells with predetermined fates (top ancestors)—as well as in those exhibiting intrinsic plasticity (undetermined ancestors).

Furthermore, we analyzed stemness, proliferation signatures, and associated hallmarks of hypoxia response and the G2M checkpoint among cells belonging to the three fates. For each cell, we computed those signatures using single-sample gene set enrichment analysis (ssGSEA, Fig 3A and Methods). All trajectories showed an over 1.9 z-score increase in both stemness and hypoxia response, following a similar pattern across these signatures (Fig 3B and Fig S5). This trend aligns with prior research that links hypoxia to enhanced stemness in EMT^41,42^. Like the EMT signature, these three trajectories stood out with their non-overlapping 95% confidence intervals when characterized by these two signatures (Fig 3B and Fig S5). Of particular interest was that by day 8, the medium EMT trajectory exhibited the highest levels of stemness and hypoxia response, with enrichment scores of 1.7 for both. In comparison, the low and high EMT trajectories displayed scores of 1.4 and 1.2, respectively (Fig 3B and Fig S5). These findings resonate with earlier studies identifying an intermediate EMT stage characterized by heightened stemness and a pronounced response to hypoxia^4,34,43^.

Upon analyzing the proliferation signature trajectories, we noted enrichment z-score declines from day 0 to day 1 (low EMT: -0.12 to -0.84, medium EMT: -0.26 to -1.04, high EMT: -0.11 to - 0.8, Fig 3B). These z-score shifts indicate notable decreases relative to the average enrichment score. A similar trend was observed in the G2M checkpoint hallmark (Fig S5). This decrease reflects the known role of TGF-beta in inhibiting cell division^44–46^. From day 1 through day 8, cells regain their proliferative capacity, evidenced by enrichment score of proliferation rebounds of 2.0 for low, 1.8 for medium, and 2.1 for high EMT (Fig 3B and Fig S5). Based on these changes, we concluded that the medium EMT trajectory was distinctive, exhibiting the most pronounced decline and the least recovery in proliferation signatures. This unique trend in the medium EMT cells corresponds with their pronounced response to the TGF-beta inducer, evident by TGFBI (a TGF-beta induced gene) showing more elevated expression in this trajectory than in the other two (Fig S7).

We then further analyzed the dynamics of individual genes pivotal to EMT, such as *CDH1* (E-cadherin) and *CDH2* (N-cadherin). We found that the medium EMT trajectory initially displays high *CDH1* expression that diminished toward the end of treatment, shifting from a z-score of 0.09 on day 0 to -0.17 on day 8 (Fig 3C). This downward trend aligns with previous findings indicating that *CDH1* downregulation triggers partial EMT^47^. Conversely, *CDH2* expression notably increased in the medium EMT trajectory, diverging from the patterns seen in the high and low EMT trajectories (Fig 3C). Furthermore, these gene expression patterns are markedly distinct, underscored by their non-overlapping 95% confidence intervals. This finding aligns with a previous study showing elevated expression of *CDH2* in partial EMT using the same cell line and treatment type^35^. Beyond *CDH2*, Zhang *et al.*^35^ highlighted elevated expression of *POSTN* and *KRT8* expressions as indicators of the partial EMT phase. In our study, the medium EMT trajectory mirrored this finding, with *POSTN* and *KRT8* expression levels surpassing those in the high and low trajectories (Fig 3D). Additionally, we detected a pronounced rise in *HIF-1A* and *Snail* expression within the medium trajectory compared to the others (Fig 3E). This finding further supports the classification of the medium EMT trajectory as partial EMT, given the known roles of these genes in hypoxia and partial EMT fates^34,48^.

To investigate whether TGF-beta treatment correlates with other essential EMT-related signaling pathways, we further conducted pairwise comparisons of various cellular signatures over time (Methods). Across all trajectories, we found positive correlations between TGF-beta signaling and the IL6-JAK-STAT3, Wnt, and PI3K-AKT-mTOR pathways, with Pearson correlation coefficients ranging across trajectories from 0.95 to 0.97, 0.92 to 0.96, and 0.76 to 0.92, respectively (Fig 3F). These observations are consistent with previous findings regarding the concurrent regulation of these pathways throughout the EMT process^9,49,50^. Particularly, the intricate interplay between TGF-beta and PI3K signaling pathways, which includes both antagonistic and cooperative interactions, has been discussed previously^9^. In our study, while the TGF-beta pathway activity increased from day 0 to day 8 across all three EMT trajectories, the PI3K pathway interestingly showed a decline in the partial EMT trajectory by the treatment’s end. In contrast, the enrichment scores for the other two trajectories remained relatively stable (Fig 3F). With PI3K signaling recognized as a prominent driver of cell growth and proliferation^51^, this observed decline aligns with the lower proliferation scores and G2M checkpoint pathway activity levels noted along the partial EMT trajectory (Fig. 3B and Fig. S5).

### Unveiling Increased EMT Heterogeneity within the Partial EMT Trajectory

To deepen our understanding of cellular heterogeneity across EMT trajectories, we studied the temporal evolution of EMT signature distributions along the three identified paths. We employed several methodologies to evaluate the within-trajectory distributions. First, we incorporated chronological sequences of triangle plots (Fig 2B) with time-ordered individual cellular EMT signature scores (Fig 4A and Fig S8). This integration elucidated the relationship between ancestral cell EMT states and their potential to transition into a specific fate (Methods). The triangle plots demonstrate that the top ancestors, showing over 75% commitment to the high/low EMT fate, consistently exhibited high/low EMT signatures during the initial stages of the treatment process. Conversely, for the top ancestors of the partial EMT fate, EMT scores were notably heterogeneous, encompassing the full spectrum from low to high EMT cell types (Fig 4B and Fig S8). This pattern is discernible throughout day 3 (Fig 4B and Fig S8), suggesting that this early phase of the partial EMT trajectory displays a greater degree of variability in EMT expression scores compared to the early phases of the other EMT trajectories.

**Figure 4.**
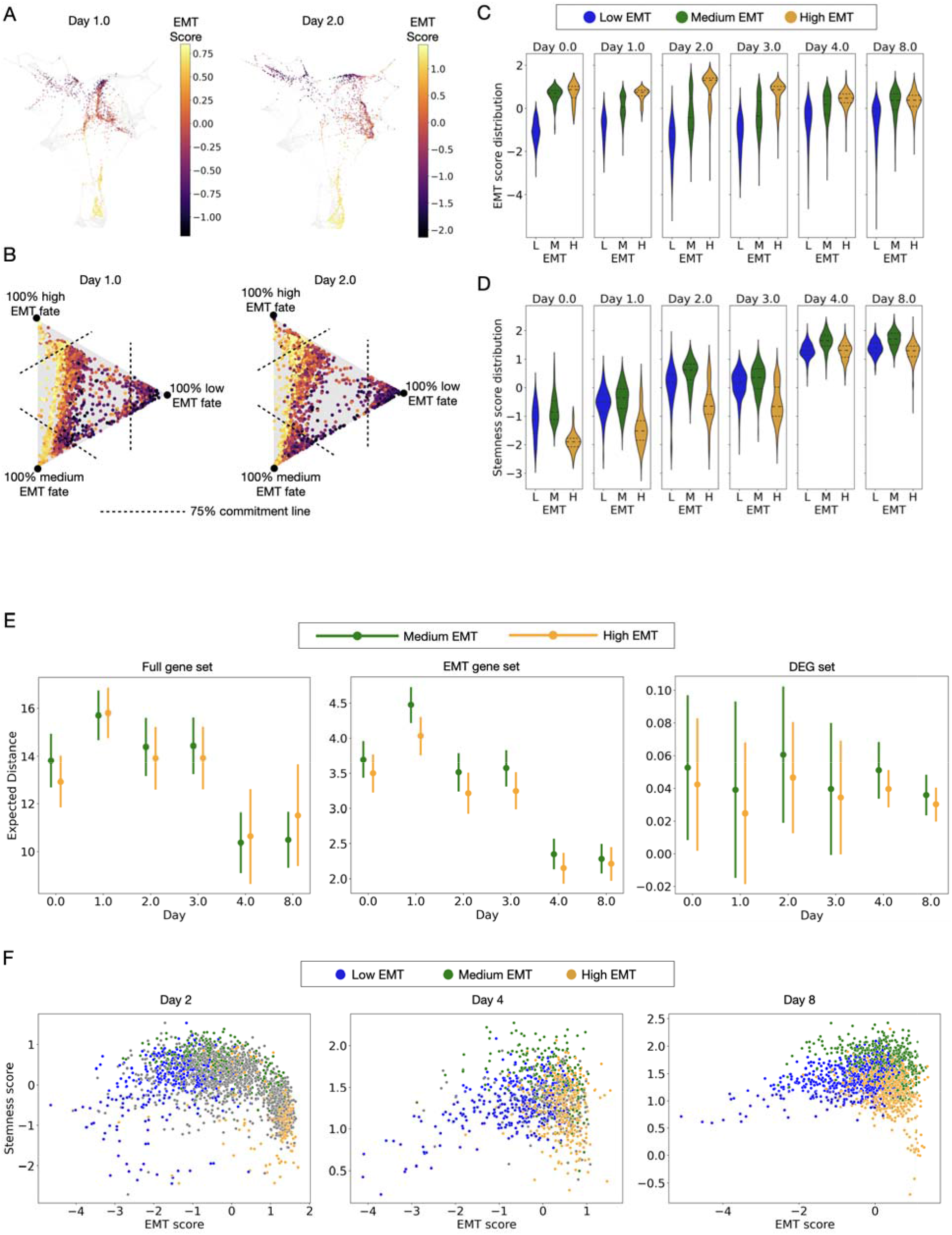
Tracing cellular signature variations across three EMT trajectories. (A) EMT signature scores for cell states from days 1 and 2 (for the complete time course see Supplementary Fig 8). (B) EMT signature scores from (A) are paired with ATF distributions and plotted within a triangle using barycentric coordinates. As in Fig 2C, a point’s location represents its ATF distribution, indicating its likelihood of differentiating into the associated fate. Concurrently, the color map showcases the EMT score. Dashed lines demarcate a 75% commitment to the fate linked to the corresponding triangle vertex. (C-D) Violin plots depict the distribution of each cellular signature score for the top ancestors of each fate, color-coded in blue for low, green for medium, and orange for high EMT trajectories (C) for EMT score (via 76GS method) and (D) for stemness score (via ssGSEA). (E) The error bar plots depict the mean of weighted pairwise distances in cellular transcriptomics (indicated at the center of each bar), and the standard deviation errors of these pairwise distances (symbolized by the length of the error bars). The low EMT trajectory was not included due to the high number of outliers, as explained in the main text and Supplementary S1. These distances are derived from the gene expression space defined by the pertinent gene sets: complete gene set, EMT gene set, and DEGs. (F) Scatter plots display paired cellular signature scores for days 2, 4, and 8. Color codes designate the top ancestors for each trajectory.

To further characterize the extent of heterogeneity within the partial EMT trajectory, we calculated pairwise cell state distances^52^ (Methods), focusing on the differences between the partial and high EMT trajectories. The low EMT trajectory was excluded due to the high number of outliers (for details see Table S5). To determine if the high variability was uniquely tied to the EMT signature, we computed cell state distances across three gene expression spaces: the full gene set, the EMT signature gene set, and genes differentially expressed between the partial and high EMT fates (Table S1 and Methods). Notably, during the four days of treatment, cell state variability defined by the average pairwise cell distances in the EMT gene expression space was markedly higher in the partial EMT compared to the high EMT trajectory (*t*-test, p < 1e-9; fold changes were 1.11, 1.09, 1.10 for days 1 through 3, respectively) (Fig 4E and Table S2). However, within the same period, such differences were non-significant when cell state heterogeneity was determined using the other two gene spaces. Exceptions were only found on days 2 and 3 within the full gene set, but they demonstrated minor fold changes of 1.03 and 1.04, respectively. This specific variability of the EMT score in the partial EMT trajectory aligns with prior research suggesting a lack of association between core EMT transcription factors and the partial EMT state^53^.

Beyond pairwise cell state distances, we also explored cellular heterogeneity by analyzing EMT and stemness score distributions among the top ancestors of the three identified EMT fates (Methods). We found that one distinguishing feature of the partial EMT trajectory was its broad variation in the EMT signature, paired with a consistent stemness signature. The top ancestors of the partial EMT trajectory exhibited a more pronounced EMT score variation than those of the high EMT fate (Levine test, p < 1e-10 for days 1-8) (Fig 4C, Table S4 and Methods). In contrast, the partial EMT trajectory exhibited a level of variation in stemness scores similar to, or even less than, that of the high EMT trajectory (Levine test, p > 0.05 on days 1, 4, and 8). On days when significant differences did occur (Levine test, p < 1e-5 on days 2 and 3), the variations were more significant in the high than the partial EMT trajectory (Fig 4D and Table S6).

To further explore the interplay between EMT and stemness signatures, we examined the joint distributions of these signatures at various time points (Methods). Our analysis revealed that the three trajectories during days 2 to 8 occupied different regions in the two-dimensional EMT and stemness score space. Specifically, the EMT signature predominantly distinguished between the low and high EMT trajectories, whereas a pronounced stemness signature demarcated the partial EMT trajectory from the other two (Fig 4F). Additionally, within the low EMT subset, a consistent positive correlation between EMT and stemness signatures was observed from days 1 to 8 (Pearson coefficients ranging from 0.22 to 0.44). In contrast, the high EMT subset presented a negative correlation between EMT and stemness signatures (Pearson coefficients ranging from - 0.3 to -0.6) (Table S8). This analysis reveals that cells with marked EMT signatures, whether extremely low or high, display reduced stemness. This trend is in line with earlier research suggesting that cells moving towards a distinctly differentiated state, whether closer to a pure E or M state along the EMT continuum, tend to exhibit less stemness^22^.

### Leveraging CRISPR Screening for Validation of Key Early Predicted Genes in EMT

To validate our identified trajectories, we compared our findings with a recent study that reported a substantial induction of the partial EMT fate following TGF-beta treatment in a background of PRC2 dysfunction, which was conducted across various epithelial cell lines including HMLER and MCF10A cells^35^. The 2D gene expression maps showed that the levels of *EED* and *EZH2*—key constituents of PRC2—were notably diminished in areas aligning with the high-probability regions for the partial EMT trajectory (Fig 5A). To validate this observation, we quantified the expressions of these genes across the three trajectories. Both genes exhibited distinct average expression trends, each distinctly demarcated by non-overlapping 95% confidence intervals. Importantly, there was a noticeable decline in *EED* and *EZH2* expressions, predominantly within the partial EMT trajectory (Fig 5B). Concurrently, within the top ancestors of the partial EMT, there was a discernible contraction in the distribution of *EED* expression, marked by a decrease in the number of cells exhibiting high gene expression, which is evidenced by a shift in the mean of the distribution (Fig 5C and Table S9). Similar patterns were observed for the *EZH2* gene (Fig 5A-C).

**Figure 5.**
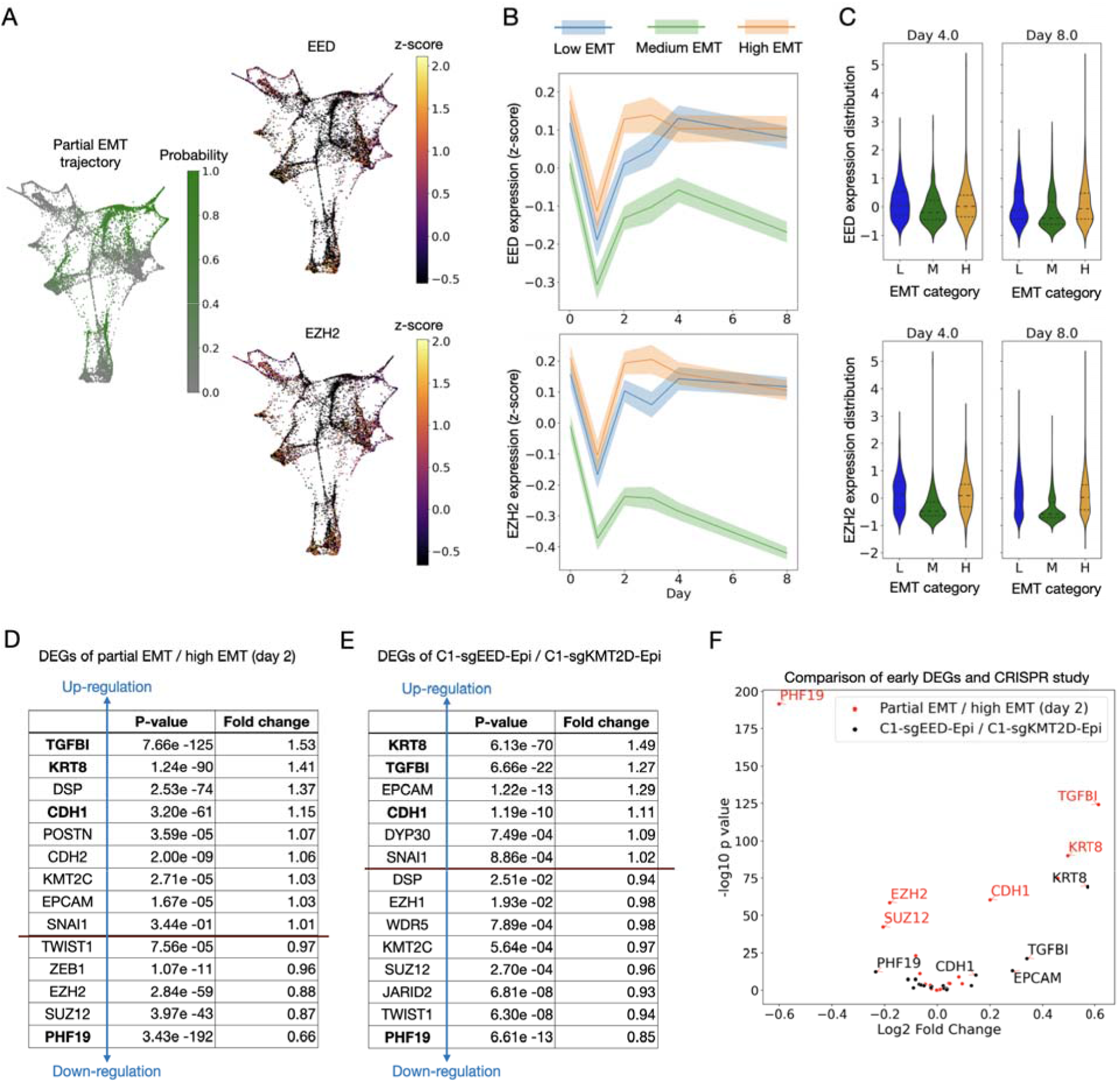
Early predictors of EMT fate through early DEGs analysis and CRISPR knock-out screening. (A-C) *EED* and *EZH2* gene expression analysis. (A) Color maps display the ATF distributions for the partial EMT trajectory alongside the expression levels of the *EED*/*EZH2* genes across all cellular states from day 0 to day 8. In the trajectory map, the color gradient signifies probability, while in the gene expression map, it indicates the level of gene expression. (B) Line plots illustrate the average dynamics of *EED*/*EZH2* gene expression over. Shaded regions denote the 95% confidence intervals. (C) Violin plots depict day-to-day distribution of expression levels for *EED*/*EZH2* gene among the top ancestor cells. (D-E) Two tables presenting the DEGs of EMT-related genes for two distinct datasets. Table (D) depicts our predicted DEGs at day 2 between the partial EMT and high EMT trajectories derived from the Deshmukh *et al*. dataset. Table (E) illustrates the DEGs in epithelial cells with *EED* knock-out (C1-sgEEG-Epi) compared to *KMT2D* knock-out cells (C1-sgKMT2D-Epi) using data from the Zhang *et al*. CRISPR study. (F) Volcano plot of the differential gene expression analysis conducted on the two datasets, as outlined in tables (D) and (E). Red dots indicate our early phase gene expression predictions, while black dots represent results from the CRISPR knockout study.

In line with these findings, the CRISPR screen study revealed that knocking out *EED* and *EZH2* promotes a partial EMT state with increased stemness^35^. This study was performed using the HMLER cell line, which, like MCF10A, is an immortalized human mammary epithelial cell line and exhibits similar changes in gene expression during TGF-beta-induced EMT as the MCF10A cell line^35,54^. Therefore, we curated an EMT-related gene list from both the time course data^18^ and the CRISPR screen study^35^ (Methods). Two mesenchymal states were identified in the CRISPR study: C1-sgEED-Mes (partial EMT with *EED* gene knockout) retained some epithelial traits, while C1-sgKMT2D-Mes (high EMT with *KMT2D* knockout) lacked them. We then examined the differential expression of the curated gene set between the partial and high EMT fates in our dataset on day 8, and between the C1-sgEED-Mes and C1-sgKMT2D-Mes cells in the CRISPR screen data (Fig S11). Remarkably, three out of the top four ranked genes—*TGFBI*, *POSTN*, and *KRT8*—were significantly upregulated in the partial EMT state in both datasets (*t*-test, p < 1e-10; fold changes in Table S11). This analysis further supports our characterization of the partial and high EMT fates within our dataset.

We then evaluated differential gene expression patterns between each pair of cellular states in our data, weighted according to their ancestral distributions one day after TGF-beta treatment. We compared our findings with a standard differential gene expression analysis conducted on two groups of CRISPR knockout epithelial cells, C1-sgEED-Epi and C1-sgKMT2D-Epi (Fig S11 and Methods). These two groups of epithelial cells were the ancestral cells for their respective EMT fates: the C1-sgEED-Mes and C1-sgKMT2D-Mes cells, respectively^35^. Notably, four of the top five ranked genes overlapped between our and the CRISPR study—*TGFBI*, *KRT8*, and *CDH1* were significantly upregulated, while *PHF19* was significantly downregulated in the partial EMT trajectory (*t*-test, p < 1e-9; fold changes are in Fig 5D-F). The concordance observed between our predictions and the results from the CRISPR screen study partially validates our inference approach of predicting early key genes in EMT.

Our methodology leverages the inherent heterogeneity of cellular states that culminate in diverse cell fates under a single EMT inducer. This approach enables the identification of crucial early-stage genes that govern specific cell destinies, effectively circumventing the necessity for extensive pre-existing biological knowledge when selecting a specific EMT inducer or applying CRISPR to knock out a specific gene for a corresponding cell fate. We further applied our early DEGs analysis to two comprehensive gene sets – the ones that we employed to delineate stemness and proliferation patterns^55,56^. Our results highlighted that in the early phase of TGF-beta induced EMT, genes such as *CENPF*, *CKS1B*, and *MKI67* were significantly upregulated in the ancestors of the high EMT state on day 2 (*t*-test, p < 1e-10 and fold changes 2.16, 1.80, and 1.76, respectively) (Fig 6A). Similar patterns were observed on days 1 and 3 (Fig S12). In contrast, in the early cellular states of the partial EMT state, genes like *LAMA3*, *LAMB3*, and *ITGB4* were prominently expressed on day 2 (*t*-test, p < 1e-10 and fold changes 1.66, 1.55, and 1.50, respectively) (Fig 6B). Again, similar trends were observed on days 1 and 3 (see Fig S12 for details). Our findings concerning the increased expression of *LAMA3*, *LAMB3*, and *ITGB4* aligns with prior research that identified their role in demarcating cancer stem cell-enriched populations in a partially mesenchymal state^57,58^. Our methodology provides novel insights by enabling the identification of DEGs across a temporal spectrum (Fig 6CD). For instance, our time-resolved analysis reveals that the differential expression of *ITGB4* in the partial EMT state, when compared to the high EMT state, is more pronounced during the early stages than it is in the later phases of EMT (Fig 6D). When comparing day 2 to day 8, the fold changes were 1.50 vs 1.09, respectively (Fig 6B). These findings potentially underscore crucial moments for timely interventions to influence the direction of EMT evolution.

**Figure 6.**
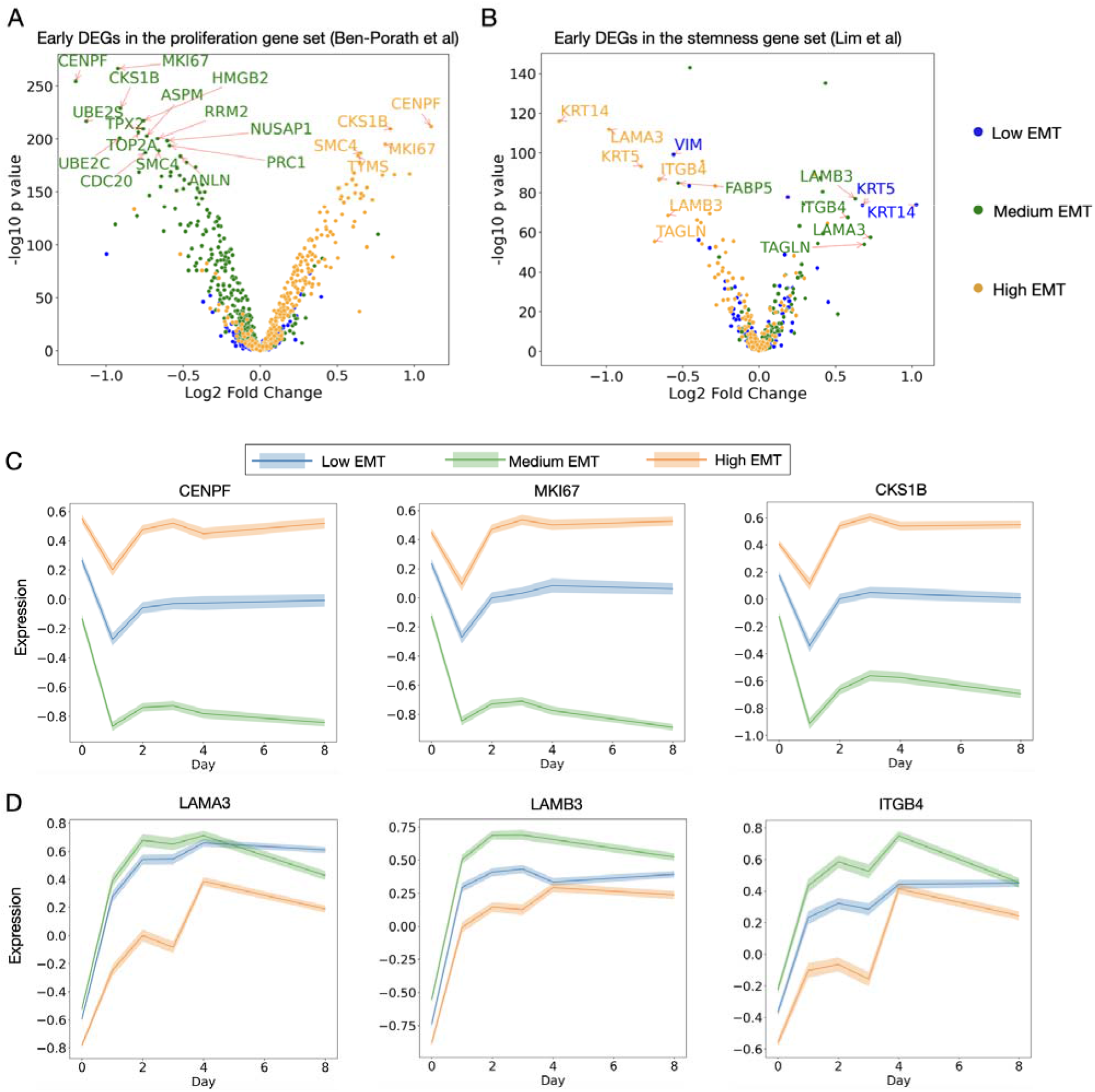
Temporal analysis of stemness and proliferation-associated differential gene expression. (A-B) Early DEGs of proliferation-related genes (A) and stemness-related genes (B). Distinct color codes showcase the differential gene expressions in cell states from a specific trajectory when contrasted with the combined cell states of the remaining two trajectories. As an example, green indicates genes with differential expression in the partial EMT trajectory compared to the joint low and high EMT trajectories. While the shown instance is based on day 2 data, comprehensive results spanning days 1-3 can be found in Supplementary Fig S12. (C-D) Line plots illustrate the average dynamics of *CENPF*, *CKS1B*, and *MKI67* (C) and *LAMA3*, *LAMB3*, and *ITGB4* (D) gene expression over. Shaded regions denote the 95% confidence intervals.

## Discussion

ScRNAseq data provides deep insights into the intricate heterogeneity of cellular states^23,24^. However, a key limitation of this type of data prevents tracking of individual lineages due to the need to sacrifice cells at each data collection time point, thus complicating the reconstruction of diverse trajectories from time series scRNAseq datasets. To circumvent this limitation, we used WOT to reconstruct lineages at the single cell level from EMT time series scRNAseq data^18^, allowing us to identify three separate trajectories culminating in distinct EMT outcomes.

Of the three trajectories, the low-EMT trajectory is distinguished by a slight decrease in EMT scores as compared to the high EMT outcome. Cells in the early stages of this trajectory exhibit pronounced epithelial characteristics, with high expression of CDH1 and low levels of CDH2.

Notably, the expression of these markers remains consistent over time (Fig 3C). These findings suggest that this trajectory could represent a failed EMT process rather than a mesenchymal-epithelial transition (MET), where cells revert from a mesenchymal to an epithelial state^59^.

The high EMT trajectory is characterized by a consistently high EMT signature from day 0 onwards, with minimal variations in EMT scores throughout the trajectory (Fig 3B). Determining the EMT signature using the 76 GS method, which assigns weights based on each gene’s correlation with *CDH1* expression levels. Consequently, the persistent high EMT score aligns the continuous low expression of *CDH1* observed within this trajectory (Fig 3C). Despite the low CDH1 expression, early-stage cells in this trajectory maintain high levels of other epithelial markers, notably *EPCAM*, while lacking mesenchymal markers such as *CDH2* (Fig 3B and Fig S7). This pattern suggests that the cells do not start in a fully mesenchymal state; it is only during treatment that they develop a full mesenchymal profile, confirming their progression through the full EMT process.

The partial EMT trajectory, aside from its medium EMT score, is marked by a low proliferation signature alongside elevated stemness. In addition, its marked variability in EMT signatures underscores its inherent EMT heterogeneity, suggesting a spectrum of phenotypes rather than a single partial EMT state. In our early DEG analysis, top-tier genes like *TGFBI*, *KRT8*, and *CDH1* were highlighted for their increased expression early in the partial EMT phase, aligning with findings from an independent CRISPR study. Expanding our DEG analysis to larger gene sets related to stemness and proliferation, we unveiled a novel set of pivotal early-stage genes, of which *ITGB4* was particularly noteworthy, that distinguish the partial EMT from the high EMT trajectories.

Through our analysis, we have uncovered several novel findings that differentiate individual EMT trajectories. These insights, when integrated with existing EMT research, can offer a more comprehensive view of the EMT landscape. First, we found that the low EMT trajectory is determined early on, within a day of treatment. This result suggests that the initial state of these cells renders them resistant to TGF-beta, providing novel insights into two prior studies on EMT resistance: one study identified a subpopulation of epithelial cells with similar capabilities to receive and process TGF-beta signals but exhibited a notably weaker downstream response compared to more sensitive cell populations^35^. Another study revealed that sustained *EPCAM* expression acts as a marker for epithelial clones in metastatic breast cancer that resist EMT induction, a trait shaped by the interplay between human *ZEB1* and its target, *GRHL2*^60^. Furthermore, TGF-beta is well-known as a promoter of mesenchymal gene expression and a suppressor of cell proliferation^44–46^. However, as far as we are aware, the pronounced reaction to TGF-beta within the partial EMT trajectory, leading to a significant reduction in proliferation and a moderate increase in mesenchymal markers, has not been emphasized in earlier research, thereby indicating a promising area for future exploration.

Additionally, we found a significant downregulation of *EED* and *EZH2* within the partial EMT trajectory. These findings are consistent with those reported by Zhang *et al.*^35^, where the addition of TGF-beta, along with compounds EED226 (inhibiting *EED*) or tazemetostat (inhibiting *EZH2*), steered cells toward a partial EMT state. The capacity of our methodology to predict the full gene expression trajectory at individual time points revealed an initial divergence in *EED* and *EZH2* expression levels at the onset of EMT (Day 0), reflecting intrinsic heterogeneity among the ancestral cells of each trajectory. Notably, this variation in expression intensified between days 2 and 4. As TGF-beta was the exclusive treatment applied during this phase, it is plausible that these alterations stem from the cellular reaction to TGF-beta, potentially amplifying the initial heterogeneity. Further experimental validation of our findings is the topic of future investigation.

Lastly, leveraging the heterogeneity of cellular responses to TGF-beta induced EMT, our method effectively pinpoints early differentially expressed genes across distinct EMT trajectories from a broad set of candidates. For instance, we distinguished *ITGB4*, *LAMA3* and *LAMB3* due to their pronounced differential expressions in the early stages of the partial versus high EMT trajectories. As previously highlighted, *ITGB4* serves as an integrin subunit that interacts with specific matrix proteins, while *LAMB3* and *LAMA3* engage with different integrin subunits than does *ITGB4*^61,62^. Future validation of our findings could employ cell surface markers encoded by these genes to isolate early-phase cells to can observe their responses under a consistent TGF-beta treatment timeline.

In summary, we uncovered three distinct trajectories leading to TGF-beta induced EMT. Most interestingly, the partial trajectory is characterized by an intermediate EMT score, elevated stemness, pronounced hypoxia response, and decreased proliferation. The reduced expression levels of the *EED* and *EZH2* genes in this trajectory are consistent with the findings of a CRISPR study. Finally, we identified key genes upregulated in the early phases of cells along this trajectory, which had not been previously linked to early EMT regulation. Our approach represents a computational method to circumvent the need for lineage tracing in single cell experiments and sheds light onto the diverse trajectories leading to different EMT fates.

In this study, we focused on gene expression alterations following EMT induction through scRNA seq data analysis. Such a scope, however, might not capture the complete EMT landscape. First, although we’ve identified early divergences in EMT trajectories, actual phenotypic changes may become apparent later, potentially attributable to differences between RNA expression and protein levels due to RNA-protein discordance^63^. In addition, variabilities in gene expression can arise from intrinsic noise^64^, extracellular environmental changes^65,66^, and differing epigenetic states^67^. The overall approach described in this study, when applied to suitable time series datasets, could help disentangle the role of the aforementioned mechanisms in driving differing responses to EMT induction and in driving heterogeneity in various cell fate transition processes^68^. Building on the success of applying OT to spatial data^69,70^, we suggest using this approach on spatial transcriptomics from tumors, paired with latent time inferences, can elucidate the tumor environment’s influence on EMT heterogeneity^71^. Furthermore, the same approach may be applied to datasets that jointly profile RNA expression and chromatin accessibility^72^ at different time points to probe the contribution of epigenetic states in driving distinct transcriptional responses^60^, and whether epigenetic changes precede gene expression changes during a cell fate transition.

Additionally, WOT is based on optimal transport theory, which assumes that cells traverse the gene expression space via the shortest overall distance^26,27^. This foundational assumption serves as an unbiased starting point for cell state transitions^28^. Future refinements could integrate prior knowledge of specific gene expression changes, adjusting gene distances based on this knowledge. This approach would allow us to leverage WOT more adaptively, inferring unknown system parts from existing biological understanding. Another requirement of the WOT approach is the use of time-series scRNAseq data^28,73^, a need which is restricted by the cost associated with conducting multiple time-point experiments. To overcome this limitation, WOT could potentially be extended to a pool of scRNAseq data without knowledge of the timeline. This advancement would entail inferring the latent time variable prior to using WOT. Given the prevalent availability of EMT data of this type^58,74^, this extension could lead to the effective identification of key genes that demonstrate significant expression across various cell lines when exposed to different EMT inducers.

## Materials and Methods

### ScRNA-Seq Data Analysis

The single cell RNA-seq datasets analyzed here were obtained from published studies^18,35^. For the dataset from Deshmukh *et al.*, we used the processed sequencing data made available by the authors; the raw sequencing reads are available from the NCBI Sequence Read Archive^75^ (BioProject accession PRJNA698642). From the dataset from Zhang *et al.,* processed single cell RNA-seq profiles of HMLER cells subjected to EED / EZH2 knockout were downloaded from the Gene Expression Omnibus (GEO) database^76^ (GEO accession GSE158115). The procedures for quality control, data normalization, batch correction, and other steps for this dataset were performed as in the original paper^35^.

We used two methods to reduce the dimensionality of the single cell RNA-seq datasets. Following the approach in^35^, we performed Uniform Manifold Approximation and Projection (UMAP) dimensional reduction using Seurat v3.61^77^ for the HMLER dataset. For the MCF10A dataset, we followed the approach in Deshmukh *et al.*^18^ and used force-directed layout embedding (FLE)^78^, a type of graph visualization algorithm, to generate a nearest-neighbor graph built from the single cell gene expression data. In the FLE visualization, each cell is portrayed as a node and is linked to its ‘k’ closest neighboring cells. The application of FLE allows for an intuitive visualization of cellular relationships, as it induces an attraction between nodes (cells) connected by an edge and a repulsion between nodes that lack an interconnecting edge. The FLE visualization was carried out using the Pegasus tool [Pegasus 1.8.0]. MCF10A cells on day 8 were divided into three distinct subpopulations (shown in Fig. 2A, day 8: blue, green, and orange points) by applying the k-means clustering algorithm (k = 3) to the FLE visualization coordinates of the day 8 subset. The choice of k = 3 was primarily informed by our existing biological understanding of the three EMT states—epithelial, partial, and mesenchymal—referenced in earlier studies^58^. To further substantiate this choice, we investigated the results of k values ranging from 2 to 6, using Silhouette scores as a metric to evaluate the clustering performance. Silhouette scores range from -1 to +1, with a higher score suggesting that objects in the same cluster are closer to each other than to objects in neighboring clusters^79^. Among these choices, k = 3 resulted in the second-highest Silhouette score of 0.65, only slightly behind k = 4, which had a score of 0.67 (Fig S13). However, k = 3 produced balanced cluster sizes of 589, 746, and 556 cells, while k = 4 resulted in a notably smaller cluster with just 32 cells alongside clusters of 589, 721, and 549 cells. Taking both Silhouette scores and cell cluster sizes into account, we selected k=3 as the most appropriate value.

### Inferring trajectories with Waddington-OT

We employed Waddington-OT (WOT)^28^ to calculate the transition probabilities between cells across successive time points as done previously^28^. Specifically, our input data consisted of the normalized gene expression matrix alongside the day annotations from the preprocessed scRNAseq data. By default in WOT, cellular states—defined by the gene expression matrix—are assumed to be uniformly distributed on each day, though adjusted by growth rates as delineated in Schiebinger *et al.*^28^. To calculate the growth rate, we used WOT’s inherent logistic function, which transforms the proliferation (or apoptosis) signature score into the birth (or death) rate for each cell. Subsequently, the net growth rate was derived from the difference between the birth and death rate. In line with WOT, we used the G2M hallmark and the apoptosis hallmark gene sets from MsigDB^80,81^ to compute the proliferation and apoptosis signature scores, respectively.

Furthermore, the cell state distributions, covering two consecutive days, were then used as the marginal distributions in our unbalanced transport optimization problem^27,28^. The cost function in this optimization task is given by the squared Euclidean distance. By default in WOT, we conducted the unbalanced transport optimization with parameters ε=0.05, λ1=1, and λ2=50, incorporating a single iteration of growth rate learning^28^. The rationale and stability of these parameters are discussed in the WOT method and have been validated in the quantification and statistical analysis section of the original paper^28^.

We then combined the OT maps predicted from every pair of adjacent time points, drawing from the time-series scRNA-seq data. This integration enabled us to generate a comprehensive probability map that traces each cell’s likelihood of being an ancestor at earlier time points and transitioning to one of the terminal cell states following TGF-beta treatment. Building on this complete map, we summed up all probabilistic trajectories at single cell resolution that culminate in different fate subpopulations. As a result, we calculated the ancestor-to-fate (ATF) distributions for each early time point, specifying the likelihood of each ancestor differentiation into distinct fate subgroups.

### Visualizing divergence of fates with barycentric coordinates

For visual representation of ATF distributions, we employed the ‘tmap_model.fates’ function from WOT^28^. Within this visualization approach, cell states are represented by a three-dimensional vector (p_1_, p_2_, p_3_) adhering to the condition p_1_+ p_2_ + p_3_ = 1. This vector characterizes the ATF distribution of a cell state transitioning into the orange, green, and blue fates. Notably, by day 8, all cells within a particular cluster show full commitment to a specific fate, as evidenced by a value of 1 in the corresponding position; for instance, cells within the orange cluster are denoted by the vector (1, 0, 0). We visualized these three-dimensional probability vectors by projecting them onto a two-dimensional triangle using barycentric coordinate representation. In this coordinate system, each vertex of the triangle is associated with a specific fate, and the cells are positioned based on their relative transition probabilities. We thus designated (a, b, c) as the vertices of the triangle on the 2D plane. The projection of the vector (p_1_, p_2_, p_3_) onto this triangle utilized the convex combination of the vertices’ locations (a, b, c) with the coefficients (p_1_, p_2_, p_3_), resulting in the location given by p_1_*a + p_2_*b + p_3_*c. Given that p_1_ + p_2_ + p_3_ = 1, each probability vector corresponds to a point within the triangle. Cells situated at the triangle’s center, represented by p = (1/3, 1/3, 1/3), denote undetermined states, suggesting an equal likelihood of differentiating into any of the fates. Conversely, cells closer to the vertices indicate a strong commitment to transition into the fate associated with the corresponding vertex.

We adopted a 75% commitment as a threshold for transitioning into a fate, given its conventional use as an upper quantile indicative of high probability. For example, for the orange fate, this threshold is symbolized as p= (0.75, p_2_, p_3_) and is represented as a line delineating a region within the upper right corner with the vertex (1,0,0). Within this area, all cells display over 75% commitment to transition into the blue fate subpopulation by day 8. This series of barycentric plots dynamically portrays the temporal evolution of major ancestral contributions, providing an intuitive visualization of our predictions of cell fate trajectories over time. To ensure robustness of our findings, we investigated the effects of adjusting the threshold value upwards to 90% in 3% increments by reanalyzing our data for each threshold. We set 90% as the highest threshold due to the scarcity of cells beyond this point, particularly in early time points. For instance, at a 91% threshold on day 0, there were only 2 top ancestor cells for the medium EMT fate and 1 top ancestor cell for the high EMT fate. In Supplement S1, we present the outcomes corresponding to these varied thresholds, emphasizing that our conclusions remain consistent irrespective of the specific threshold utilized.

### Validating trajectories with geodesic interpolation

To assess the reliability of the WOT-inferred cellular trajectories, we employed a cross-validation approach as outlined in the “validation of geodesic interpolation” section in the WOT methods^28^. In this approach, for every test data time point, its adjacent time points were used as training data, creating a series of consecutive time point triplets. Utilizing WOT’s built-in function, ‘wot.graphics.plot_ot_validation_summary_stats’, we then computed the Wasserstein 2-distance between distributions^40^. This distance measures the least amount of “cost” required to transform one distribution into another, with “cost” denoted by the squared Euclidean distance of moving a unit mass within the distribution. We thus obtained the deviation, using the Wasserstein 2-distance, between the inferred and the actual midpoint test data. This deviation is depicted by the blue curve in Fig S3C. As per WOT’s default settings, our results were benchmarked against three null models: 1) the Wasserstein distance between the test data and the data from the preceding day (depicted by green curves); 2) the Wasserstein distance between the test data and the data from the subsequent day (illustrated by purple curves); and 3) the Wasserstein distance between the test data and the estimated data from the interpolation based on a random map (highlighted by orange curves). In the random map model, WOT assumed all cell state transitions to be equally probable. Compared to these null models, our inferred distributions consistently exhibited the smallest deviations at most time points. For instance, the WOT-interpolated deviation values for days 1-4 are 8.16, 6.5, 6.85, and 6.85, which are smaller than the random coupling interpolation values of 8.85, 7.78, 7.08, and 7.4 for the same days (Fig S3C). Only on day 4 did our interpolation’s deviation surpass that of the day 4 to day 8 data comparison (6.85 vs 5.47). This outcome may be due to the near completion of cell transition by day 4 (Fig 1A).

### Assessing EMT Scores: The 76 GS and KS Methods

EMT scores were calculated using two distinct methodologies, each employing different gene sets and metrics. The consistency between these methods has been verified through a comparative study involving multiple individual samples^39^. In the 76 GS method^36,38^, we computed the EMT score as a weighted sum of the expression levels of 76 EMT-related genes. The weight assigned to each gene was determined by its correlation with the *CDH1* (E-cadherin) expression level. The scores were subsequently adjusted such that the mean is 0. As a result, a negative score signifies that a cell’s EMT state is closer to the epithelial (E) state than the mesenchymal (M) state. We then rescaled the scores by taking their negatives, thus aligning the direction of the scores with the progression from the E to M state. The second method, known as the KS method, was initially established based on a comparison between the cumulative distribution functions (CDFs) of the E and M signatures^37^. According to this method, the EMT score is computed as the maximum difference between the two CDFs, i.e. the CDF of the M signature minus the CDF of the E signature. Therefore, a positive score for a sample indicates its closeness to the M state, and vice versa.

### Computation of cellular signature scores by ssGSEA

For determining the expression level of the stemness signature, we adopted gene sets from Lim *et al.*^55^, specifically designed to distinguish between stemness and mature cell signatures by investigating mammary stem and luminal cells. Additionally, we employed the proliferation gene set from Ben-Porath *et al.*^56^. This set was compiled by merging three distinct gene groups: those that are functionally involved in proliferation, those with cyclical expression within the cell cycle, and those that were instrumental in the clustering of proliferative subpopulations within human breast tumor expression data. For signaling pathway analysis, we investigated the gene sets of hypoxia, G2M, mitotic spindle, TGF-beta, PI3K-AKT-mTOR, Wnt, and IL6-JAK-STAT3 hallmarks from MSigDB^80,81^. We performed single-sample Gene Set Enrichment Analysis (ssGSEA) on all gene sets using GSEAPY (v1.0.4), a Python package^82^. The enrichment scores for gene sets were transformed into z-scores, with adjustments made by shifting the mean and normalizing by the standard deviation.

### Inference of average expression levels of cellular/genetic signatures in the diverse EMT trajectories

To determine average signature expressions across different trajectories, we employed the ‘update_trends_vis’ function from WOT, as detailed in Schiebinger *et al*^28^. Specifically, this function requires a single trajectory, identified as described in the “Inferring trajectories with Waddington-OT” section, along with signature scores for each individual cell. These scores were determined either by calculating the cellular signature scores as described above, or by using gene expression scores extracted from individual levels within the gene expression matrix, as elaborated in the “scRNA-Seq Data Analysis” section. The ‘update_trends_vis’ function subsequently computes the average scores for each distinct EMT trajectory at every time point by calculating the cumulative score for each cell state weighted by the probability of that cell state serving as an ancestor for each of the three fates. By iterating this computation for data from each time point, this method reconstructs the varied trajectories of score dynamics, with each path pointing towards a unique fate (Fig 3A and Fig S4-6). To elucidate the correlations between different signatures, we paired various score trajectories and projected them onto a 2D plane (Fig 3C and Fig S7B).

### Quantitative analysis and visualization of top ancestor signature distributions

For each top ancestor group, we visualized the distribution of cellular signatures using violin plots created with the seaborn tool^83^. Additionally, we used seaborn’s scatter plots to study the joint distributions of various cellular signature pairs within each of the three top ancestor groups and the undetermined ancestors (Fig 4D). Utilizing the pandas library^84^, we computed a comprehensive set of relevant statistics, including means, variances, and t-tests, based on the cellular signature scores of the top ancestor sets (Tables S4, S6, and S7). In our assessment of variances for the EMT signature score, we considered both the inclusion and exclusion of outliers. These outliers were identified as cells that were positioned outside the 1^st^ to 99^th^ percentile range. Additionally, we employed the ‘Series.corr’ function from pandas to determine Pearson correlations from pairwise cellular signature scores within each top ancestor group (Table S8).

### Cellular transcriptomic heterogeneity measured by pairwise distances

To examine the variability within cellular transcriptomes, we computed the pairwise distance between each cell state based on the normalized RNA expression. Adapting the methodology from Hinohara *et al*^52^, we obtained the mean pairwise distance by summing these distances and dividing by the total number of cell state pairs. However, diverging from Hinohara et al’s approach, this distance was weighted by multiplying the probabilities of the cellular states in each pair, guided by ATF distributions. This approach yields distinct expected pairwise distances that correspond to the specific ATF distribution for each fate at every time point. These pairwise distances and their associated probabilistic weights are also used to calculate the variances.

These heterogeneity computations were carried out based on different RNA expression spaces defined by the gene sets we used to define heterogeneity. We used the full gene set, provided by all genes in the scRNAseq dataset, the EMT gene set, and the DEG set identified from comparisons between the high and medium EMT fate populations (Methods, differential expression analysis). For consistency in dimensionality, given that we used 72 genes in the EMT gene set (the original set had 76 genes but only 72 overlapped with the genes used in our scRNAseq data), we performed Principal Component Analysis (PCA) to select the 72 principal components for the distance in space of the full gene set. We also selected the 72 top-ranked genes in the differentially expressed gene sets (Table S1). As a validation, we also calculated the pairwise distance in the original dimension (15,000) of the full gene set to ensure consistency with the distance measured in the 72-dimensional space derived from PCA (Fig S9 and Table S3).

### Differential expression analysis

In order to identify differentially expressed genes (DEGs) between the partial EMT and high EMT fates, we compared the expression level of each gene between these two cell populations using the ‘stats.ttest_ind’ function from scipy^85^ to conduct a t-test between the groups, producing p-values and t-statistics. Additionally, we calculated the fold change between the mean gene expression levels from one group to the other. Drawing from the principle of dimensional consistency highlighted in the previous section, we curated two distinct gene sets, each with the top 36 genes. This amounts to a total of 72 genes in the DEG set, consistent with the size of the EMT gene set. These genes are significantly differential by their heightened expression levels in either the medium EMT or high EMT states (Table S1). We followed two criteria for gene selection: (1) selecting the smallest 36 p-values with a fold change greater than 100, and (2) choosing the largest 36 fold-changes with p-values smaller than 1e-7 (Table S1). By applying these two criteria, we obtained two sets of genes (Table S1) for our downstream analysis. Importantly, all conclusions drawn from both sets of genes were consistent. The main text presents the results derived from the first gene set, while detailed results from the second gene set can be found in Supplementary Fig S9 and Table S2. For inferring early DEGs between the partial EMT and complete EMT trajectories, we employed the ‘wot.tmap.diff_exp’ function from the WOT package^28^. This function calculates the differential expression of each gene across two cell sets, which includes all cell states collected at a specific time point. The differentiating factor between these two cell sets are the weights assigned to their cell states, which were determined based on their probabilities of transitioning into the corresponding fates. Notably, by day 8, when the cell states are determined by their fates, these weights take on a binary value (either 1 or 0), aligning with the conventional approach to identifying DEGs. For the dataset from the CRISPR knockout study, where all cell states are definitively determined and have been preprocessed by Seurat v3.61^77^, the ‘findMarkers’ function from the Seurat package in R was used.

### Code availability

All code used to process data and generate figures is available on a public Github repository at https://github.com/Michorlab/OT-EMT.

## Acknowledgments

We would like to thank the Michor lab, Dr. Petra den Hollander, Dr. Sahand Hormoz, and Dr. Sendurai Mani for helpful discussions and comments. We gratefully acknowledge support of the Ludwig Center at Harvard and the Dana-Farber Cancer Institute’s Center for Cancer Evolution.

## References

1. Hay, E. D. An overview of epithelio-mesenchymal transformation. Acta Anat (Basel) 154, 8– 20 (1995).

2. Kang, Y. & Massagué, J. Epithelial-Mesenchymal Transitions: Twist in Development and Metastasis. Cell 118, 277–279 (2004).

3. Thiery, J. P. & Sleeman, J. P. Complex networks orchestrate epithelial–mesenchymal transitions. Nat Rev Mol Cell Biol 7, 131–142 (2006).

4. Nieto, M. A., Huang, R. Y.-J., Jackson, R. A. & Thiery, J. P. EMT: 2016. Cell 166, 21–45 (2016).

5. Dongre, A. & Weinberg, R. A. New insights into the mechanisms of epithelial–mesenchymal transition and implications for cancer. Nat Rev Mol Cell Biol 20, 69–84 (2019).

6. Wendt, M. K., Taylor, M. A., Schiemann, B. J. & Schiemann, W. P. Down-regulation of epithelial cadherin is required to initiate metastatic outgrowth of breast cancer. MBoC 22, 2423–2435 (2011).

7. Wang, Y. & Zhou, B. P. Epithelial-mesenchymal Transition---A Hallmark of Breast Cancer Metastasis. Cancer Hallm 1, 38–49 (2013).

8. Lu, M., Jolly, M. K., Levine, H., Onuchic, J. N. & Ben-Jacob, E. MicroRNA-based regulation of epithelial–hybrid–mesenchymal fate determination. Proceedings of the National Academy of Sciences 110, 18144–18149 (2013).

9. Zhang, L., Zhou, F. & ten Dijke, P. Signaling interplay between transforming growth factor-β receptor and PI3K/AKT pathways in cancer. Trends in Biochemical Sciences 38, 612–620 (2013).

10. Roche, J. The Epithelial-to-Mesenchymal Transition in Cancer. Cancers 10, 52 (2018).

11. Vasaikar, S. V. et al. EMTome: a resource for pan-cancer analysis of epithelial-mesenchymal transition genes and signatures. Br J Cancer 124, 259–269 (2021).

12. Taipale, J. & Beachy, P. A. The Hedgehog and Wnt signalling pathways in cancer. Nature 411, 349–354 (2001).

13. Katoh, Y. & Katoh, M. FGFR2-related pathogenesis and FGFR2-targeted therapeutics (Review). International Journal of Molecular Medicine 23, 307–311 (2009).

14. Moustafa, A.-E. A., Achkhar, A. & Yasmeen, A. EGF-receptor signaling and epithelial-mesenchymal transition in human carcinomas. FBS 4, 671–684 (2012).

15. Espinoza, I. & Miele, L. Deadly crosstalk: Notch signaling at the intersection of EMT and cancer stem cells. Cancer Lett 341, 41–45 (2013).

16. Heldin, C.-H., Vanlandewijck, M. & Moustakas, A. Regulation of EMT by TGFβ in cancer. FEBS Letters 586, 1959–1970 (2012).

17. Cook, D. P. & Vanderhyden, B. C. Context specificity of the EMT transcriptional response. Nat Commun 11, 2142 (2020).

18. Deshmukh, A. P. et al. Identification of EMT signaling cross-talk and gene regulatory networks by single-cell RNA sequencing. Proceedings of the National Academy of Sciences 118, e2102050118 (2021).

19. Waddington, C. H. The Strategy of the Genes. (Routledge, 2014).

20. Ferrell, J. E. Bistability, Bifurcations, and Waddington’s Epigenetic Landscape. Current Biology 22, R458–R466 (2012).

21. Mani, S. A. et al. The Epithelial-Mesenchymal Transition Generates Cells with Properties of Stem Cells. Cell 133, 704–715 (2008).

22. Jolly, M. K. et al. Towards elucidating the connection between epithelial–mesenchymal transitions and stemness. Journal of The Royal Society Interface 11, 20140962 (2014).

23. Mazutis, L. et al. Single-cell analysis and sorting using droplet-based microfluidics. Nat Protoc 8, 870–891 (2013).

24. Tanay, A. & Regev, A. Scaling single-cell genomics from phenomenology to mechanism. Nature 541, 331–338 (2017).

25. Wagner, D. E. & Klein, A. M. Lineage tracing meets single-cell omics: opportunities and challenges. Nat Rev Genet 21, 410–427 (2020).

26. Villani, C. Optimal Transport: Old and New. (Springer Berlin Heidelberg, 2016).

27. Chizat, L., Peyré, G., Schmitzer, B. & Vialard, F.-X. Scaling algorithms for unbalanced optimal transport problems. Math. Comp. 87, 2563–2609 (2018).

28. Schiebinger, G. et al. Optimal-Transport Analysis of Single-Cell Gene Expression Identifies Developmental Trajectories in Reprogramming. Cell 176, 928–943.e22 (2019).

29. Haghverdi, L., Büttner, M., Wolf, F. A., Buettner, F. & Theis, F. J. Diffusion pseudotime robustly reconstructs lineage branching. Nat Methods 13, 845–848 (2016).

30. Saelens, W., Cannoodt, R., Todorov, H. & Saeys, Y. A comparison of single-cell trajectory inference methods. Nat Biotechnol 37, 547–554 (2019).

31. Gorin, G., Svensson, V. & Pachter, L. Protein velocity and acceleration from single-cell multiomics experiments. Genome Biology 21, 39 (2020).

32. Bergen, V., Lange, M., Peidli, S., Wolf, F. A. & Theis, F. J. Generalizing RNA velocity to transient cell states through dynamical modeling. Nat Biotechnol 38, 1408–1414 (2020).

33. Qiu, Q. et al. Massively parallel and time-resolved RNA sequencing in single cells with scNT-seq. Nat Methods 17, 991–1001 (2020).

34. Lundgren, K., Nordenskjöld, B. & Landberg, G. Hypoxia, Snail and incomplete epithelial– mesenchymal transition in breast cancer. Br J Cancer 101, 1769–1781 (2009).

35. Zhang, Y. et al. Genome-wide CRISPR screen identifies PRC2 and KMT2D-COMPASS as regulators of distinct EMT trajectories that contribute differentially to metastasis. Nat Cell Biol 24, 554–564 (2022).

36. Byers, L. A. et al. An Epithelial–Mesenchymal Transition Gene Signature Predicts Resistance to EGFR and PI3K Inhibitors and Identifies Axl as a Therapeutic Target for Overcoming EGFR Inhibitor Resistance. Clinical Cancer Research 19, 279–290 (2013).

37. Tan, T. Z. et al. Epithelial-mesenchymal transition spectrum quantification and its efficacy in deciphering survival and drug responses of cancer patients. EMBO Molecular Medicine 6, 1279–1293 (2014).

38. Guo, C. C. et al. Dysregulation of EMT Drives the Progression to Clinically Aggressive Sarcomatoid Bladder Cancer. Cell Rep 27, 1781–1793.e4 (2019).

39. Chakraborty, P., George, J. T., Tripathi, S., Levine, H. & Jolly, M. K. Comparative Study of Transcriptomics-Based Scoring Metrics for the Epithelial-Hybrid-Mesenchymal Spectrum. Frontiers in Bioengineering and Biotechnology 8, (2020).

40. Ambrosio, L., Gigli, N. & Savare, G. Gradient Flows: In Metric Spaces and in the Space of Probability Measures. (Birkhäuser, Basel, 2008).

41. Bao, B. et al. The biological kinship of hypoxia with CSC and EMT and their relationship with deregulated expression of miRNAs and tumor aggressiveness. Biochimica et Biophysica Acta (BBA) - Reviews on Cancer 1826, 272–296 (2012).

42. Emami Nejad, A., et al. The role of hypoxia in the tumor microenvironment and development of cancer stem cell: a novel approach to developing treatment. Cancer Cell International 21, 62 (2021).

43. Lu, X. & Kang, Y. Hypoxia and Hypoxia-Inducible Factors: Master Regulators of Metastasis. Clinical Cancer Research 16, 5928–5935 (2010).

44. Massagué, J., Blain, S. W. & Lo, R. S. TGFβ Signaling in Growth Control, Cancer, and Heritable Disorders. Cell 103, 295–309 (2000).

45. Donovan, J. & Slingerland, J. Transforming growth factor-β and breast cancer: Cell cycle arrest by transforming growth factor-β and its disruption in cancer. Breast Cancer Research 2, 116 (2000).

46. Zhang, Y., Alexander, P. B. & Wang, X.-F. TGF-β Family Signaling in the Control of Cell Proliferation and Survival. Cold Spring Harb Perspect Biol 9, a022145 (2017).

47. Aban, C. E. et al. Downregulation of E-cadherin in pluripotent stem cells triggers partial EMT. Sci Rep 11, 2048 (2021).

48. Aggarwal, V. et al. P4HA2: A link between tumor-intrinsic hypoxia, partial EMT and collective migration. Advances in Cancer Biology - Metastasis 5, 100057 (2022).

49. Yao, Z. et al. TGF-β IL-6 axis mediates selective and adaptive mechanisms of resistance to molecular targeted therapy in lung cancer. Proceedings of the National Academy of Sciences 107, 15535–15540 (2010).

50. Akhmetshina, A. et al. Activation of canonical Wnt signalling is required for TGF-β-mediated fibrosis. Nat Commun 3, 735 (2012).

51. Carpenter, C. L. & Cantley, L. C. Phosphoinositide 3-kinase and the regulation of cell growth. Biochimica et Biophysica Acta (BBA) - Reviews on Cancer 1288, M11–M16 (1996).

52. Hinohara, K. et al. KDM5 Histone Demethylase Activity Links Cellular Transcriptomic Heterogeneity to Therapeutic Resistance. Cancer Cell 34, 939–953.e9 (2018).

53. Tyler, M. & Tirosh, I. Decoupling epithelial-mesenchymal transitions from stromal profiles by integrative expression analysis. Nat Commun 12, 2592 (2021).

54. Elenbaas, B. et al. Human breast cancer cells generated by oncogenic transformation of primary mammary epithelial cells. Genes Dev. 15, 50–65 (2001).

55. Lim, E. et al. Transcriptome analyses of mouse and human mammary cell subpopulations reveal multiple conserved genes and pathways. Breast Cancer Research 12, R21 (2010).

56. Ben-Porath, I. et al. An embryonic stem cell–like gene expression signature in poorly differentiated aggressive human tumors. Nat Genet 40, 499–507 (2008).

57. Bierie, B. et al. Integrin-β4 identifies cancer stem cell-enriched populations of partially mesenchymal carcinoma cells. Proceedings of the National Academy of Sciences 114, E2337–E2346 (2017).

58. Puram, S. V. et al. Single-Cell Transcriptomic Analysis of Primary and Metastatic Tumor Ecosystems in Head and Neck Cancer. Cell 171, 1611–1624.e24 (2017).

59. Yang, J. et al. Guidelines and definitions for research on epithelial–mesenchymal transition. Nat Rev Mol Cell Biol 21, 341–352 (2020).

60. Saini, M. et al. Resistance to mesenchymal reprogramming sustains clonal propagation in metastatic breast cancer. Cell Reports 42, 112533 (2023).

61. Giancotti, F. G. & Tarone, G. Positional Control of Cell Fate Through Joint Integrin/Receptor Protein Kinase Signaling. Annual Review of Cell and Developmental Biology 19, 173–206 (2003).

62. Spaderna, S., et al. A Transient, EMT-Linked Loss of Basement Membranes Indicates Metastasis and Poor Survival in Colorectal Cancer. Gastroenterology 131, 830–840 (2006).

63. Paul, I. et al. Parallelized multidimensional analytic framework applied to mammary epithelial cells uncovers regulatory principles in EMT. Nat Commun 14, 688 (2023).

64. Raser, J. M. & O’Shea, E. K. Noise in Gene Expression: Origins, Consequences, and Control. Science 309, 2010–2013 (2005).

65. Schmalhofer, O., Brabletz, S. & Brabletz, T. E-cadherin, β-catenin, and ZEB1 in malignant progression of cancer. Cancer Metastasis Rev 28, 151–166 (2009).

66. Bocci, F. et al. Toward understanding cancer stem cell heterogeneity in the tumor microenvironment. Proceedings of the National Academy of Sciences 116, 148–157 (2019).

67. Feinberg, A. P. & Levchenko, A. Epigenetics as a mediator of plasticity in cancer. Science 379, eaaw3835 (2023).

68. Francesconi, M. et al. Single cell RNA-seq identifies the origins of heterogeneity in efficient cell transdifferentiation and reprogramming. eLife 8, e41627 (2019).

69. Cang, Z. & Nie, Q. Inferring spatial and signaling relationships between cells from single cell transcriptomic data. Nat Commun 11, 2084 (2020).

70. Cang, Z. et al. Screening cell–cell communication in spatial transcriptomics via collective optimal transport. Nat Methods 20, 218–228 (2023).

71. Deng, Y. et al. Spatial profiling of chromatin accessibility in mouse and human tissues. Nature 609, 375–383 (2022).

72. Ranzoni, A. M. et al. Integrative Single-Cell RNA-Seq and ATAC-Seq Analysis of Human Developmental Hematopoiesis. Cell Stem Cell 28, 472–487.e7 (2021).

73. Massri, A. J. et al. Developmental single-cell transcriptomics in the Lytechinus variegatus sea urchin embryo. Development 148, dev198614 (2021).

74. McFaline-Figueroa, J. L. et al. A pooled single-cell genetic screen identifies regulatory checkpoints in the continuum of the epithelial-to-mesenchymal transition. Nat Genet 51, 1389–1398 (2019).

75. Leinonen, R., Sugawara, H., Shumway, M., & on behalf of the International Nucleotide Sequence Database Collaboration. The Sequence Read Archive. Nucleic Acids Research 39, D19–D21 (2011).

76. Barrett, T. et al. NCBI GEO: archive for functional genomics data sets—update. Nucleic Acids Research 41, D991–D995 (2013).

77. Stuart, T. et al. Comprehensive Integration of Single-Cell Data. Cell 177, 1888–1902.e21 (2019).

78. Jacomy, M., Venturini, T., Heymann, S. & Bastian, M. ForceAtlas2, a Continuous Graph Layout Algorithm for Handy Network Visualization Designed for the Gephi Software. PLOS ONE 9, e98679 (2014).

79. Rousseeuw, P. J. Silhouettes: A graphical aid to the interpretation and validation of cluster analysis. Journal of Computational and Applied Mathematics 20, 53–65 (1987).

80. Subramanian, A. et al. Gene set enrichment analysis: A knowledge-based approach for interpreting genome-wide expression profiles. Proceedings of the National Academy of Sciences 102, 15545–15550 (2005).

81. Liberzon, A. et al. Molecular signatures database (MSigDB) 3.0. Bioinformatics 27, 1739– 1740 (2011).

82. Fang, Z., Liu, X. & Peltz, G. GSEApy: a comprehensive package for performing gene set enrichment analysis in Python. Bioinformatics 39, btac757 (2023).

83. Waskom, M. L. seaborn: statistical data visualization. Journal of Open Source Software 6, 3021 (2021).

84. McKinney, W. Data Structures for Statistical Computing in Python. in 56–61 (Austin, Texas, 2010). doi:10.25080/Majora-92bf1922-00a.

85. Virtanen, P. et al. SciPy 1.0: fundamental algorithms for scientific computing in Python. Nat Methods 17, 261–272 (2020).

