## Supplementary data for "Reconstruction of single cell lineage trajectories and identification of diversity in fates during the epithelial-to-mesenchymal transition"

### **Supplementary Appendix**

#### Appendix S1. Threshold Stability Analysis for Top Ancestor Designation

To verify the stability of our findings relative to the threshold selected for designating top ancestors, we modified our initial 75% transitioning probability to 90%, testing all thresholds in 3% increments from 75% to 90%. For clarity, we showcase only the results for the 90% threshold in this section. Results from other tested percentages fall between the 75% and 90% benchmarks.

1) The timeline of cell commitment: A substantial fraction of the undefined ancestors were evident at day 0. Yet, as the TGF-beta treatment progressed, their prevalence dropped dramatically from 91.62% on day 0 to 15.92% by day 4, when using the 75% threshold. With a heightened threshold of 90%, the decline is seen from 98.83% to 24.85%, which doesn't alter our conclusion. Primary ancestors corresponding to the low EMT fate emerged between days 0 and 1. Those leaning towards the other EMT fates started to appear by day 2, implying varied commitment schedules (Fig 2B). These patterns are further substantiated by our triangular plots (see Fig 2C and Methods). Before the initiation of treatment, top ancestors for the low EMT fate were double that of the combined count for the other two fates at the 75% threshold (5.52% vs 2.85% on day 0). Adjusting the threshold to 90% results in proportions of 1.06% in low EMT and 0.11% in the other fates. By the treatment's second day, the percentages of top ancestors for all three fates were closely matched: 13.57%, 12.35%, and 13.31% for the low, medium, and high EMT trajectories at the 75% threshold. This underscores a more noticeable delay in tendencies towards the medium and high EMT fates. At the 90% threshold, the distributions were 8.96%, 6.85%, and 4.50% on day 3, highlighting a pronounced delay for the medium and high EMT fates.

2) Trajectory heterogeneity: Beyond evaluating pairwise cell state distances, we further investigated cellular EMT heterogeneity by examining the distribution of EMT scores among the top ancestors for the three pinpointed EMT fates (Fig 4B and Methods). In line with our previous observations from the pairwise distances, top ancestors from the partial EMT trajectory displayed a markedly diverse EMT score compared to those aligned with the high EMT fate at the 75% threshold. This distinction was statistically significant as indicated by the Levene test (p-values < 1e-5 from days 1 to 8, Table S4). When adjusting the threshold to 90%, the emergence of this significant variation appeared slightly later (p-values < 0.05 from days 2 to 8, Table S4). When analyzing stemness scores, the partial EMT trajectory displayed subtler fluctuations in comparison to its EMT counterparts, especially when contrasted with the high EMT. On days 1, 4, and 8, with the 75% threshold, p-values exceeded 0.05. On the days showcasing significant deviations, the high EMT trajectory revealed higher variations (Fig 4B, Fig S10, and Table S6). Adjusting to the 90% threshold, the findings remained consistent (Table S6).

3) Correlations between EMT and stemness signatures: To further describe the relationship between EMT and stemness signatures across various trajectories, we utilized 2D scatter plots to study the concurrent distributions of these markers over different time intervals (Methods). When identifying top ancestor sets with a 75% threshold, the low EMT subset consistently exhibited a positive correlation between EMT and stemness signatures from days 1 to 8, as evidenced by correlation coefficients spanning 0.22 to 0.44. In contrast, the high EMT subset displayed a negative trend, with coefficients oscillating between -0.3 and -0.6 (Table S8). When we shifted to a 90% threshold, we noticed analogous trends from days 3 to 8. The emergence of these trends was slightly delayed compared to the 75% threshold, attributable to the sparse cell count on days 1 and 2 (specifically, only 11 cells on day 1 and 23 cells on day 2 in the high EMT trajectory). For the low EMT subset, the correlation coefficients ranged between 0.29 and 0.46. In contrast, the high EMT subset demonstrated coefficients between -0.07 and -0.6.

### Supplementary figures

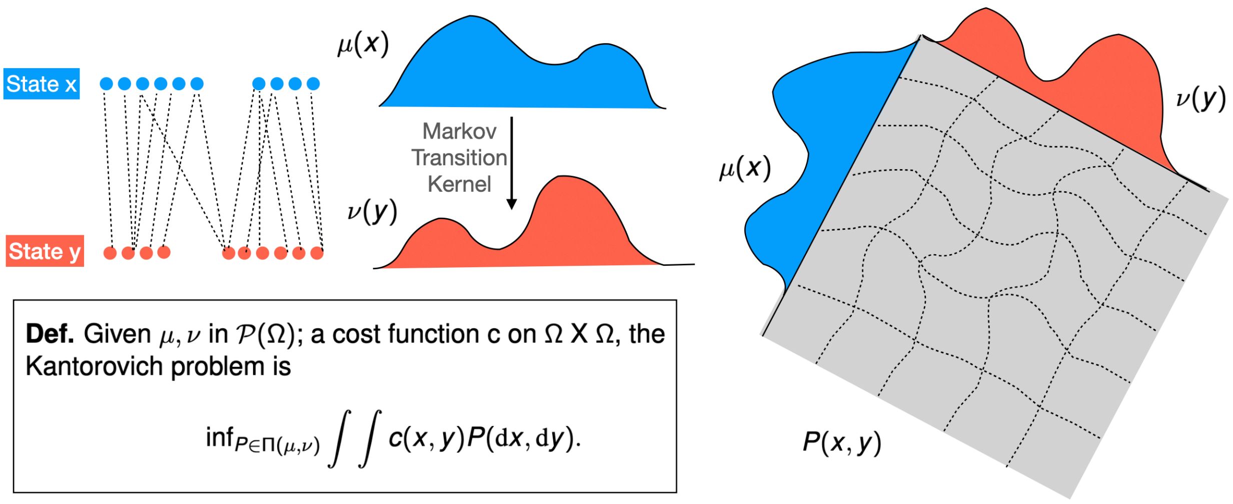


*Figure S1* Given two marginal distributions μ(x) and ν(x), optimal transport can estimate the unique joint distribution P(x, y) that best aligns with both marginal distributions. This is achieved by solving the Kantorovich problem, which aims to minimize the total distance between the two distributions while ensuring mass is conserved[1–3].

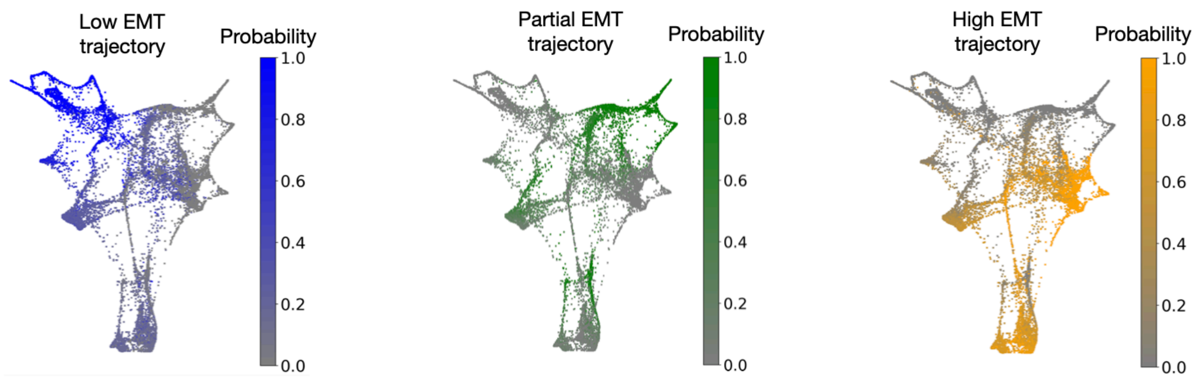


*Figure S2* FLE plots of cells from all time points, with the color map highlighting ATF distributions. These plots discern three distinct trajectories that define the three fates under investigation, separated by color.

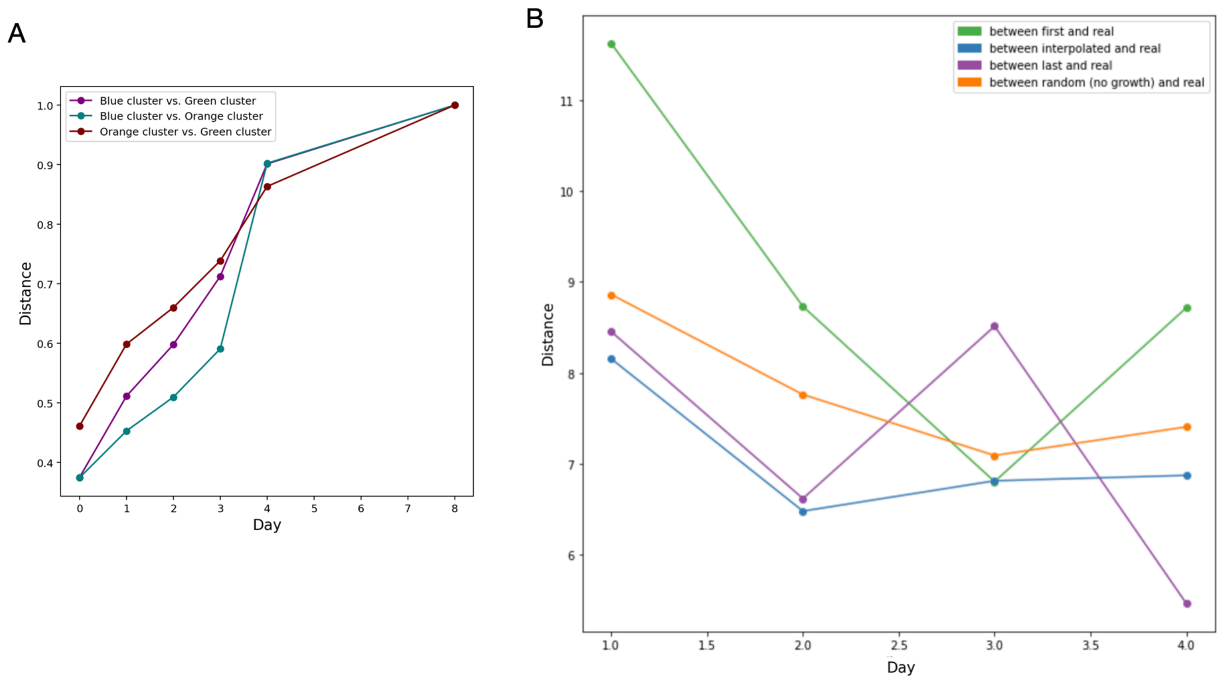


*Figure S3* (A) Wasserstein distance calculations between ATF distributions of fate clusters over time highlight how trajectories progressively diverge at each time point. (B) Wasserstein distance between the distribution of our estimate of ancestor distributions (Methods) and the real data (blue curve) compared with the other distances, including the Wasserstein distance between the test data and its preceding day's actual data (green), the Wasserstein distance between the test data and its subsequent day's actual data (purple), and the Wasserstein distance between the test data and the estimated data from the interpolation based on a random map (orange).

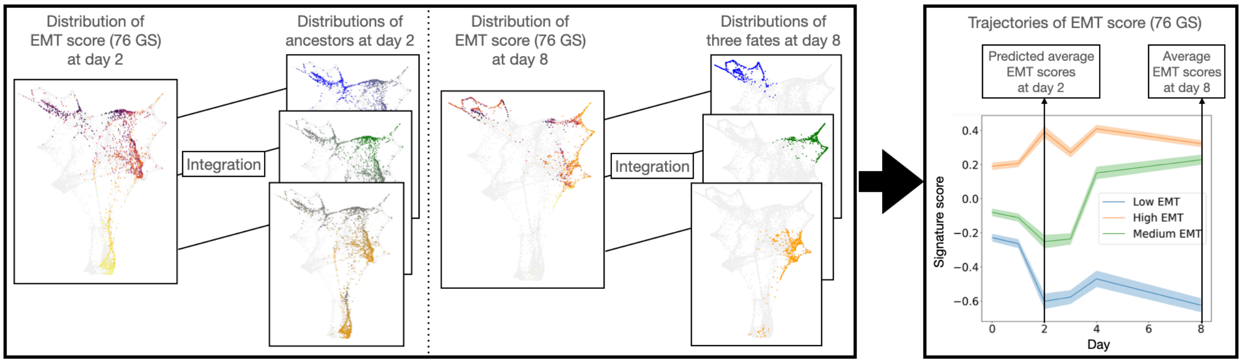


*Figure S4* The schematic illustrates the steps to infer the dynamic average EMT score along the trajectories. The EMT score calculated through the 76 GS method is integrated with the ATF distributions corresponding to each fate at specific early time points. Average expressions associated with each fate is computed.

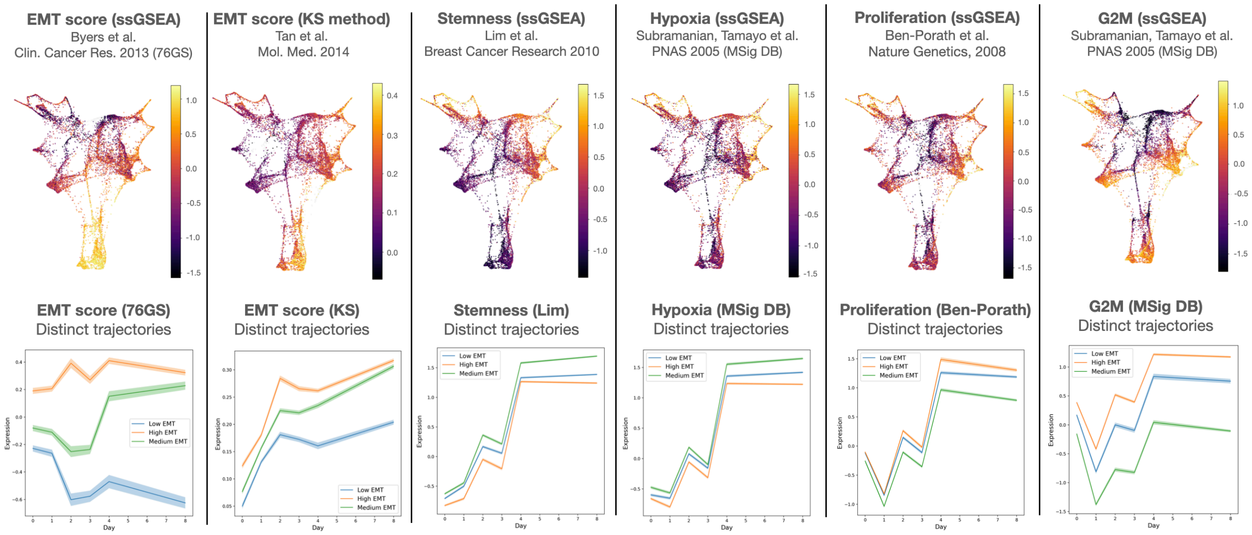


*Figure S5* Upper row: Color maps from left to right depict EMT signatures (by 76GS method and by KS method), stemness signature, hypoxia hallmark, proliferation signature, and G2M hallmark scores for each cellular state gathered from day 0 to day 8. Lower row: The averaged cellular signature scores are displayed across the three distinct EMT trajectories.

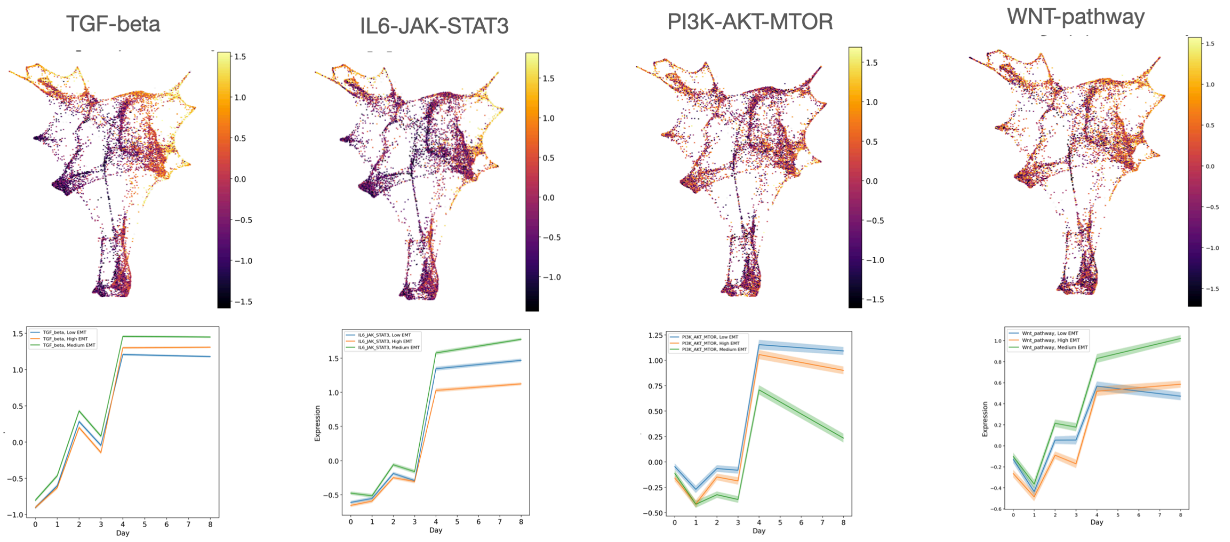


*Figure S6* Upper row: Color maps from left to right TGF-beta pathway, IL6-JAK-STAT3 pathway, PI3-AKT-mTOR pathway, and wnt pathway hallmark scores for each cellular state gathered from day 0 to day 8. Lower row: The averaged signature scores are displayed across the three distinct EMT trajectories.

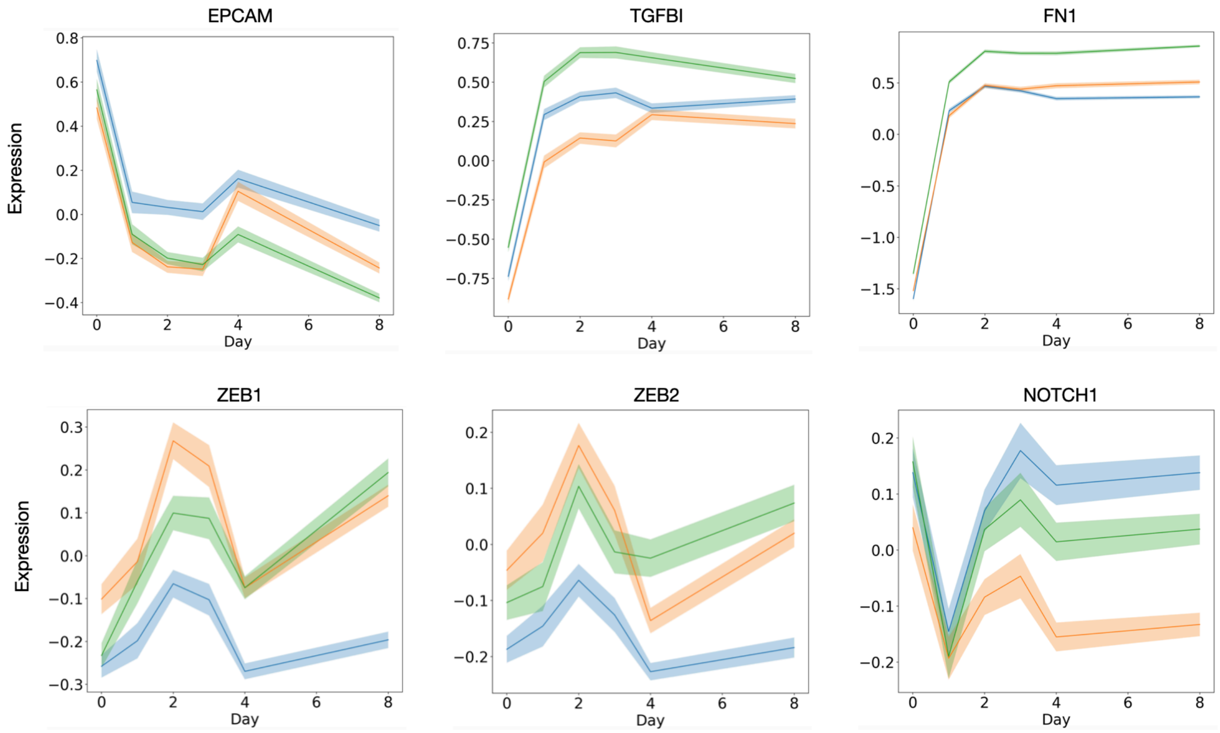


*Figure S7* Distinct gene expression across the EMT trajectories leading to the three different fates are shown.

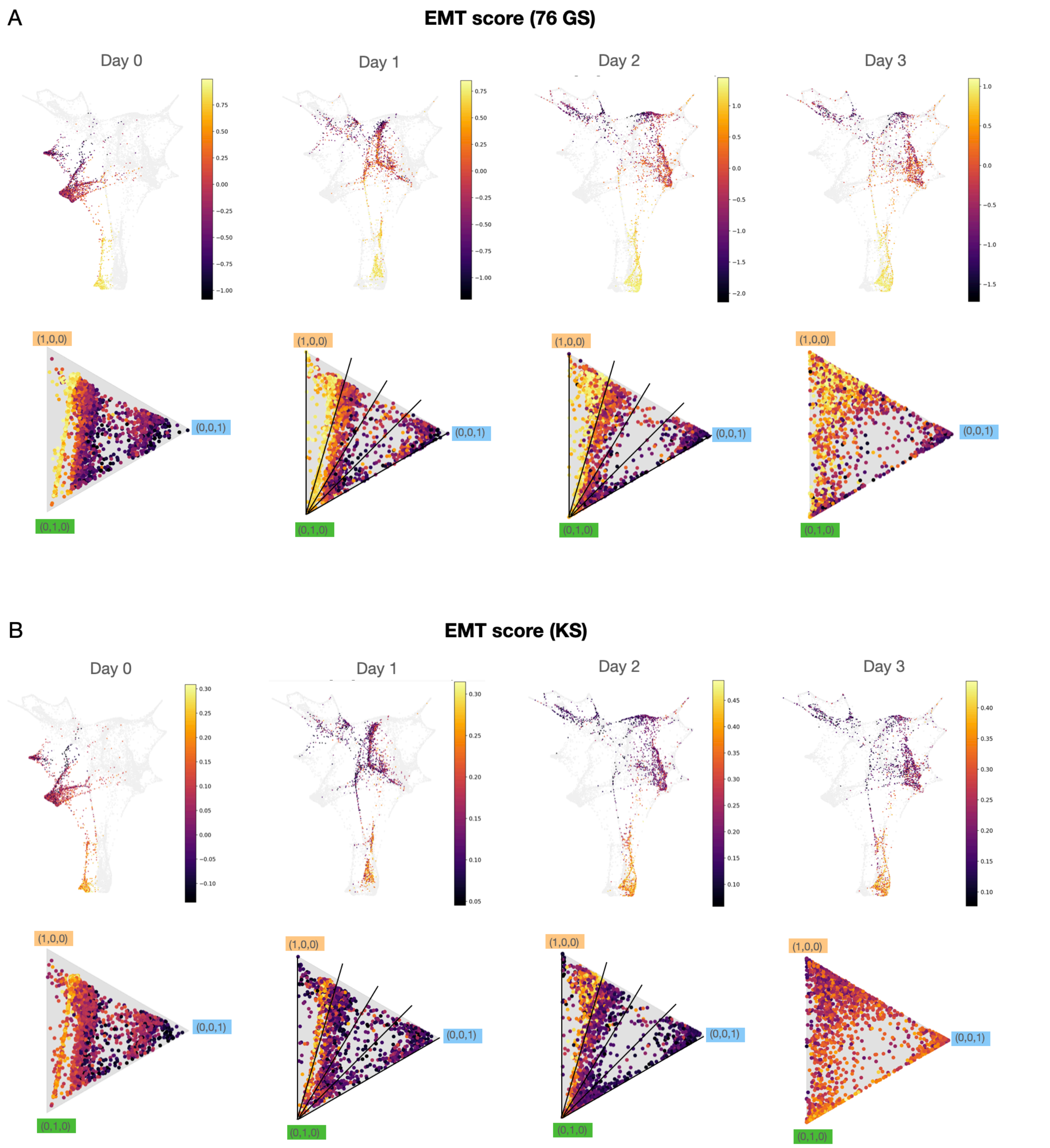


*Figure S8* A time-series of cellular EMT score signatures plotted against their potential differentiation fate in corresponding time-series barycentric plots. **(A)** EMT score computed by 76GS method and **(B)** EMT score computed by KS method. Upper row: a time series of cellular EMT score signature. Lower row: time series of cellular EMT score signature in (A) the corresponding time-series triangle plot, where the position represents the probability of differentiating into each fate, and the color map represents the EMT score. The five solid auxiliary lines depict five fixed ratios (0, 0.25, 0.50, 0.75, 1) between the likelihoods of transitioning into a complete EMT fate (symbolized by orange and vector 1,0,0) and a low EMT fate (represented by blue and vector 0,0,1).

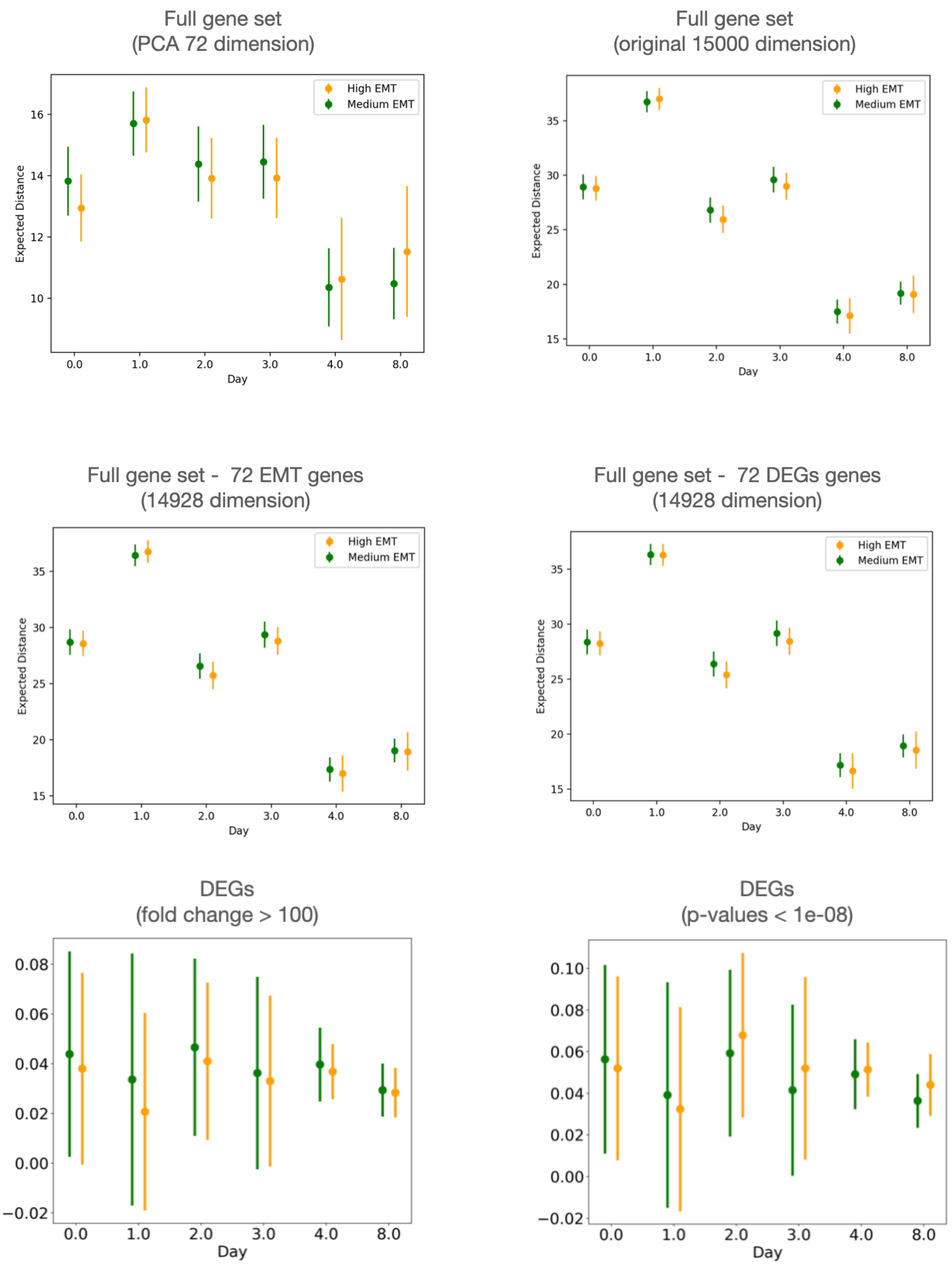


*Figure S9* Mean pairwise distances in cellular transcriptomics (indicated at the center of each bar), and the standard errors of these pairwise distances (symbolized by the length of the error bars). Upper row: the full gene expression space is presented before and after PCA. Middle row: the full gene expression space is portrayed after excluding the EMT gene set and the DEG set. Lower row: the gene expression space determined by DEGs based on a fold change $> 100$, then sorted by p-values (left panel), or based on p-values $< 1e-8$, followed by sorting the top genes by fold changes (right panel).

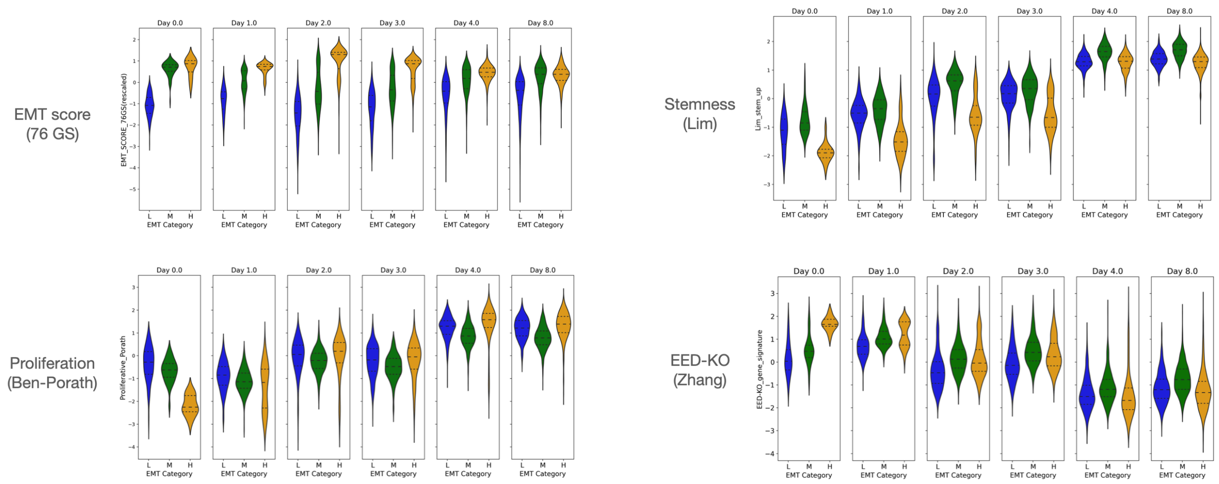


*Figure S10* Violin plots depicting the distribution of each cellular signature score for the top ancestors of each fate, color-coded as blue for low (L), green for medium (M), and orange for high (H) EMT trajectories.

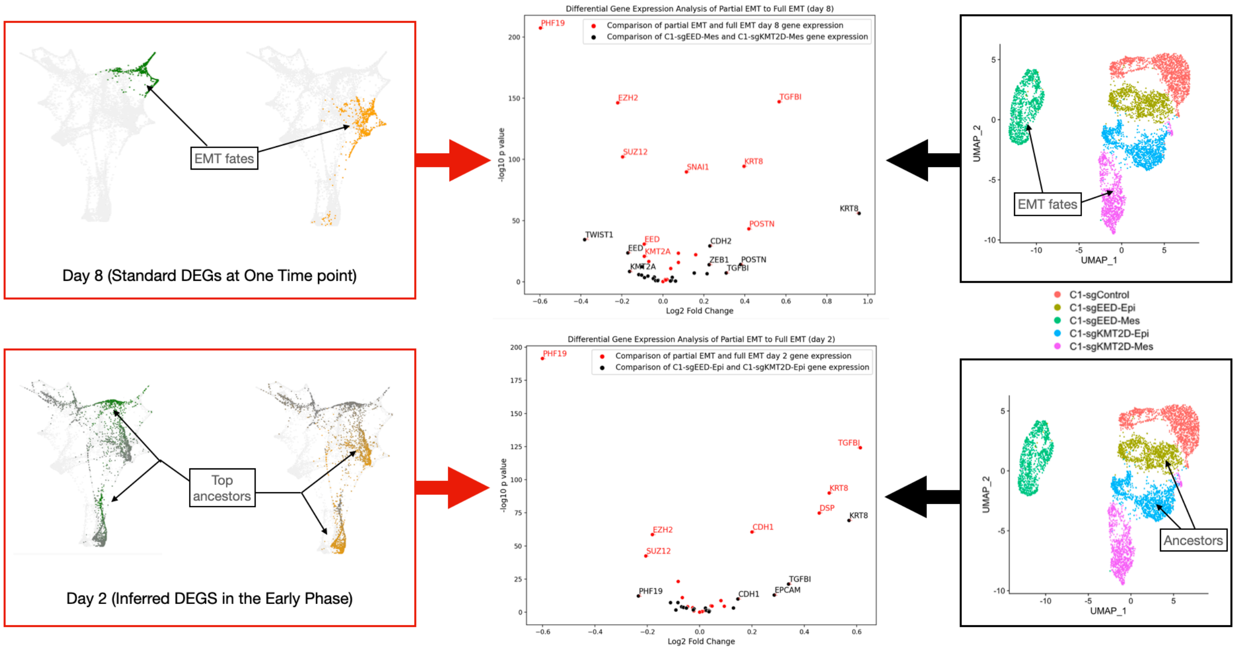


*Figure S11* Comparison of gene expression profiles between independent datasets. Left panel: Standard differential gene expression analysis at day 8 and early phase differential gene expression analysis at day 2 were performed, using the MCF10A dataset (Deshmukh et al). Right panel: Standard differential gene expression analysis was conducted separately for C1-sgEED-Epi/C1-sgKMT2D-Epi and C1-sgEED-Mes/C1-sgKMT2D-Mes (Zhang et al.). The results are displayed as black dots in the middle panel's volcano plot. Middle panel: volcano plots representing the results of the differential gene expression analysis conducted on the two datasets.

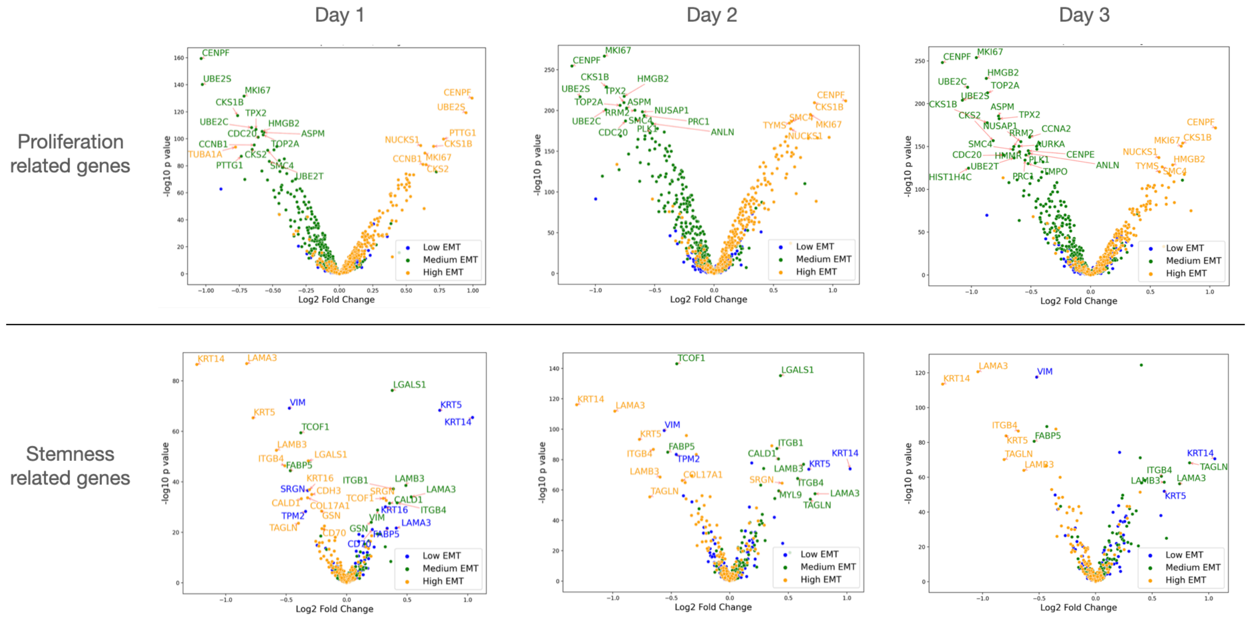


*Figure S12* Early DEGs of proliferation and stemness-related genes. Comparisons were conducted between early phase cellular states of one fate against the combined early phase cellular states of the other two fates. The distinct colors represent the differential gene expression associated with each of the three fates. The data reflect results from multiple time points, from day 1 to day 3.

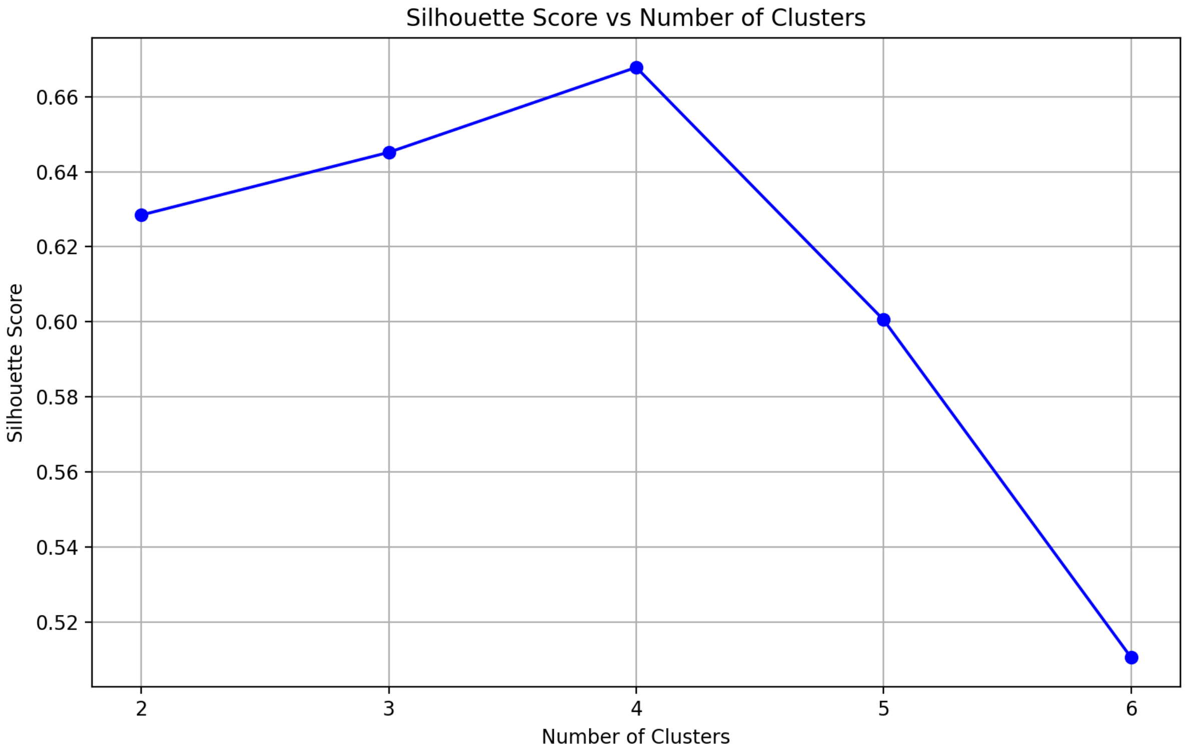


*Figure S13* Silhouette scores for different numbers of clusters (k) in K-means clustering.

### Supplementary tables

Differential expressed genes - 36 genes (fold change > 100)
(High expression in the partial EMT fate)

| | Gene | T-statistic | P-value | Fold Change |
|---|---|---|---|---|
| 5602 | AC007064.24 | 16.948708 | 2.273743e-58 | 105.042133 |
| 7043 | NR5A2 | -15.225729 | 2.695803e-48 | 649.147879 |
| 9749 | CDKL4 | 13.343261 | 3.704455e-38 | 1419.233104 |
| 5713 | AC011747.4 | 13.124216 | 4.835404e-37 | 136.608825 |
| 4732 | RASGRP2 | 12.279823 | 7.130148e-33 | 103.439527 |
| 2839 | CLDN6 | 11.848833 | 7.899196e-31 | 1087.517552 |
| 4118 | RP5-1185I7.1 | 11.604682 | 1.071768e-29 | 2688.833361 |
| 13156 | RP11-468E2.6 | 11.496002 | 3.374214e-29 | 215.891216 |
| 4372 | ALS2CR11 | -11.132874 | 1.462167e-27 | 113.049655 |
| 5070 | RP11-439C8.2 | -10.715100 | 9.891660e-26 | 126.042155 |
| 3288 | VAX2 | 10.412863 | 1.920809e-24 | 315.465631 |
| 13201 | RP11-1078H9.5 | 10.391302 | 2.367119e-24 | 964.035491 |
| 5660 | SLC2A14 | -10.336540 | 4.017615e-24 | 101.504973 |
| 6363 | SIDT1 | -10.057024 | 5.766344e-23 | 113.438466 |
| 5172 | RP11-624M8.1 | 9.918170 | 2.117174e-22 | 1746.370614 |
| 2151 | GJA1 | 9.541110 | 6.702070e-21 | 325.257688 |
| 862 | GPR37L1 | 9.193047 | 1.470179e-19 | 162.584465 |
| 12438 | PCDHB8 | 9.042691 | 5.414212e-19 | 141.936058 |
| 4816 | CTB-12A17.3 | 9.042691 | 5.414212e-19 | 141.936058 |
| 6848 | RP11-1105G2.4 | -9.020182 | 6.570623e-19 | 101.475869 |
| 6501 | CPEB2-AS1 | 8.536992 | 3.789582e-17 | 110.222963 |
| 9814 | CREB3L1 | 8.325443 | 2.104192e-16 | 1685.128318 |
| 3972 | LINC00565 | 8.134983 | 9.537010e-16 | 301.626479 |
| 2912 | CCDC116 | 8.016841 | 2.397981e-15 | 138.568936 |
| 4595 | SCGB2A1 | 7.596258 | 5.801184e-14 | 251.805709 |
| 4790 | RP4-734P14.4 | -7.453211 | 1.656227e-13 | 146.100676 |
| 5365 | SLC26A10 | -7.190251 | 1.087930e-12 | 145.509095 |
| 2621 | AC012363.4 | -7.087557 | 2.232373e-12 | 157.089735 |
| 5892 | RP11-482M8.1 | -7.087557 | 2.232373e-12 | 157.089735 |
| 3182 | RP1-90J4.1 | 6.931461 | 6.540458e-12 | 102.034410 |
| 14622 | SSBP3-AS1 | 6.806893 | 1.518815e-11 | 217.058580 |
| 13519 | GDF5 | 6.493131 | 1.193515e-10 | 1084.484418 |
| 2458 | TNFRSF14 | 6.460892 | 1.467841e-10 | 383.251081 |
| 3684 | TLL2 | 5.992200 | 2.676688e-09 | 145.412695 |
| 13348 | RP11-77K12.4 | 5.978627 | 2.902975e-09 | 145.869658 |
| 2590 | RP11-166B2.8 | 5.751992 | 1.098373e-08 | 556.383024 |

Differential expressed genes - 36 genes (fold change > 100)
(High expression in the complete EMT fate)

| | Gene | T-statistic | P-value | Fold Change |
|---|---|---|---|---|
| 7509 | RP11-138C9.1 | -11.678123 | 4.913674e-30 | 446.627364 |
| 9811 | RP11-84A19.4 | -11.015190 | 4.856718e-27 | 164.329867 |
| 6904 | TNFRSF13C | -9.944134 | 1.662023e-22 | 18515.697160 |
| 8156 | CARNS1 | -9.696440 | 1.636984e-21 | 139.687058 |
| 10001 | RP11-746M1.1 | -9.324751 | 4.622569e-20 | 142.451844 |
| 5397 | RP11-84C10.4 | -9.081948 | 3.859103e-19 | 515.398756 |
| 12789 | AL358113.1 | -9.081948 | 3.859103e-19 | 515.398756 |
| 13623 | CTD-2583A14.10 | -8.776412 | 5.205831e-18 | 578.915494 |
| 5993 | LBX2 | -8.723185 | 8.127058e-18 | 576.004023 |
| 14452 | FAM90A1 | 8.613316 | 2.023067e-17 | 990.375158 |
| 4661 | DCHS1 | -8.590324 | 2.445361e-17 | 348.861696 |
| 4468 | CR2 | 8.452886 | 7.525397e-17 | 979.151881 |
| 14931 | AC092415.1 | -8.419453 | 9.868411e-17 | 493.639030 |
| 14145 | AC006004.1 | -7.778207 | 1.489292e-14 | 106.154697 |
| 8378 | AC005592.3 | -7.653083 | 3.805478e-14 | 1239.385380 |
| 5192 | NEURL1 | -7.643549 | 4.085210e-14 | 104.012704 |
| 6250 | RP11-231N3.1 | -7.422878 | 2.064171e-13 | 785.865599 |
| 6980 | SORCS1 | -7.236926 | 7.823454e-13 | 213.602628 |
| 4865 | RP5-902P8.10 | -6.963397 | 5.258333e-12 | 108.854000 |
| 7273 | RP11-11C20.3 | -6.428336 | 1.807214e-10 | 200.527670 |
| 14989 | PPAPDC3 | -6.414575 | 1.972725e-10 | 130.692960 |
| 13675 | RP11-304F15.7 | 6.334556 | 3.272852e-10 | 395.626387 |
| 5541 | RP11-839G9.1 | 6.332493 | 3.315612e-10 | 350.751055 |
| 5328 | GLIPR1L1 | -6.243161 | 5.794501e-10 | 155.957976 |
| 6206 | HBA2 | -6.242349 | 5.823820e-10 | 155.906556 |
| 7452 | RP11-665N17.4 | -5.986121 | 2.775821e-09 | 146.289510 |
| 9406 | RP11-318E3.9 | -5.969828 | 3.059526e-09 | 135.596757 |
| 6998 | SERINC4 | -5.868489 | 5.574320e-09 | 330.257995 |
| 10003 | VIL1 | -5.838981 | 6.626776e-09 | 504.248161 |
| 7009 | CLDN9 | -5.836436 | 6.726138e-09 | 129.112848 |
| 14276 | RP11-771K4.1 | -5.631938 | 2.181462e-08 | 281.209558 |
| 3348 | AC073257.2 | -5.631938 | 2.181462e-08 | 281.209558 |
| 3559 | CTC-558O2.1 | 5.538356 | 3.690822e-08 | 102.152021 |
| 4195 | RP11-583F2.5 | 5.538356 | 3.690822e-08 | 102.152021 |
| 14202 | RP11-384P7.6 | -5.506988 | 4.394453e-08 | 187.198390 |
| 12288 | C4orf6 | -5.506988 | 4.394453e-08 | 187.198390 |

Differential expressed genes - 36 genes (p-values < 1e-7)
(High expression in the partial EMT fate)

| | Gene | T-statistic | P-value | Fold Change |
|---|---|---|---|---|
| 4118 | RP5-1185I7.1 | 11.604682 | 1.071768e-29 | 2688.833361 |
| 5172 | RP11-624M8.1 | 9.918170 | 2.117174e-22 | 1746.370614 |
| 9814 | CREB3L1 | 8.325443 | 2.104192e-16 | 1685.128318 |
| 9749 | CDKL4 | 13.343261 | 3.704455e-38 | 1419.233104 |
| 2839 | CLDN6 | 11.848833 | 7.899196e-31 | 1087.517552 |
| 13519 | GDF5 | 6.493131 | 1.193515e-10 | 1084.484418 |
| 13201 | RP11-1078H9.5 | 10.391302 | 2.367119e-24 | 964.035491 |
| 7043 | NR5A2 | -15.225729 | 2.695803e-48 | 649.147879 |
| 2590 | RP11-166B2.8 | 5.751992 | 1.098373e-08 | 556.383024 |
| 2458 | TNFRSF14 | 6.460892 | 1.467841e-10 | 383.251081 |
| 2151 | GJA1 | 9.541110 | 6.702070e-21 | 325.257688 |
| 3288 | VAX2 | 10.412863 | 1.920809e-24 | 315.465631 |
| 3972 | LINC00565 | 8.134983 | 9.537010e-16 | 301.626479 |
| 4595 | SCGB2A1 | 7.596258 | 5.801184e-14 | 251.805709 |
| 14622 | SSBP3-AS1 | 6.806893 | 1.518815e-11 | 217.058580 |
| 13156 | RP11-468E2.6 | 11.496002 | 3.374214e-29 | 215.891216 |
| 862 | GPR37L1 | 9.193047 | 1.470179e-19 | 162.584465 |
| 2621 | AC012363.4 | -7.087557 | 2.232373e-12 | 157.089735 |
| 5892 | RP11-482M8.1 | -7.087557 | 2.232373e-12 | 157.089735 |
| 5905 | RP11-114G22.1 | -5.395712 | 8.102382e-08 | 148.872771 |
| 4790 | RP4-734P14.4 | -7.453211 | 1.656227e-13 | 146.100676 |
| 13348 | RP11-77K12.4 | 5.978627 | 2.902975e-09 | 145.869658 |
| 5365 | SLC26A10 | -7.190251 | 1.087930e-12 | 145.509095 |
| 3684 | TLL2 | 5.992200 | 2.676688e-09 | 145.412695 |
| 12438 | PCDHB8 | 9.042691 | 5.414212e-19 | 141.936058 |
| 4816 | CTB-12A17.3 | 9.042691 | 5.414212e-19 | 141.936058 |
| 2912 | CCDC116 | 8.016841 | 2.397981e-15 | 138.568936 |
| 5713 | AC011747.4 | 13.124216 | 4.835404e-37 | 136.608825 |
| 5070 | RP11-439C8.2 | -10.715100 | 9.891660e-26 | 126.042155 |
| 6363 | SIDT1 | -10.057024 | 5.766344e-23 | 113.438466 |
| 4372 | ALS2CR11 | -11.132874 | 1.462167e-27 | 113.049655 |
| 6501 | CPEB2-AS1 | 8.536992 | 3.789582e-17 | 110.222963 |
| 5602 | AC007064.24 | 16.948708 | 2.273743e-58 | 105.042133 |
| 4732 | RASGRP2 | 12.279823 | 7.130148e-33 | 103.439527 |
| 3182 | RP1-90J4.1 | 6.931461 | 6.540458e-12 | 102.034410 |
| 5660 | SLC2A14 | -10.336540 | 4.017615e-24 | 101.504973 |

Differential expressed genes - 36 genes (p-values < 1e-7)
(High expression in the complete EMT fate)

| | Gene | T-statistic | P-value | Fold Change |
|---|---|---|---|---|
| 6904 | TNFRSF13C | -9.944134 | 1.662023e-22 | 18515.697160 |
| 8378 | AC005592.3 | -7.653083 | 3.805478e-14 | 1239.385380 |
| 14452 | FAM90A1 | 8.613316 | 2.023067e-17 | 990.375158 |
| 4468 | CR2 | 8.452886 | 7.525397e-17 | 979.151881 |
| 6250 | RP11-231N3.1 | -7.422878 | 2.064171e-13 | 785.865599 |
| 13623 | CTD-2583A14.10 | -8.776412 | 5.205831e-18 | 578.915494 |
| 5993 | LBX2 | -8.723185 | 8.127058e-18 | 576.004023 |
| 5397 | RP11-84C10.4 | -9.081948 | 3.859103e-19 | 515.398756 |
| 12789 | AL358113.1 | -9.081948 | 3.859103e-19 | 515.398756 |
| 10003 | VIL1 | -5.838981 | 6.626776e-09 | 504.248161 |
| 14931 | AC092415.1 | -8.419453 | 9.868411e-17 | 493.639030 |
| 7509 | RP11-138C9.1 | -11.678123 | 4.913674e-30 | 446.627364 |
| 13675 | RP11-304F15.7 | 6.334556 | 3.272852e-10 | 395.626387 |
| 5541 | RP11-839G9.1 | 6.332493 | 3.315612e-10 | 350.751055 |
| 4661 | DCHS1 | -8.590324 | 2.445361e-17 | 348.861696 |
| 6998 | SERINC4 | -5.868489 | 5.574320e-09 | 330.257995 |
| 14276 | RP11-771K4.1 | -5.631938 | 2.181462e-08 | 281.209558 |
| 3348 | AC073257.2 | -5.631938 | 2.181462e-08 | 281.209558 |
| 6980 | SORCS1 | -7.236926 | 7.823454e-13 | 213.602628 |
| 7273 | RP11-11C20.3 | -6.428336 | 1.807214e-10 | 200.527670 |
| 12288 | C4orf6 | -5.506988 | 4.394453e-08 | 187.198390 |
| 14202 | RP11-384P7.6 | -5.506988 | 4.394453e-08 | 187.198390 |
| 9811 | RP11-84A19.4 | -11.015190 | 4.856718e-27 | 164.329867 |
| 5328 | GLIPR1L1 | -6.243161 | 5.794501e-10 | 155.957976 |
| 6206 | HBA2 | -6.242349 | 5.823820e-10 | 155.906556 |
| 7452 | RP11-665N17.4 | -5.986121 | 2.775821e-09 | 146.289510 |
| 10001 | RP11-746M1.1 | -9.324751 | 4.622569e-20 | 142.451844 |
| 8156 | CARNS1 | -9.696440 | 1.636984e-21 | 139.687058 |
| 9406 | RP11-318E3.9 | -5.969828 | 3.059526e-09 | 135.596757 |
| 14774 | AC090673.2 | -5.399577 | 7.933494e-08 | 130.860823 |
| 14989 | PPAPDC3 | -6.414575 | 1.972725e-10 | 130.692960 |
| 7009 | CLDN9 | -5.836436 | 6.726138e-09 | 129.112848 |
| 4865 | RP5-902P8.10 | -6.963397 | 5.258333e-12 | 108.854000 |
| 14145 | AC006004.1 | -7.778207 | 1.489292e-14 | 106.154697 |
| 5192 | NEURL1 | -7.643549 | 4.085210e-14 | 104.012704 |
| 10188 | PROCA1 | -5.476950 | 5.189300e-08 | 103.307237 |

*Table S1* Compilation of the top 31 differentially expressed genes observed in the partial EMT fate (left columns) and the full EMT fate (right columns). The upper table of DEG set is based on a fold change > 100, then sorted by p-values, and the lower table of DEG set is based on p-values < 1e-8, followed by sorting the top genes by fold changes.

<table>
<tr><th></th><th colspan="6">Statistic of pairwise distances</th><th>Differential analysis between the High EMT and Medium EMT</th></tr>
<tr><td rowspan="13">Full gene set<br>(PCA)</td><td></td><td>Day</td><td>EMT State</td><td>Expected Distance</td><td>Variance</td><td>N_Samples</td><td rowspan="13">Day 0.0 - T-statistic: -8.498713622377151, P-value: 2.4444651835751492e-17<br>Day 1.0 - T-statistic: 1.1204591977429041, P-value: 0.26257657358976116<br>Day 2.0 - T-statistic: -3.685727419010319, P-value: 0.0002305880257408696<br>Day 3.0 - T-statistic: -3.9318055241698167, P-value: 8.564922510284918e-05<br>Day 4.0 - T-statistic: 1.128867177868205, P-value: 0.25907200767670446<br>Day 8.0 - T-statistic: 5.416793061041387, P-value: 6.44593186436604e-08</td></tr>
<tr><td>0</td><td>0.0</td><td>High EMT</td><td>12.942481</td><td>14.275918</td><td>2734</td></tr>
<tr><td>1</td><td>0.0</td><td>Medium EMT</td><td>13.824241</td><td>15.154270</td><td>2734</td></tr>
<tr><td>2</td><td>1.0</td><td>High EMT</td><td>15.821649</td><td>13.592416</td><td>2303</td></tr>
<tr><td>3</td><td>1.0</td><td>Medium EMT</td><td>15.700772</td><td>13.211000</td><td>2303</td></tr>
<tr><td>4</td><td>2.0</td><td>High EMT</td><td>13.914682</td><td>20.628944</td><td>2381</td></tr>
<tr><td>5</td><td>2.0</td><td>Medium EMT</td><td>14.384348</td><td>18.033624</td><td>2381</td></tr>
<tr><td>6</td><td>3.0</td><td>High EMT</td><td>13.927695</td><td>20.477954</td><td>2132</td></tr>
<tr><td>7</td><td>3.0</td><td>Medium EMT</td><td>14.451023</td><td>17.292291</td><td>2132</td></tr>
<tr><td>8</td><td>4.0</td><td>High EMT</td><td>10.634625</td><td>47.521079</td><td>1147</td></tr>
<tr><td>9</td><td>4.0</td><td>Medium EMT</td><td>10.361767</td><td>19.490338</td><td>1147</td></tr>
<tr><td>10</td><td>8.0</td><td>High EMT</td><td>11.527830</td><td>54.640249</td><td>1891</td></tr>
<tr><td>11</td><td>8.0</td><td>Medium EMT</td><td>10.478976</td><td>16.258290</td><td>1891</td></tr>
<tr><td rowspan="13">EMT gene set</td><td></td><td>Day</td><td>EMT State</td><td>Expected Distance</td><td>Variance</td><td>N_Samples</td><td rowspan="13">Day 0.0 - T-statistic: -7.886097058376222, P-value: 3.7371491597092114e-15<br>Day 1.0 - T-statistic: -16.255246742326758, P-value: 8.129003863958924e-58<br>Day 2.0 - T-statistic: -10.564884326916586, P-value: 8.352314870355229e-26<br>Day 3.0 - T-statistic: -11.665275425268007, P-value: 5.643232611245925e-31<br>Day 4.0 - T-statistic: -6.357353566469898, P-value: 2.4690995888194686e-10<br>Day 8.0 - T-statistic: -2.766807211043642, P-value: 0.005688353513441644</td></tr>
<tr><td>0</td><td>0.0</td><td>High EMT</td><td>3.501307</td><td>0.883344</td><td>2734</td></tr>
<tr><td>1</td><td>0.0</td><td>Medium EMT</td><td>3.698273</td><td>0.822179</td><td>2734</td></tr>
<tr><td>2</td><td>1.0</td><td>High EMT</td><td>4.035182</td><td>0.911161</td><td>2303</td></tr>
<tr><td>3</td><td>1.0</td><td>Medium EMT</td><td>4.476537</td><td>0.786624</td><td>2303</td></tr>
<tr><td>4</td><td>2.0</td><td>High EMT</td><td>3.219057</td><td>1.008076</td><td>2381</td></tr>
<tr><td>5</td><td>2.0</td><td>Medium EMT</td><td>3.518250</td><td>0.901478</td><td>2381</td></tr>
<tr><td>6</td><td>3.0</td><td>High EMT</td><td>3.252536</td><td>0.839384</td><td>2132</td></tr>
<tr><td>7</td><td>3.0</td><td>Medium EMT</td><td>3.576365</td><td>0.803589</td><td>2132</td></tr>
<tr><td>8</td><td>4.0</td><td>High EMT</td><td>2.151285</td><td>0.580462</td><td>1147</td></tr>
<tr><td>9</td><td>4.0</td><td>Medium EMT</td><td>2.352396</td><td>0.567374</td><td>1147</td></tr>
<tr><td>10</td><td>8.0</td><td>High EMT</td><td>2.213821</td><td>0.693097</td><td>1891</td></tr>
<tr><td>11</td><td>8.0</td><td>Medium EMT</td><td>2.284232</td><td>0.531548</td><td>1891</td></tr>
<tr><td rowspan="13">DEGs gene set<br>(f-change > 100)</td><td></td><td>Day</td><td>EMT State</td><td>Expected Distance</td><td>Variance</td><td>N_Samples</td><td rowspan="13">Day 0.0 - T-statistic: -0.290551836303597, P-value: 0.7714051232869783<br>Day 1.0 - T-statistic: 0.45072771279893803, P-value: 0.6522070315748654<br>Day 2.0 - T-statistic: 1.2868494309580503, P-value: 0.19820935025692266<br>Day 3.0 - T-statistic: 1.8960118589550194, P-value: 0.058026052290491015<br>Day 4.0 - T-statistic: 4.281495755515565, P-value: 1.9328821954292443e-05<br>Day 8.0 - T-statistic: 4.177583278804744, P-value: 3.012789339901324e-05</td></tr>
<tr><td>0</td><td>0.0</td><td>High EMT</td><td>0.053429</td><td>0.027381</td><td>2734</td></tr>
<tr><td>1</td><td>0.0</td><td>Medium EMT</td><td>0.054710</td><td>0.025747</td><td>2734</td></tr>
<tr><td>2</td><td>1.0</td><td>High EMT</td><td>0.043503</td><td>0.038041</td><td>2303</td></tr>
<tr><td>3</td><td>1.0</td><td>Medium EMT</td><td>0.040995</td><td>0.033253</td><td>2303</td></tr>
<tr><td>4</td><td>2.0</td><td>High EMT</td><td>0.050781</td><td>0.017051</td><td>2381</td></tr>
<tr><td>5</td><td>2.0</td><td>Medium EMT</td><td>0.045984</td><td>0.016032</td><td>2381</td></tr>
<tr><td>6</td><td>3.0</td><td>High EMT</td><td>0.044028</td><td>0.020221</td><td>2132</td></tr>
<tr><td>7</td><td>3.0</td><td>Medium EMT</td><td>0.036142</td><td>0.016655</td><td>2132</td></tr>
<tr><td>8</td><td>4.0</td><td>High EMT</td><td>0.069427</td><td>0.011817</td><td>1147</td></tr>
<tr><td>9</td><td>4.0</td><td>Medium EMT</td><td>0.053114</td><td>0.004833</td><td>1147</td></tr>
<tr><td>10</td><td>8.0</td><td>High EMT</td><td>0.073236</td><td>0.016215</td><td>1891</td></tr>
<tr><td>11</td><td>8.0</td><td>Medium EMT</td><td>0.057913</td><td>0.009225</td><td>1891</td></tr>
<tr><td rowspan="13">DEGs gene set<br>(p-value < 1e-07)</td><td></td><td>Day</td><td>EMT State</td><td>Expected Distance</td><td>Variance</td><td>N_Samples</td><td rowspan="13">Day 0.0 - T-statistic: -0.86955761517252, P-value: 0.384580390879379<br>Day 1.0 - T-statistic: -0.9515321873762885, P-value: 0.34138423607957247<br>Day 2.0 - T-statistic: -1.9912700188302463, P-value: 0.04650827272845152<br>Day 3.0 - T-statistic: -1.2912536550655862, P-value: 0.19668576329980175<br>Day 4.0 - T-statistic: -5.428236457024454, P-value: 6.291183031966002e-08<br>Day 8.0 - T-statistic: -0.5249631281797797, P-value: 0.5996396101376378</td></tr>
<tr><td>0</td><td>0.0</td><td>High EMT</td><td>0.054854</td><td>0.025549</td><td>2734</td></tr>
<tr><td>1</td><td>0.0</td><td>Medium EMT</td><td>0.058664</td><td>0.026942</td><td>2734</td></tr>
<tr><td>2</td><td>1.0</td><td>High EMT</td><td>0.039717</td><td>0.035047</td><td>2303</td></tr>
<tr><td>3</td><td>1.0</td><td>Medium EMT</td><td>0.045156</td><td>0.040191</td><td>2303</td></tr>
<tr><td>4</td><td>2.0</td><td>High EMT</td><td>0.063858</td><td>0.018145</td><td>2381</td></tr>
<tr><td>5</td><td>2.0</td><td>Medium EMT</td><td>0.072146</td><td>0.023106</td><td>2381</td></tr>
<tr><td>6</td><td>3.0</td><td>High EMT</td><td>0.049911</td><td>0.020778</td><td>2132</td></tr>
<tr><td>7</td><td>3.0</td><td>Medium EMT</td><td>0.055884</td><td>0.024849</td><td>2132</td></tr>
<tr><td>8</td><td>4.0</td><td>High EMT</td><td>0.047501</td><td>0.002003</td><td>1147</td></tr>
<tr><td>9</td><td>4.0</td><td>Medium EMT</td><td>0.059768</td><td>0.003855</td><td>1147</td></tr>
<tr><td>10</td><td>8.0</td><td>High EMT</td><td>0.042109</td><td>0.002338</td><td>1891</td></tr>
<tr><td>11</td><td>8.0</td><td>Medium EMT</td><td>0.042951</td><td>0.002521</td><td>1891</td></tr>
</table>

*Table S2* Detailed statistics of the pairwise distances, calculated using four gene sets (each row), are presented in the first main column. The second column includes the mean (expected distance), variance, and sample number. Additionally, a comparison between the means of high EMT and medium EMT is given in the right main column, featuring T-statistics and P-values, presented in the third main column.

| | Statistic of pairwise distances | Difference between the High EMT and Medium EMT |
|---|---|---|

**Full gene set (15000 genes)**

Statistic of pairwise distances:

| | Day | EMT State | Expected Distance | Variance | N_Samples |
|---|---|---|---|---|---|
| 0 | 0.0 | High EMT | 28.798462 | 15.049018 | 2734 |
| 1 | 0.0 | Medium EMT | 28.939525 | 15.515680 | 2734 |
| 2 | 1.0 | High EMT | 36.997907 | 12.116807 | 2303 |
| 3 | 1.0 | Medium EMT | 36.717035 | 11.300432 | 2303 |
| 4 | 2.0 | High EMT | 25.953145 | 18.750257 | 2381 |
| 5 | 2.0 | Medium EMT | 26.805133 | 15.874791 | 2381 |
| 6 | 3.0 | High EMT | 28.995012 | 18.579608 | 2132 |
| 7 | 3.0 | Medium EMT | 29.588970 | 16.336435 | 2132 |
| 8 | 4.0 | High EMT | 17.135363 | 31.802404 | 1147 |
| 9 | 4.0 | Medium EMT | 17.522630 | 14.558329 | 1147 |
| 10 | 8.0 | High EMT | 19.079738 | 35.375565 | 1891 |
| 11 | 8.0 | Medium EMT | 19.183305 | 13.345966 | 1891 |

Difference between the High EMT and Medium EMT:

Day 0.0 - T-statistic: -1.33414491639208, P-value: 0.1822119479560432
Day 1.0 - T-statistic: 2.785393059906075, P-value: 0.005368152661580604
Day 2.0 - T-statistic: -7.065090993203701, P-value: 1.8373293155558473e-12
Day 3.0 - T-statistic: -4.641268704771038, P-value: 3.5668159374019497e-06
Day 4.0 - T-statistic: -1.926273039378543, P-value: 0.05419382245520678
Day 8.0 - T-statistic: -0.6452201334680709, P-value: 0.5188236998410556

**Full gene set - 72 EMT genes (14928 genes)**

Statistic of pairwise distances:

| | Day | EMT State | Expected Distance | Variance | N_Samples |
|---|---|---|---|---|---|
| 0 | 0.0 | High EMT | 28.574234 | 14.843187 | 2734 |
| 1 | 0.0 | Medium EMT | 28.693944 | 15.274357 | 2734 |
| 2 | 1.0 | High EMT | 36.767697 | 11.991214 | 2303 |
| 3 | 1.0 | Medium EMT | 36.435824 | 11.194138 | 2303 |
| 4 | 2.0 | High EMT | 25.740676 | 18.403067 | 2381 |
| 5 | 2.0 | Medium EMT | 26.562695 | 15.598358 | 2381 |
| 6 | 3.0 | High EMT | 28.802436 | 18.328069 | 2132 |
| 7 | 3.0 | Medium EMT | 29.363225 | 16.126905 | 2132 |
| 8 | 4.0 | High EMT | 16.993416 | 31.466841 | 1147 |
| 9 | 4.0 | Medium EMT | 17.355507 | 14.318522 | 1147 |
| 10 | 8.0 | High EMT | 18.942524 | 35.015289 | 1891 |
| 11 | 8.0 | Medium EMT | 19.037069 | 13.207015 | 1891 |

Difference between the High EMT and Medium EMT:

Day 0.0 - T-statistic: -1.1405724167183335, P-value: 0.25409787275992474
Day 1.0 - T-statistic: 3.307594643067476, P-value: 0.0009482307168023021
Day 2.0 - T-statistic: -6.87880657564541, P-value: 6.816849782798876e-12
Day 3.0 - T-statistic: -4.411306783048043, P-value: 1.0528457161369128e-05
Day 4.0 - T-statistic: -1.8123250616102782, P-value: 0.07006678199586328
Day 8.0 - T-statistic: -0.592055644251013, P-value: 0.5538487371336432

**Full gene set - 72 DEGs genes (14928 genes)**

Statistic of pairwise distances:

| | Day | EMT State | Expected Distance | Variance | N_Samples |
|---|---|---|---|---|---|
| 0 | 0.0 | High EMT | 28.255901 | 14.504386 | 2734 |
| 1 | 0.0 | Medium EMT | 28.399692 | 15.055068 | 2734 |
| 2 | 1.0 | High EMT | 36.314138 | 11.807439 | 2303 |
| 3 | 1.0 | Medium EMT | 36.328877 | 10.939412 | 2303 |
| 4 | 2.0 | High EMT | 25.415645 | 18.114890 | 2381 |
| 5 | 2.0 | Medium EMT | 26.387443 | 15.497685 | 2381 |
| 6 | 3.0 | High EMT | 28.445679 | 17.939688 | 2132 |
| 7 | 3.0 | Medium EMT | 29.183589 | 15.915531 | 2132 |
| 8 | 4.0 | High EMT | 16.655490 | 30.989751 | 1147 |
| 9 | 4.0 | Medium EMT | 17.201246 | 14.123087 | 1147 |
| 10 | 8.0 | High EMT | 18.547412 | 34.491854 | 1891 |
| 11 | 8.0 | Medium EMT | 18.934259 | 13.039790 | 1891 |

Difference between the High EMT and Medium EMT:

Day 0.0 - T-statistic: -1.382872578539498, P-value: 0.1667604276812233
Day 1.0 - T-statistic: -0.14830029127263453, P-value: 0.8821122800182609
Day 2.0 - T-statistic: -8.179091316819186, P-value: 3.6349412218900536e-16
Day 3.0 - T-statistic: -5.855776159353483, P-value: 5.105196673313958e-09
Day 4.0 - T-statistic: -2.7518837940678917, P-value: 0.005971997253599919
Day 8.0 - T-statistic: -2.440022041775959, P-value: 0.014732012504938283

*Table S3* Detailed statistics of pairwise distances computed using three different gene sets (each row) are outlined in the first main column. All the three rows use the full gene set, but the second excludes the 72 EMT-related genes, and the third row excludes the 72 DEGs listed in Table S1. The second column statistics include mean (expected distance), variance, and sample number. Additionally, a comparison of the means between the high EMT and medium EMT, featuring T-statistics and P-values, is provided in the third main column.

EMT score, by 76GS method (75% top ancestors)

| | Day | Pair | Levene Statistic | Levene p-value | T-test Statistic | T-test p-value | Mean Group1 | Mean Group2 | Variance Group1 | Variance Group2 |
|---|---|---|---|---|---|---|---|---|---|---|
| 0 | 0.0 | Low EMT vs Medium EMT | 12.347598 | 0.000542 | -23.450441 | 0.0 | -1.072511 | 0.620413 | 0.259053 | 0.122936 |
| 1 | 0.0 | Low EMT vs High EMT | 1.41202 | 0.236398 | -14.743103 | 0.0 | -1.072511 | 0.726355 | 0.259053 | 0.186205 |
| 2 | 0.0 | Medium EMT vs High EMT | 0.916886 | 0.34133 | -1.081462 | 0.282911 | 0.620413 | 0.726355 | 0.122936 | 0.186205 |
| 3 | 1.0 | Medium EMT vs High EMT | 56.222071 | 0.0 | -10.607564 | 0.0 | 0.064054 | 0.635603 | 0.268057 | 0.090942 |
| 4 | 1.0 | Medium EMT vs Low EMT | 2.284196 | 0.131422 | 16.771802 | 0.0 | 0.064054 | -0.856927 | 0.268057 | 0.373864 |
| 5 | 1.0 | High EMT vs Low EMT | 51.622858 | 0.0 | 24.280633 | 0.0 | 0.635603 | -0.856927 | 0.090942 | 0.373864 |
| 6 | 2.0 | High EMT vs Low EMT | 28.784887 | 0.0 | 39.28534 | 0.0 | 0.970182 | -1.538226 | 0.494843 | 0.806734 |
| 7 | 2.0 | High EMT vs Medium EMT | 44.27853 | 0.0 | 19.030708 | 0.0 | 0.970182 | -0.305083 | 0.494843 | 0.889973 |
| 8 | 2.0 | Low EMT vs Medium EMT | 2.19799 | 0.138703 | -16.628777 | 0.0 | -1.538226 | -0.305083 | 0.806734 | 0.889973 |
| 9 | 3.0 | High EMT vs Medium EMT | 42.614207 | 0.0 | 17.989646 | 0.0 | 0.584925 | -0.310389 | 0.360403 | 0.682954 |
| 10 | 3.0 | High EMT vs Low EMT | 37.799059 | 0.0 | 36.6255 | 0.0 | 0.584925 | -1.24591 | 0.360403 | 0.711708 |
| 11 | 3.0 | Medium EMT vs Low EMT | 0.035301 | 0.851032 | 13.522366 | 0.0 | -0.310389 | -1.24591 | 0.682954 | 0.711708 |
| 12 | 4.0 | High EMT vs Low EMT | 100.8851 | 0.0 | 21.836769 | 0.0 | 0.434322 | -0.621325 | 0.146789 | 0.758056 |
| 13 | 4.0 | High EMT vs Medium EMT | 62.948551 | 0.0 | 8.383379 | 0.0 | 0.434322 | 0.085325 | 0.146789 | 0.423611 |
| 14 | 4.0 | Low EMT vs Medium EMT | 7.211371 | 0.007457 | -10.360736 | 0.0 | -0.621325 | 0.085325 | 0.758056 | 0.423611 |
| 15 | 8.0 | Medium EMT vs High EMT | 51.045856 | 0.0 | -3.079453 | 0.002117 | 0.228478 | 0.323103 | 0.432259 | 0.202847 |
| 16 | 8.0 | Medium EMT vs Low EMT | 26.294534 | 0.0 | 18.21756 | 0.0 | 0.228478 | -0.624904 | 0.432259 | 0.812004 |
| 17 | 8.0 | High EMT vs Low EMT | 143.906259 | 0.0 | 25.045698 | 0.0 | 0.323103 | -0.624904 | 0.202847 | 0.812004 |

EMT score, by 76GS method (90% top ancestors)

| | Day | Pair | Levene Statistic | Levene p-value | T-test Statistic | T-test p-value | Mean Group1 | Mean Group2 | Variance Group1 | Variance Group2 |
|---|---|---|---|---|---|---|---|---|---|---|
| 0 | 0.0 | Low EMT vs Medium EMT | 2.157547 | 0.152636 | -6.28564 | 0.000001 | -1.631174 | 0.373108 | 0.196975 | 0.001447 |
| 1 | 0.0 | Low EMT vs High EMT | 1.274742 | 0.26846 | NaN | NaN | -1.631174 | -0.126174 | 0.196975 | NaN |
| 2 | 0.0 | Medium EMT vs High EMT | inf | 0.0 | NaN | NaN | 0.373108 | -0.126174 | 0.001447 | NaN |
| 3 | 1.0 | Low EMT vs High EMT | 5.569171 | 0.021156 | -12.689715 | 0.0 | -1.485779 | 0.602346 | 0.280106 | 0.08251 |
| 4 | 1.0 | Low EMT vs Medium EMT | 16.345653 | 0.000125 | -16.291784 | 0.0 | -1.485779 | 0.526225 | 0.280106 | 0.022929 |
| 5 | 1.0 | High EMT vs Medium EMT | 1.252842 | 0.272521 | 0.955585 | 0.34746 | 0.602346 | 0.526225 | 0.08251 | 0.022929 |
| 6 | 2.0 | Low EMT vs Medium EMT | 24.702389 | 0.000002 | -19.76398 | 0.0 | -1.981631 | 0.708945 | 0.795766 | 0.170549 |
| 7 | 2.0 | Low EMT vs High EMT | 0.361011 | 0.54877 | -13.300185 | 0.0 | -1.981631 | 0.679536 | 0.795766 | 0.782281 |
| 8 | 2.0 | Medium EMT vs High EMT | 9.029101 | 0.003736 | 0.188956 | 0.850698 | 0.708945 | 0.679536 | 0.170549 | 0.782281 |
| 9 | 3.0 | Low EMT vs High EMT | 10.733293 | 0.001182 | -18.415598 | 0.0 | -1.414263 | 0.462706 | 0.791107 | 0.408855 |
| 10 | 3.0 | Low EMT vs Medium EMT | 3.674053 | 0.056116 | -13.644359 | 0.0 | -1.414263 | -0.172029 | 0.791107 | 0.548028 |
| 11 | 3.0 | High EMT vs Medium EMT | 3.947916 | 0.048067 | 6.880188 | 0.0 | 0.462706 | -0.172029 | 0.408855 | 0.548028 |
| 12 | 4.0 | High EMT vs Low EMT | 97.658156 | 0.0 | 22.711824 | 0.0 | 0.449949 | -0.771463 | 0.135711 | 0.78614 |
| 13 | 4.0 | High EMT vs Medium EMT | 52.979797 | 0.0 | 9.019407 | 0.0 | 0.449949 | 0.052257 | 0.135711 | 0.406247 |
| 14 | 4.0 | Low EMT vs Medium EMT | 9.578964 | 0.002091 | -10.899655 | 0.0 | -0.771463 | 0.052257 | 0.78614 | 0.406247 |
| 15 | 8.0 | Medium EMT vs High EMT | 51.045856 | 0.0 | -3.079453 | 0.002117 | 0.228478 | 0.323103 | 0.432259 | 0.202847 |
| 16 | 8.0 | Medium EMT vs Low EMT | 26.294534 | 0.0 | 18.21756 | 0.0 | 0.228478 | -0.624904 | 0.432259 | 0.812004 |
| 17 | 8.0 | High EMT vs Low EMT | 143.906259 | 0.0 | 25.045698 | 0.0 | 0.323103 | -0.624904 | 0.202847 | 0.812004 |

*Table S4* Statistics of the EMT signature score (76 GS method) calculated for the top ancestors across the three EMT trajectories for all time points, spanning from day 0 to day 8. The difference of variations among the trajectories were computed using the Levene test, while the difference of means were analyzed using the t-test. Additionally, the table displays the mean and variation for the top ancestors of each trajectory group on every day. In the context of these values, 'Group 1' corresponds to the first trajectory listed in the "EMT pair" column, while 'Group 2' signifies the second. Upper table: results determined by the 75% threshold for identifying the top ancestor set. Lower table: results based on the 90% threshold.

Statistics of EMT scores (76 GS), all cells

| | Day 0 means | Day 0 variances | Day 1 means | Day 1 variances | Day 2 means | Day 2 variances | Day 3 means | Day 3 variances | Day 4 means | Day 4 variances |
|---|---|---|---|---|---|---|---|---|---|---|
| **High EMT** | 0.726355 | 0.186205 | 0.635603 | 0.090942 | 0.970182 | 0.494843 | 0.584925 | 0.360403 | 0.434322 | 0.146789 |
| **Low EMT** | -1.072511 | 0.259053 | -0.856927 | 0.373864 | -1.538226 | 0.806734 | -1.245910 | 0.711708 | -0.621325 | 0.758056 |
| **Medium EMT** | 0.620413 | 0.122936 | 0.064054 | 0.268057 | -0.305083 | 0.889973 | -0.310389 | 0.682954 | 0.085325 | 0.423611 |
| **Undetermined** | 0.044529 | 0.368108 | 0.071171 | 0.367132 | 0.192809 | 1.075312 | 0.132190 | 0.559176 | 0.135254 | 0.469203 |

Statistics of EMT scores (76 GS), excluding outliers

| | Day 0 means | Day 0 variances | Day 1 means | Day 1 variances | Day 2 means | Day 2 variances | Day 3 means | Day 3 variances | Day 4 means | Day 4 variances |
|---|---|---|---|---|---|---|---|---|---|---|
| **High EMT** | 0.726355 | 0.186205 | 0.635603 | 0.090942 | 0.970182 | 0.494843 | 0.584925 | 0.360403 | 0.434322 | 0.146789 |
| **Low EMT** | -0.942329 | 0.159718 | -0.754350 | 0.258286 | -1.411997 | 0.577694 | -1.112071 | 0.494644 | -0.531103 | 0.519276 |
| **Medium EMT** | 0.620413 | 0.122936 | 0.064054 | 0.268057 | -0.305083 | 0.889973 | -0.292312 | 0.640357 | 0.098816 | 0.385136 |
| **Undetermined** | 0.050415 | 0.358362 | 0.078823 | 0.351082 | 0.205525 | 1.031855 | 0.132190 | 0.559176 | 0.135254 | 0.469203 |

*Table S5* Daily computed means and variances of EMT scores for the top ancestors of each EMT fate (day by day in each column), alongside the results for undetermined ancestors. The table compares the upper part (all cells) and the lower part (excluding outliers), where the upper part presents results for the entire top ancestor population and the lower part shows results after excluding the lower 1% of outliers from the distribution.

Stemness score, by ssGSEA and Lim et al. (75% top ancestors)

| | Day | Pair | Levene Statistic | Levene p-value | T-test Statistic | T-test p-value | Mean Group1 | Mean Group2 | Variance Group1 | Variance Group2 |
|---|---|---|---|---|---|---|---|---|---|---|
| **0** | 0.0 | Low EMT vs Medium EMT | 4.480206 | 0.035478 | -4.869787 | 0.000002 | -1.173722 | -0.750861 | 0.351928 | 0.236991 |
| **1** | 0.0 | Low EMT vs High EMT | 11.394061 | 0.000915 | 5.124832 | 0.000001 | -1.173722 | -1.88455 | 0.351928 | 0.097586 |
| **2** | 0.0 | Medium EMT vs High EMT | 4.212019 | 0.043582 | 9.516167 | 0.0 | -0.750861 | -1.88455 | 0.236991 | 0.097586 |
| **3** | 1.0 | Medium EMT vs High EMT | 1.246641 | 0.265077 | 16.141223 | 0.0 | -0.39769 | -1.427993 | 0.244227 | 0.363709 |
| **4** | 1.0 | Medium EMT vs Low EMT | 0.516415 | 0.47276 | 3.139963 | 0.001805 | -0.39769 | -0.550397 | 0.244227 | 0.265763 |
| **5** | 1.0 | High EMT vs Low EMT | 2.49855 | 0.114854 | -14.021236 | 0.0 | -1.427993 | -0.550397 | 0.363709 | 0.265763 |
| **6** | 2.0 | High EMT vs Low EMT | 0.279892 | 0.596956 | -10.993662 | 0.0 | -0.560861 | 0.035679 | 0.461974 | 0.479976 |
| **7** | 2.0 | High EMT vs Medium EMT | 25.698573 | 0.000001 | -23.307153 | 0.0 | -0.560861 | 0.535123 | 0.461974 | 0.202806 |
| **8** | 2.0 | Low EMT vs Medium EMT | 16.963365 | 0.000043 | -10.504516 | 0.0 | 0.035679 | 0.535123 | 0.479976 | 0.202806 |
| **9** | 3.0 | High EMT vs Medium EMT | 30.578043 | 0.0 | -19.162702 | 0.0 | -0.53576 | 0.300446 | 0.427005 | 0.234589 |
| **10** | 3.0 | High EMT vs Low EMT | 35.246037 | 0.0 | -14.823936 | 0.0 | -0.53576 | 0.105893 | 0.427005 | 0.237148 |
| **11** | 3.0 | Medium EMT vs Low EMT | 0.220665 | 0.63871 | 4.835717 | 0.000002 | 0.300446 | 0.105893 | 0.234589 | 0.237148 |
| **12** | 4.0 | High EMT vs Low EMT | 5.192336 | 0.02297 | -1.140509 | 0.254443 | 1.270952 | 1.294703 | 0.088881 | 0.070302 |
| **13** | 4.0 | High EMT vs Medium EMT | 0.003355 | 0.953832 | -14.145389 | 0.0 | 1.270952 | 1.627407 | 0.088881 | 0.092205 |
| **14** | 4.0 | Low EMT vs Medium EMT | 3.567005 | 0.05945 | -13.770133 | 0.0 | 1.294703 | 1.627407 | 0.070302 | 0.092205 |
| **15** | 8.0 | Medium EMT vs High EMT | 1.549171 | 0.213482 | 25.998243 | 0.0 | 1.696935 | 1.242699 | 0.07859 | 0.111146 |
| **16** | 8.0 | Medium EMT vs Low EMT | 6.119109 | 0.013517 | 19.365101 | 0.0 | 1.696935 | 1.388087 | 0.07859 | 0.067239 |
| **17** | 8.0 | High EMT vs Low EMT | 11.250238 | 0.000818 | -8.706501 | 0.0 | 1.242699 | 1.388087 | 0.111146 | 0.067239 |

Stemness score, by ssGSEA and Lim et al. (90% top ancestors)

| | Day | Pair | Levene Statistic | Levene p-value | T-test Statistic | T-test p-value | Mean Group1 | Mean Group2 | Variance Group1 | Variance Group2 |
|---|---|---|---|---|---|---|---|---|---|---|
| **0** | 0.0 | Low EMT vs Medium EMT | 1.334398 | 0.257454 | -2.621046 | 0.013812 | -1.798879 | -0.97894 | 0.188489 | 0.032111 |
| **1** | 0.0 | Low EMT vs High EMT | 1.647476 | 0.209824 | NaN | NaN | -1.798879 | -2.058722 | 0.188489 | NaN |
| **2** | 0.0 | Medium EMT vs High EMT | 1736749524563072991757521125376.0 | 0.0 | NaN | NaN | -0.97894 | -2.058722 | 0.032111 | NaN |
| **3** | 1.0 | Low EMT vs High EMT | 2.501511 | 0.118378 | 6.80059 | 0.0 | -0.856515 | -2.00057 | 0.284133 | 0.136282 |
| **4** | 1.0 | Low EMT vs Medium EMT | 2.986765 | 0.088009 | -1.857884 | 0.067059 | -0.856515 | -0.615266 | 0.284133 | 0.10762 |
| **5** | 1.0 | High EMT vs Medium EMT | 0.148454 | 0.702928 | -10.650776 | 0.0 | -2.00057 | -0.615266 | 0.136282 | 0.10762 |
| **6** | 2.0 | Low EMT vs Medium EMT | 7.424457 | 0.007042 | -4.539919 | 0.00001 | -0.254717 | 0.282267 | 0.59154 | 0.157905 |
| **7** | 2.0 | Low EMT vs High EMT | 0.851353 | 0.357517 | 8.386844 | 0.0 | -0.254717 | -1.661178 | 0.59154 | 0.338042 |
| **8** | 2.0 | Medium EMT vs High EMT | 2.810434 | 0.098314 | 16.334513 | 0.0 | 0.282267 | -1.661178 | 0.157905 | 0.338042 |
| **9** | 3.0 | Low EMT vs High EMT | 48.062777 | 0.0 | 10.78731 | 0.0 | 0.049557 | -0.815741 | 0.237105 | 0.759034 |
| **10** | 3.0 | Low EMT vs Medium EMT | 0.527751 | 0.468061 | -6.639987 | 0.0 | 0.049557 | 0.384561 | 0.237105 | 0.175939 |
| **11** | 3.0 | High EMT vs Medium EMT | 54.220471 | 0.0 | -14.322943 | 0.0 | -0.815741 | 0.384561 | 0.759034 | 0.175939 |
| **12** | 4.0 | High EMT vs Low EMT | 3.733445 | 0.053811 | 0.22279 | 0.823776 | 1.267052 | 1.261846 | 0.08766 | 0.069993 |
| **13** | 4.0 | High EMT vs Medium EMT | 2.247333 | 0.134454 | -15.663474 | 0.0 | 1.267052 | 1.676076 | 0.08766 | 0.072166 |
| **14** | 4.0 | Low EMT vs Medium EMT | 0.055005 | 0.814679 | -16.300591 | 0.0 | 1.261846 | 1.676076 | 0.069993 | 0.072166 |
| **15** | 8.0 | Medium EMT vs High EMT | 1.549171 | 0.213482 | 25.998243 | 0.0 | 1.696935 | 1.242699 | 0.07859 | 0.111146 |
| **16** | 8.0 | Medium EMT vs Low EMT | 6.119109 | 0.013517 | 19.365101 | 0.0 | 1.696935 | 1.388087 | 0.07859 | 0.067239 |
| **17** | 8.0 | High EMT vs Low EMT | 11.250238 | 0.000818 | -8.706501 | 0.0 | 1.242699 | 1.388087 | 0.111146 | 0.067239 |

*Table S6* Statistics of the stemness signature score (derived from Lim et al. gene set) were calculated for the top ancestors across the three EMT trajectories for all time points, spanning from day 0 to day 8. The difference of variations among the trajectories were computed using the Levene test, while the difference of means were analyzed using the t-test. Additionally, the table displays the mean and variation for the top ancestors of each trajectory group on every day. In the context of these values, 'Group 1' corresponds to the first trajectory listed in the "EMT pair" column, while 'Group 2' signifies the second. Upper table: results determined by the 75% threshold for identifying the top ancestor set. Lower table: results based on the 90% threshold.

Proliferation score, by ssGSEA and Ben-Porath et al. (75% top ancestors)

| | Day | Pair | Levene Statistic | Levene p-value | T-test Statistic | T-test p-value | Mean Group1 | Mean Group2 | Variance Group1 | Variance Group2 |
|---|---|---|---|---|---|---|---|---|---|---|
| **0** | 0.0 | Low EMT vs Medium EMT | 10.160539 | 0.001656 | 2.700274 | 0.007499 | -0.369399 | -0.66106 | 0.586137 | 0.259085 |
| **1** | 0.0 | Low EMT vs High EMT | 5.028088 | 0.026248 | 10.02371 | 0.0 | -0.369399 | -2.169226 | 0.586137 | 0.193882 |
| **2** | 0.0 | Medium EMT vs High EMT | 0.17744 | 0.674772 | 11.583194 | 0.0 | -0.66106 | -2.169226 | 0.259085 | 0.193882 |
| **3** | 1.0 | Medium EMT vs High EMT | 94.678447 | 0.0 | 2.904459 | 0.003949 | -1.110924 | -1.363735 | 0.263853 | 1.016774 |
| **4** | 1.0 | Medium EMT vs Low EMT | 9.052907 | 0.002775 | -3.67977 | 0.000263 | -1.110924 | -0.902536 | 0.263853 | 0.414814 |
| **5** | 1.0 | High EMT vs Low EMT | 51.904491 | 0.0 | -5.167992 | 0.0 | -1.363735 | -0.902536 | 1.016774 | 0.414814 |
| **6** | 2.0 | High EMT vs Low EMT | 1.655559 | 0.198671 | 0.678106 | 0.49795 | -0.017461 | -0.064258 | 0.88204 | 0.644092 |
| **7** | 2.0 | High EMT vs Medium EMT | 28.875157 | 0.0 | 3.539344 | 0.000432 | -0.017461 | -0.236539 | 0.88204 | 0.263422 |
| **8** | 2.0 | Low EMT vs Medium EMT | 22.549507 | 0.000003 | 3.141999 | 0.001759 | -0.064258 | -0.236539 | 0.644092 | 0.263422 |
| **9** | 3.0 | High EMT vs Medium EMT | 25.801959 | 0.0 | 5.343928 | 0.0 | -0.206644 | -0.496502 | 0.686733 | 0.310146 |
| **10** | 3.0 | High EMT vs Low EMT | 2.354009 | 0.125333 | 0.49777 | 0.618775 | -0.206644 | -0.234804 | 0.686733 | 0.485788 |
| **11** | 3.0 | Medium EMT vs Low EMT | 16.534443 | 0.000054 | -5.001197 | 0.000001 | -0.496502 | -0.234804 | 0.310146 | 0.485788 |
| **12** | 4.0 | High EMT vs Low EMT | 9.695978 | 0.001918 | 3.173086 | 0.001571 | 1.386745 | 1.22707 | 0.66463 | 0.244212 |
| **13** | 4.0 | High EMT vs Medium EMT | 5.562146 | 0.018665 | 8.840504 | 0.0 | 1.386745 | 0.852516 | 0.66463 | 0.254645 |
| **14** | 4.0 | Low EMT vs Medium EMT | 0.128979 | 0.719627 | 8.734583 | 0.0 | 1.22707 | 0.852516 | 0.244212 | 0.254645 |
| **15** | 8.0 | Medium EMT vs High EMT | 9.622776 | 0.001963 | -16.116834 | 0.0 | 0.786232 | 1.304383 | 0.2447 | 0.392277 |
| **16** | 8.0 | Medium EMT vs Low EMT | 0.004536 | 0.946316 | -13.969659 | 0.0 | 0.786232 | 1.186944 | 0.2447 | 0.226485 |
| **17** | 8.0 | High EMT vs Low EMT | 10.911511 | 0.000981 | 3.771444 | 0.000169 | 1.304383 | 1.186944 | 0.392277 | 0.226485 |

Proliferation score, by ssGSEA and Ben-Porath et al. (90% top ancestors)

| | Day | Pair | Levene Statistic | Levene p-value | T-test Statistic | T-test p-value | Mean Group1 | Mean Group2 | Variance Group1 | Variance Group2 |
|---|---|---|---|---|---|---|---|---|---|---|
| **0** | 0.0 | Low EMT vs Medium EMT | 2.148121 | 0.153504 | 0.364115 | 0.718413 | -0.99409 | -1.174839 | 0.477468 | 0.001096 |
| **1** | 0.0 | Low EMT vs High EMT | 1.177809 | 0.287058 | NaN | NaN | -0.99409 | -2.79267 | 0.477468 | NaN |
| **2** | 0.0 | Medium EMT vs High EMT | 14820959572726510952197914624.0 | 0.0 | NaN | NaN | -1.174839 | -2.79267 | 0.001096 | NaN |
| **3** | 1.0 | Low EMT vs High EMT | 4.24152 | 0.043278 | 6.425908 | 0.0 | -1.078742 | -2.483922 | 0.502657 | 0.099334 |
| **4** | 1.0 | Low EMT vs Medium EMT | 1.692432 | 0.197211 | 0.01849 | 0.985297 | -1.078742 | -1.081984 | 0.502657 | 0.245616 |
| **5** | 1.0 | High EMT vs Medium EMT | 1.916847 | 0.17714 | -8.414803 | 0.0 | -2.483922 | -1.081984 | 0.099334 | 0.245616 |
| **6** | 2.0 | Low EMT vs Medium EMT | 7.75971 | 0.00589 | -0.47781 | 0.633341 | -0.179341 | -0.107777 | 0.940721 | 0.27794 |
| **7** | 2.0 | Low EMT vs High EMT | 0.937215 | 0.334412 | 9.452783 | 0.0 | -0.179341 | -2.204963 | 0.940721 | 0.715497 |
| **8** | 2.0 | Medium EMT vs High EMT | 1.681256 | 0.199206 | 12.647273 | 0.0 | -0.107777 | -2.204963 | 0.27794 | 0.715497 |
| **9** | 3.0 | Low EMT vs High EMT | 68.600724 | 0.0 | 6.141634 | 0.0 | -0.152743 | -0.849467 | 0.486715 | 1.493162 |
| **10** | 3.0 | Low EMT vs Medium EMT | 14.871498 | 0.000138 | 3.772802 | 0.000191 | -0.152743 | -0.405734 | 0.486715 | 0.221874 |
| **11** | 3.0 | High EMT vs Medium EMT | 140.776436 | 0.0 | -3.965786 | 0.000097 | -0.849467 | -0.405734 | 1.493162 | 0.221874 |
| **12** | 4.0 | High EMT vs Low EMT | 9.835502 | 0.001797 | 2.539527 | 0.011355 | 1.395125 | 1.240573 | 0.766595 | 0.257235 |
| **13** | 4.0 | High EMT vs Medium EMT | 10.258926 | 0.001444 | 8.283164 | 0.0 | 1.395125 | 0.828002 | 0.766595 | 0.208271 |
| **14** | 4.0 | Low EMT vs Medium EMT | 0.586072 | 0.444344 | 8.885102 | 0.0 | 1.240573 | 0.828002 | 0.257235 | 0.208271 |
| **15** | 8.0 | Medium EMT vs High EMT | 9.622776 | 0.001963 | -16.116834 | 0.0 | 0.786232 | 1.304383 | 0.2447 | 0.392277 |
| **16** | 8.0 | Medium EMT vs Low EMT | 0.004536 | 0.946316 | -13.969659 | 0.0 | 0.786232 | 1.186944 | 0.2447 | 0.226485 |
| **17** | 8.0 | High EMT vs Low EMT | 10.911511 | 0.000981 | 3.771444 | 0.000169 | 1.304383 | 1.186944 | 0.392277 | 0.226485 |

*Table S7* Statistics of the proliferation signature score (derived from Ben-Porath et al gene set) were calculated for the top ancestors across the three EMT trajectories for all time points, spanning from day 0 to day 8. The difference of variations among the trajectories were computed using the Levene test, while the difference of means were analyzed using the t-test. Additionally, the table displays the mean and variation for the top ancestors of each trajectory group on every day. In the context of these values, 'Group 1' corresponds to the first trajectory listed in the "EMT pair" column, while 'Group 2' signifies the second. Upper table: results determined by the 75% threshold for identifying the top ancestor set. Lower table: results based on the 90% threshold.

EMT score (Group 1) and Stemness (Group 2) - 75% threshold

| Day | | means_1 | variances_1 | means_2 | variances_2 | correlations |
|---|---|---|---|---|---|---|
| Day 0 | High EMT | 0.726355 | 0.186205 | -1.884550 | 0.097586 | 0.025134 |
| | Low EMT | -1.072511 | 0.259053 | -1.173722 | 0.351928 | 0.421023 |
| | Medium EMT | 0.620413 | 0.122936 | -0.750861 | 0.236991 | -0.420864 |
| | Undetermined | 0.044529 | 0.368108 | -0.805632 | 0.215333 | -0.410723 |
| Day 1 | High EMT | 0.635603 | 0.090942 | -1.427993 | 0.363709 | -0.358138 |
| | Low EMT | -0.856927 | 0.373864 | -0.550397 | 0.265763 | 0.278544 |
| | Medium EMT | 0.064054 | 0.268057 | -0.397690 | 0.244227 | -0.592494 |
| | Undetermined | 0.071171 | 0.367132 | -0.685929 | 0.298768 | -0.530534 |
| Day 2 | High EMT | 0.970182 | 0.494843 | -0.560861 | 0.461974 | -0.452733 |
| | Low EMT | -1.538226 | 0.806734 | 0.035679 | 0.479976 | 0.324676 |
| | Medium EMT | -0.305083 | 0.889973 | 0.535123 | 0.202806 | -0.599306 |
| | Undetermined | 0.192809 | 1.075312 | -0.022364 | 0.323694 | -0.535845 |

| Day | | means_1 | variances_1 | means_2 | variances_2 | correlations |
|---|---|---|---|---|---|---|
| Day 3 | High EMT | 0.584925 | 0.360403 | -0.535760 | 0.427005 | -0.604038 |
| | Low EMT | -1.245910 | 0.711708 | 0.105893 | 0.237148 | 0.215084 |
| | Medium EMT | -0.310389 | 0.682954 | 0.300446 | 0.234589 | -0.521833 |
| | Undetermined | 0.132190 | 0.559176 | -0.181665 | 0.315992 | -0.578274 |
| Day 4 | High EMT | 0.434322 | 0.146789 | 1.270952 | 0.088881 | -0.104017 |
| | Low EMT | -0.621325 | 0.758056 | 1.294703 | 0.070302 | 0.442763 |
| | Medium EMT | 0.085325 | 0.423611 | 1.627407 | 0.092205 | -0.339982 |
| | Undetermined | 0.135254 | 0.469203 | 1.362257 | 0.075350 | -0.075643 |
| Day 8 | High EMT | 0.323103 | 0.202847 | 1.242699 | 0.111146 | -0.296016 |
| | Low EMT | -0.624904 | 0.812004 | 1.388087 | 0.067239 | 0.359263 |
| | Medium EMT | 0.228478 | 0.432259 | 1.696935 | 0.078590 | -0.260133 |

EMT score (Group 1) and Stemness (Group 2) - 90% threshold

| Day | | means_1 | variances_1 | means_2 | variances_2 | correlations |
|---|---|---|---|---|---|---|
| Day 0 | High EMT | -0.126174 | NaN | -2.058722 | NaN | NaN |
| | Low EMT | -1.631174 | 0.196975 | -1.798879 | 0.188489 | -0.080882 |
| | Medium EMT | 0.373108 | 0.001447 | -0.978940 | 0.032111 | 1.000000 |
| | Undetermined | 0.017278 | 0.406774 | -0.821341 | 0.227332 | -0.333245 |
| Day 1 | High EMT | 0.602346 | 0.082510 | -2.000570 | 0.136282 | 0.113166 |
| | Low EMT | -1.485779 | 0.280106 | -0.856515 | 0.284133 | 0.240833 |
| | Medium EMT | 0.526225 | 0.022929 | -0.615266 | 0.107620 | -0.393298 |
| | Undetermined | 0.032085 | 0.392596 | -0.671995 | 0.323374 | -0.522139 |
| Day 2 | High EMT | 0.679536 | 0.782281 | -1.661178 | 0.338042 | 0.077873 |
| | Low EMT | -1.981631 | 0.795766 | -0.254717 | 0.591540 | 0.339670 |
| | Medium EMT | 0.708945 | 0.170549 | 0.282267 | 0.157905 | -0.526763 |
| | Undetermined | 0.109370 | 1.205251 | 0.009501 | 0.386366 | -0.587001 |

| Day | | means_1 | variances_1 | means_2 | variances_2 | correlations |
|---|---|---|---|---|---|---|
| Day 3 | High EMT | 0.462706 | 0.408855 | -0.815741 | 0.759034 | -0.602853 |
| | Low EMT | -1.414263 | 0.791107 | 0.049557 | 0.237105 | 0.294257 |
| | Medium EMT | -0.172029 | 0.548028 | 0.384561 | 0.175939 | -0.587650 |
| | Undetermined | 0.147628 | 0.671855 | -0.204923 | 0.358997 | -0.630680 |
| Day 4 | High EMT | 0.449949 | 0.135711 | 1.267052 | 0.087660 | -0.065054 |
| | Low EMT | -0.771463 | 0.786140 | 1.261846 | 0.069993 | 0.458174 |
| | Medium EMT | 0.052257 | 0.406247 | 1.676076 | 0.072166 | -0.244681 |
| | Undetermined | 0.118668 | 0.432779 | 1.355432 | 0.079172 | -0.142342 |
| Day 8 | High EMT | 0.323103 | 0.202847 | 1.242699 | 0.111146 | -0.296016 |
| | Low EMT | -0.624904 | 0.812004 | 1.388087 | 0.067239 | 0.359263 |
| | Medium EMT | 0.228478 | 0.432259 | 1.696935 | 0.078590 | -0.260133 |

*Table S8*. Daily calculations of means and variances for EMT scores (group 1) and stemness scores (group 2) of the top ancestors for each EMT fate, in addition to the results for undetermined ancestors. The table also includes computed Pearson correlations between these two scores. Upper table: results determined by the 75% threshold for identifying the top ancestor set. Lower table: results based on the 90% threshold.

EED gene

| | Day | Pair | Levene Statistic | Levene p-value | T-test Statistic | T-test p-value | Mean Group1 | Mean Group2 | Variance Group1 | Variance Group2 |
|---|---|---|---|---|---|---|---|---|---|---|
| **0** | 0.0 | Low EMT vs Medium EMT | 4.279729 | 0.039806 | 2.068751 | 0.039806 | -0.004529 | -0.296355 | 0.984109 | 0.48234 |
| **1** | 0.0 | Low EMT vs High EMT | 1.042995 | 0.308594 | 1.021271 | 0.308594 | -0.004529 | -0.259568 | 0.984109 | 1.622216 |
| **2** | 0.0 | Medium EMT vs High EMT | 0.025852 | 0.872688 | -0.160787 | 0.872688 | -0.296355 | -0.259568 | 0.48234 | 1.622216 |
| **3** | 1.0 | Medium EMT vs High EMT | 8.6674 | 0.003489 | -2.944045 | 0.003489 | -0.422357 | -0.085868 | 0.488633 | 1.692847 |
| **4** | 1.0 | Medium EMT vs Low EMT | 4.249956 | 0.039844 | -2.061542 | 0.039844 | -0.422357 | -0.249972 | 0.488633 | 0.974513 |
| **5** | 1.0 | High EMT vs Low EMT | 1.699669 | 0.193188 | 1.303714 | 0.193188 | -0.085868 | -0.249972 | 1.692847 | 0.974513 |
| **6** | 2.0 | High EMT vs Low EMT | 24.248766 | 0.000001 | 4.924304 | 0.000001 | 0.260665 | -0.102291 | 1.159829 | 0.583911 |
| **7** | 2.0 | High EMT vs Medium EMT | 53.393045 | 0.0 | 7.307054 | 0.0 | 0.260665 | -0.270579 | 1.159829 | 0.424912 |
| **8** | 2.0 | Low EMT vs Medium EMT | 8.577675 | 0.003529 | 2.928767 | 0.003529 | -0.102291 | -0.270579 | 0.583911 | 0.424912 |
| **9** | 3.0 | High EMT vs Medium EMT | 18.319546 | 0.000021 | 4.280134 | 0.000021 | 0.177827 | -0.162689 | 1.349913 | 0.914207 |
| **10** | 3.0 | High EMT vs Low EMT | 3.941752 | 0.047423 | 1.985385 | 0.047423 | 0.177827 | 0.020201 | 1.349913 | 0.961676 |
| **11** | 3.0 | Medium EMT vs Low EMT | 5.195419 | 0.023008 | -2.279346 | 0.023008 | -0.162689 | 0.020201 | 0.914207 | 0.961676 |
| **12** | 4.0 | High EMT vs Low EMT | 0.214094 | 0.643713 | -1.161323 | 0.245885 | 0.105966 | 0.159854 | 0.453319 | 0.3353 |
| **13** | 4.0 | High EMT vs Medium EMT | 5.269866 | 0.022033 | 3.487884 | 0.000522 | 0.105966 | -0.07716 | 0.453319 | 0.279116 |
| **14** | 4.0 | Low EMT vs Medium EMT | 5.163573 | 0.02344 | 4.918908 | 0.000001 | 0.159854 | -0.07716 | 0.3353 | 0.279116 |
| **15** | 8.0 | Medium EMT vs High EMT | 13.833237 | 0.000208 | -7.180864 | 0.0 | -0.168616 | 0.104036 | 0.372398 | 0.523983 |
| **16** | 8.0 | Medium EMT vs Low EMT | 8.572875 | 0.00348 | -6.631898 | 0.0 | -0.168616 | 0.08026 | 0.372398 | 0.431467 |
| **17** | 8.0 | High EMT vs Low EMT | 0.810855 | 0.36803 | 0.62055 | 0.535002 | 0.104036 | 0.08026 | 0.523983 | 0.431467 |

*Table S9* Statistics of the EED gene expression levels were calculated for the top ancestors across the three EMT trajectories for all time points, spanning from day 0 to day 8. The difference of variations among the trajectories were computed using the Levene test, while the difference of means were analyzed using the t-test. Additionally, the table displays the mean and variation for the top ancestors of each trajectory group on every day. In the context of these values, 'Group 1' corresponds to the first trajectory listed in the "EMT pair" column, while 'Group 2' signifies the second.

EZH2 gene

| | Day | Pair | Levene Statistic | Levene p-value | T-test Statistic | T-test p-value | Mean Group1 | Mean Group2 | Variance Group1 | Variance Group2 |
|---|---|---|---|---|---|---|---|---|---|---|
| **0** | 0.0 | Low EMT vs Medium EMT | 14.537998 | 0.000181 | 3.812873 | 0.000181 | 0.156532 | -0.431688 | 1.238986 | 0.416651 |
| **1** | 0.0 | Low EMT vs High EMT | 0.001025 | 0.974495 | 0.032018 | 0.974495 | 0.156532 | 0.146939 | 1.238986 | 3.814014 |
| **2** | 0.0 | Medium EMT vs High EMT | 3.939938 | 0.050761 | -1.984928 | 0.050761 | -0.431688 | 0.146939 | 0.416651 | 3.814014 |
| **3** | 1.0 | Medium EMT vs High EMT | 19.845047 | 0.000012 | -4.454778 | 0.000012 | -0.536948 | -0.024079 | 0.378876 | 1.926753 |
| **4** | 1.0 | Medium EMT vs Low EMT | 19.645314 | 0.000012 | -4.432303 | 0.000012 | -0.536948 | -0.152225 | 0.378876 | 1.169502 |
| **5** | 1.0 | High EMT vs Low EMT | 0.883673 | 0.347846 | 0.940039 | 0.347846 | -0.024079 | -0.152225 | 1.926753 | 1.169502 |
| **6** | 2.0 | High EMT vs Low EMT | 5.564139 | 0.018633 | 1.513378 | 0.130679 | 0.217301 | 0.094531 | 1.247735 | 0.861617 |
| **7** | 2.0 | High EMT vs Medium EMT | 177.660833 | 0.0 | 9.574365 | 0.0 | 0.217301 | -0.458175 | 1.247735 | 0.23235 |
| **8** | 2.0 | Low EMT vs Medium EMT | 152.641426 | 0.0 | 9.148047 | 0.0 | 0.094531 | -0.458175 | 0.861617 | 0.23235 |
| **9** | 3.0 | High EMT vs Medium EMT | 75.864165 | 0.0 | 8.710004 | 0.0 | 0.233341 | -0.422613 | 1.416943 | 0.417595 |
| **10** | 3.0 | High EMT vs Low EMT | 8.126074 | 0.004469 | 2.850627 | 0.004469 | 0.233341 | 0.00489 | 1.416943 | 0.901403 |
| **11** | 3.0 | Medium EMT vs Low EMT | 40.206435 | 0.0 | -6.340854 | 0.0 | -0.422613 | 0.00489 | 0.417595 | 0.901403 |
| **12** | 4.0 | High EMT vs Low EMT | 0.765258 | 0.381971 | -0.779161 | 0.436134 | 0.138654 | 0.175288 | 0.407458 | 0.410432 |
| **13** | 4.0 | High EMT vs Medium EMT | 26.712793 | 0.0 | 9.420052 | 0.0 | 0.138654 | -0.335715 | 0.407458 | 0.273922 |
| **14** | 4.0 | Low EMT vs Medium EMT | 35.732165 | 0.0 | 9.932308 | 0.0 | 0.175288 | -0.335715 | 0.410432 | 0.273922 |
| **15** | 8.0 | Medium EMT vs High EMT | 85.996756 | 0.0 | -15.414196 | 0.0 | -0.420054 | 0.105603 | 0.198793 | 0.498383 |
| **16** | 8.0 | Medium EMT vs Low EMT | 130.107131 | 0.0 | -15.548729 | 0.0 | -0.420054 | 0.117364 | 0.198793 | 0.476549 |
| **17** | 8.0 | High EMT vs Low EMT | 2.049167 | 0.152524 | -0.305218 | 0.760248 | 0.105603 | 0.117364 | 0.498383 | 0.476549 |

*Table S10* Statistics of the EZH2 gene expression levels were calculated for the top ancestors across the three EMT trajectories for all time points, spanning from day 0 to day 8. The difference of variations among the trajectories were computed using the Levene test, while the difference of means were analyzed using the t-test. Additionally, the table displays the mean and variation for the top ancestors of each trajectory group on every day. In the context of these values, 'Group 1' corresponds to the first trajectory listed in the "EMT pair" column, while 'Group 2' signifies the second.

A

Single gene expression (our prediction for Deshmusk et al)
partial EMT compared to full EMT Day 8

Up-regulation

| | P-value | Fold change |
|---|---|---|
| **TGFBI** | 7.2151e -148 | 1.4812 |
| **POSTN** | 6.0077e -44 | 1.3372 |
| **KRT8** | 4.4580e -95 | 1.3156 |
| **DSP** | 8.6890e -23 | 1.1175 |
| **SNAI1** | 1.6754e -90 | 1.0828 |
| **CDH2** | 1.8064e -16 | 1.0539 |
| **TWIST1** | 4.7397e -24 | 1.0538 |
| **CDH1** | 1.5549e -11 | 1.0269 |
| **JARID2** | 1.5197e -02 | 1.0141 |
| **ZEB2** | 1.5063e -02 | 1.0084 |
| **EPCAM** | 2.7944e -17 | 0.9536 |
| **KMT2A** | 1.6705e -21 | 0.9391 |
| **EED** | 1.7288e -31 | 0.9388 |
| **SUZ12** | 8.5457e -103 | 0.8727 |
| **EZH2** | 5.2996e -147 | 0.8584 |
| **PHF19** | 3.8191e -208 | 0.6611 |

Down-regulation

B

Single gene expression (transcriptomics from Zhang et al)
EED-KO mesenchymal cells compared to KMT2D-KO mesenchymal cells

Up-regulation

| | P-value | Fold change |
|---|---|---|
| **KRT8** | 1.3332e -56 | 1.9414 |
| **CDH2** | 5.6672e -30 | 1.1722 |
| **POSTN** | 8.8751e -15 | 1.3004 |
| **ZEB1** | 1.2055e- 14 | 1.1693 |
| **TGFBI** | 6.8129e -08 | 1.2387 |
| **EZH2** | 7.6606e -08 | 1.1119 |
| **PHF19** | 2.9178e -07 | 1.1612 |
| **CDH1** | 5.2352e -02 | 0.9842 |
| **ZEB2** | 4.0405e -04 | 0.9399 |
| **SUZ12** | 1.1552e -01 | 0.9758 |
| **JARID2** | 4.0498e -06 | 0.9306 |
| **MTF2** | 1.4611e -06 | 0.9218 |
| **KMT2A** | 4.3251e -09 | 0.8935 |
| **SNAI1** | 5.9715e- 13 | 0.9320 |
| **EED** | 2.7530e -24 | 0.8884 |
| **TWIST1** | 3.9236e -35 | 0.7676 |

Down-regulation

*Table S11* Two tables displaying the differential expression of the curated EMT-related genes for two separate datasets. (A) Left table illustrates the DEGs at day 8 between the partial EMT and high EMT trajectories, based on the dataset from Deshmukh et al. (B) Right table presents the DEGs in mesenchymal cells with EED knock-out (C1-sgEEG-Mes) compared to KMT2D knock-out cells (C1-sgKMT2D-Mes), derived from the CRISPR study by Zhang et al. The p-value is calculated using a t-test comparing the mean expressions of two populations, and the fold change is the ratio of the former to the latter, i.e., partial EMT to full EMT in (A), and C1-sgEEG-Mes to C1-sgKMT2D-Mes in (B).

### References


1. Villani, C. *Optimal Transport: Old and New*. (Springer Berlin Heidelberg, 2016).
2. Chizat, L., Peyré, G., Schmitzer, B. & Vialard, F.-X. Scaling algorithms for unbalanced optimal transport problems. *Math. Comput.* **87**, 2563–2609 (2018).
3. Schiebinger, G. *et al.* Optimal-Transport Analysis of Single-Cell Gene Expression Identifies Developmental Trajectories in Reprogramming. *Cell* **176**, 928-943.e22 (2019).
